# Genome sequence of the cluster root forming white lupin

**DOI:** 10.1101/708917

**Authors:** Bárbara Hufnagel, André Marques, Alexandre Soriano, Laurence Marquès, Fanchon Divol, Patrick Doumas, Erika Sallet, Davide Mancinotti, Sébastien Carrere, William Marande, Sandrine Arribat, Jean Keller, Cécile Huneau, Thomas Blein, Delphine Aime, Malika Laguerre, Jemma Taylor, Veit Schubert, Matthew Nelson, Fernando Geu-Flores, Martin Crespi, Karine Gallardo-Guerrero, Pierre-Marc Delaux, Jérôme Salse, Hélène Bergès, Romain Guyot, Jérôme Gouzy, Benjamin Péret

## Abstract

White lupin (*Lupinus albus L.*) is a legume that produces seeds recognized for their high protein content and good nutritional value (lowest glycemic index of all grains, high dietary fiber content, and zero gluten or starch)^1–5^. White lupin can form nitrogen-fixing nodules but has lost the ability to form mycorrhizal symbiosis with fungi^6^. Nevertheless, its root system is well adapted to poor soils: it produces cluster roots, constituted of dozens of determinate lateral roots that improve soil exploration and phosphate remobilization^7^. As phosphate is a limited resource that comes from rock reserves^8^, the production of cluster roots is a trait of interest to improve fertilizers efficiency. Using long reads sequencing technologies, we provide a high-quality genome sequence of a modern variety of white lupin (2n=50, 451 Mb), as well as *de novo* assemblies of a landrace and a wild relative. We describe how domestication impacted soil exploration capacity through the early establishment of lateral and cluster roots. We identify the *APETALA2* transcription factor *LaPUCHI-1*, homolog of the Arabidopsis morphogenesis coordinator^9^, as a potential regulator of this trait. Our high-quality genome and companion genomic and transcriptomic resources enable the development of modern breeding strategies to increase and stabilize yield and to develop new varieties with reduced allergenic properties (caused by conglutins^10^), which would favor the deployment of this promising culture.

---

We generated 164x sequencing coverage of the genome of *Lupinus albus* cv. AMIGA using 30 single-molecule real-time (SMRT) cells on PacBio Sequel platform as well as 208x (119 Gb) of Illumina paired-end sequences for the assembly polishing. The contig sequences obtained by a meta assembly strategy based on CANU and FALCON^11^ (Supplementary Note 1) were scaffolded in a first step using a Bionano optical map and in a second step using a high density genetic map^12^. The chromosome-level assembly (termed Lalb, Extended Data Table 1, Extended Data Fig. 1) covers the 25 nuclear chromosomes along with chloroplast genome, leaving 64 unanchored contigs (8.8Mb – 2% of the assembly). The maximum number of sequence gaps is four (chromosomes 10 and 11) and ten chromosomes contain a single sequence gap illustrating the high and homogenous contiguity across chromosomes.

We generated RNA-seq data from ten different organs widely covering gene expression in WL (Supplementary Note 1). The assembled reads were integrated using EuGene-EP pipeline^13^. Genome annotation identified 38,258 protein-coding genes and 3,129 non-protein-coding genes (Extended Data Table 1 and Supplementary Note 1). Evidence of transcription was found for 92% of the annotated genes. Quality of the annotation was evaluated with a Benchmarking of Universal Single-Copy Orthologs (BUSCO^15^) analysis, yielding a completeness score of 97.7%. The WL Genome portal (www.whitelupin.fr) gathers a genome browser and several other user-friendly tools for molecular analysis.

*De novo* identification of repeated elements revealed a highly repetitive genome (60%), with over 75% repeats matching known transposable elements (TEs, Fig. 1a). TEs were most commonly long terminal repeats (LTRs) retrotransposons (34%), with remarkable accumulation of Ty3/gypsy Tekay, CRM chromoviruses and Ty1/copia SIRE towards the central regions of chromosome assemblies along presumed (peri)centromeric regions (Fig. 1b). Class II TEs accounted for *ca.* 0.8% of the genome (Supplementary Note 2) and is in accordance to the lower abundance of this class of repeats in other legume species^14,15^. A high amount of satellite DNA (satDNA) sequences was found, comprising ∼15% of the genome (Supplementary Note 2). A high association of CRM clade retroelements here referred to as Centromeric Retrotransposon of White Lupin (CRWL) and satDNA peaks was observed, suggesting a more centromere-specific distribution of these repeats (Fig. 1b-c) compared to other TEs whereas a wider distribution towards the pericentromeres of WL chromosomes was observed (Extended Data Fig. 2).

**Figure 1.**
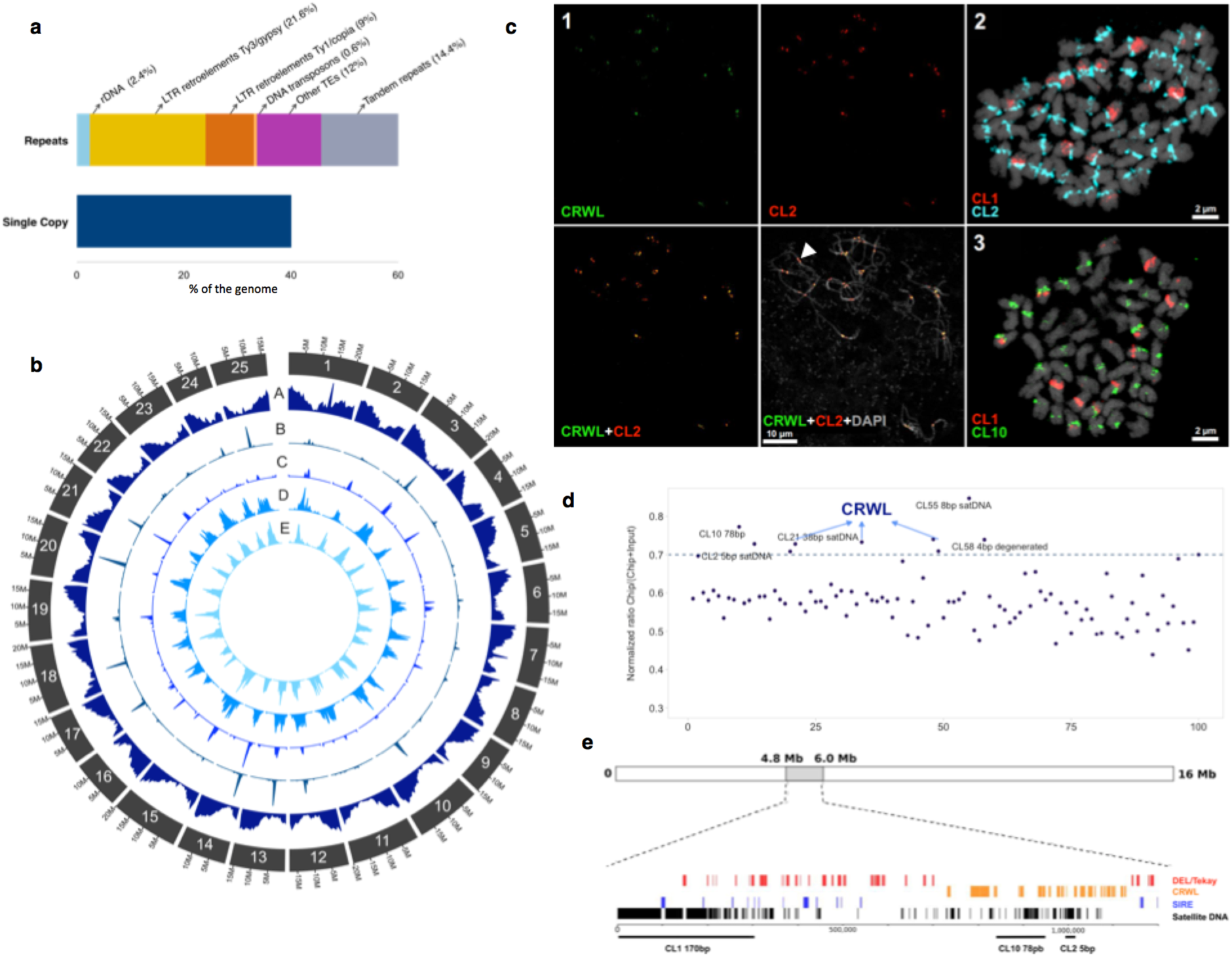
Overall characterization of repeated elements abundance in white lupin genome. **(a)** Proportion of single copy and repeated sequences for the different classes of repeats. **(b)** Density distribution along the chromosomes of the repetitive classes A. genes, B. CRM, C. satDNA, D. Tekay and E. SIRE. Density is represented in 0.5 Mb bins. **(c)** FISH mapping of the main repeats by super-resolution microscopy (3D-SIM). (1) Co-localization of CRWL and CL2 satDNA on centromeric regions of meiotic (pachytene) chromosomes. (2-3) Distribution of the most abundant satellite DNAs CL1, CL2 and CL10 in somatic metaphase chromosomes. **(d)** LalbCENH3-ChIPseq reads mapped against the first 100 RepeatExplorer clusters of the WL genome. The main centromeric sequences found in LalbCENH3-ChIPseq are highlighted. (**e**) Typical centromere composition of a WL chromosome, chromomosome 14.

Raising an anti-LalbCENH3 antibody, we mapped functional centromeres using immunostaining (Supplementary Fig. 1) and performed LalbCENH3-ChIPseq confirming the association of CRWL main clusters (Fig. 1d) with functional centromeres. These regions are also enriched with four families of centromeric satDNA: CL2-5bp, CL10-78bp, CL21-38bp and CL55-8bp (Fig. 1d). The total amount of cenDNA represents about 11% (49.55 Mb) of the genome. In contrast, the most abundant satDNA CL1-170bp did not show significant enrichment with the immunoprecipitated DNA, suggesting that this element is excluded from functional centromeres. A typical (peri)centromeric region of a WL chromosome contains the most abundant CL1-170 bp repeats representing 18% of the region. These sequences are organized in blocks separated by SIRE retrotransposons. Centromere-associated satellite repeats are present in shorter arrays such as CL2-5bp and CL10-78bp intermingled with CRWL elements (Fig. 1e, Supplementary Fig. 1). These results identify a specific centromeric sequence pattern with a highly diverse structure in WL that strongly differs from known centromeric sequences.

To provide a first overview of WL diversity and domestication patterns we sequenced 14 WL accessions, including 11 modern varieties, 1 landrace and 2 “wild” relatives (Supplementary Note 3). The accessions presented a total of 4,643,439 SNPs (Fig. 2a) and were divided into three clades based on pairwise dissimilarity and principal component analysis, reflecting their recent breeding history: winter varieties (vernalization responsive, slow growth, cold adapted), spring varieties (vernalization unresponsive, fast growth, strong vigor, reduced life-cycle) and landraces/wild types (Fig. 2b, Supplementary Note 3).

**Figure 2:**
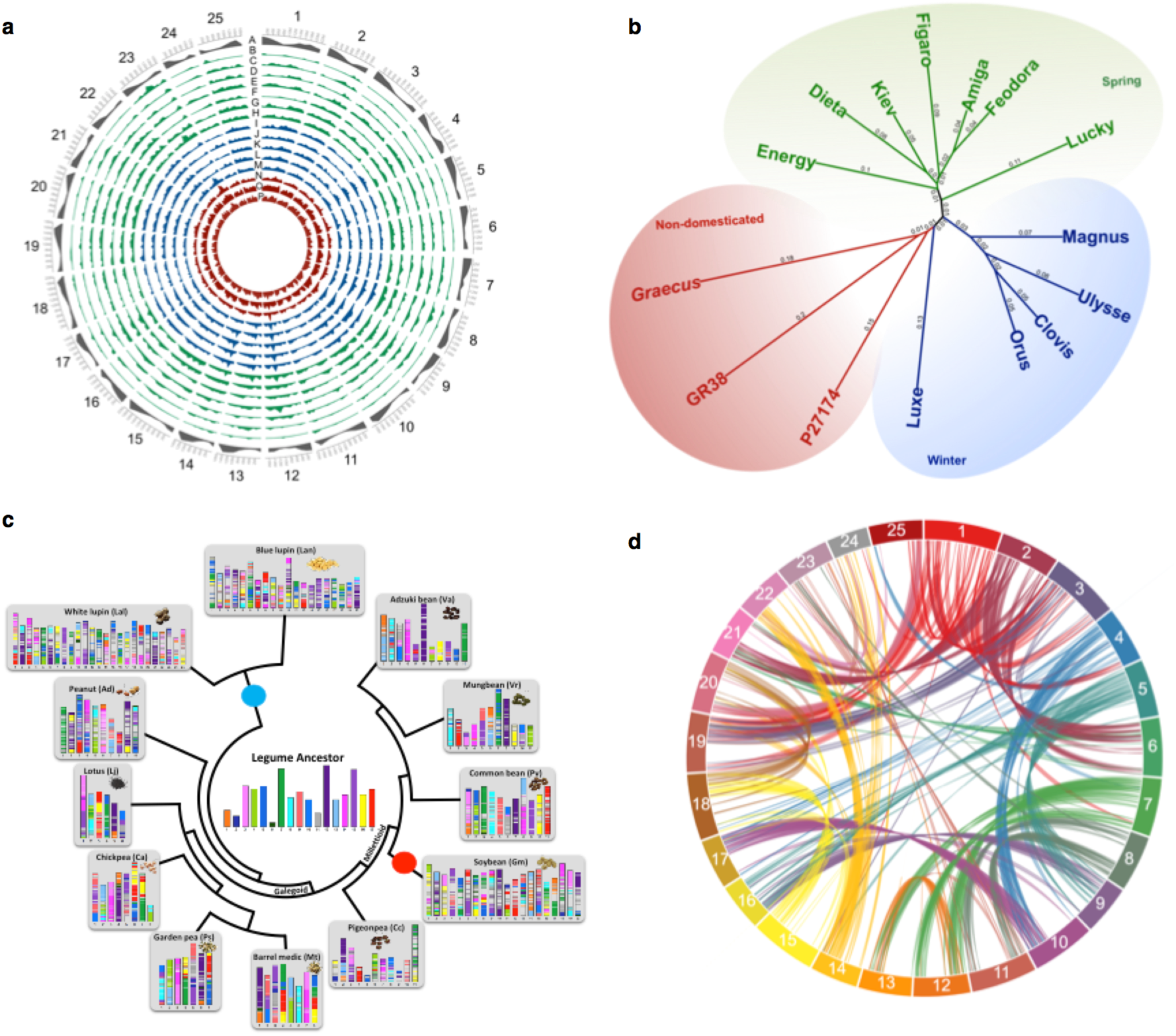
White Lupin diversity and evolution history. **(a)** SNP density identified by resequencing of 15 varieties of white lupin. In the outer track (A in grey) gene density is shown. Spring varieties are represented in green (B-H), winter varieties are represented in blue (I-M) and non-domesticated/landrace varieties are represented in red (N-P). From the outer to inner track: (B) AMIGA, (C) FEODORA, (D) KIEV, (E) DIETA, (F) FIGARO, (G) ENERGY, (H) LUCKY, (I) ORUS, (J) CLOVIS, (K) MAGNUS, (L) ULYSSE, (M) LUXE, (N) P27174, (O) GR38 AND (P) GRAECUS. The SNP density is represented in 1Mb bins. **(b)** Neighbor-joining phylogenetic tree of white lupin accessions based on SNPs. The 15 accessions are divided in three clades: winter, spring and non-domesticated/landrace. **(c)** Legume evolutionary history. Evolutionary scenario of the modern legumes (white and narrow-leafed lupin, garden pea, peanut, Lotus, barrel medic, chickpea, pigeonpea, soybean, common bean, mungbean, adzuki bean) from the reconstructed ancestral legume karyotype (ALK, center). The modern genomes are illustrated with different colors reflecting the origin from the ancestral ALK chromosomes. Polyploidization events are shown with red (duplication) and blue (triplication) dots on the tree branches. **(d)** Syntenic regions inside white lupin genome. The colored lines link colinearity blocks that represent syntenic regions that are bigger than 50 kb.

We selected the wild accession GRAECUS and an Ethiopian landrace (P27174) to further investigate the impact of domestication on WL genome. These two genotypes strongly differ in their seed protein content, with GRAECUS accumulating high molecular weight ß-conglutin precursors that have allergenic properties^10^ and may have been counter-selected during domestication (Extended Data Fig. 3a, Supplementary Note 4). In addition, both the Ethiopian landrace and GRAECUS accumulates high levels of alkaloids compared to the sweet (low-alkaloid) AMIGA variety (Extended Data Fig. 3d). In order to further characterize the mechanisms controlling these seed quality related traits, we sequenced these two genotypes using Nanopore long-read technology, at a depth of 27.6x and 32.4x for GRAECUS and P27174, respectively (Supplementary Note 3) and generated *de novo* assemblies that demonstrated high level of structural variations (Extended Data Fig. 4a, Supplementary Table 3, Supplementary Note 3). Genome analysis allowed us to identify a family of 6 ß-conglutins that accumulates in the seeds (Supplementary Note 4, Extended Data Fig. 3a-c, Supplementary Tables 4-5), some of which represent good candidates potentially involved in allergenic responses to WL. With respect to the anti-nutritional alkaloids, we identify a list of 66 candidates containing several metabolic enzyme-coding genes that lie within the *pauper* QTL^12,16^, known to be responsible for the lack of alkaloids in the sweet variety AMIGA (Extended Data Fig. 3d-e, Supplementary Table 7).

We retraced the paleohistory of 12 legume genomes including WL and covering the Genistoid, Dalbergioid, Galegoid and Millettoid clades. Independent blocks of synteny (Supplementary Note 5) allowed the identification of an ancestral legume karyotype (ALK) made of 17 conserved ancestral regions (CARs), Supplementary Table 7. This revealed specific polyploidization events in the case of soybean and lupins (WL and narrow-leafed lupin, NLL), giving rise to mosaic genomes composed of 17 shuffled CARs (Fig. 2c). From the ALK, NLL and WL experienced a common triplication event that led to a 1:3 chromosomal relationships between ALK and these lupin genomes (Extended Data Fig. 5a), from which WL differentiated from NLL through 8 major species-specific chromosomal shuffling events (fusions and fissions, Extended Data Fig. 5b). The modern WL genome most likely experienced a 51-chromosome intermediate followed by a massive chromosome number reduction through fusions to reach the 25 modern chromosomes. An intragenomic analysis for segmental duplications (Fig. 2d) identified 928 blocks bigger than 10kb pinpointing a triplication feature that can be observed in several chromosomes segments (*e.g.* Chr07, which has two homologue regions with Chr12 and one with Chr13). Reciprocal pair-wise comparisons^17^ of the 38,258 WL genes with 104,607 genes from its closest relative NLL, the model legume *Medicago truncatula*^18^ and *Arabidopsis thaliana* identified 25,615 orthologs clusters (Extended Data Fig. 4b). 473 out of these groups contain only WL paralog genes (1,242 in total), probably as a result of the predicted genome triplication event (Supplementary Table 10).

Most terrestrial plants can form mycorrhizal symbioses that greatly improve mineral nutrition. Lupins however, lost the ability to form such associations (est. 12-14 My) and some species developed cluster roots *ca.* 2.5My ago^6^. This resulted in the loss of all mycorrhizal specific genes in the WL genome. As part of the legume family, WL can form nitrogen-fixing nodules upon interaction with *Bradyrhizobium lupini*. We identified that common symbiotic genes such as *LaSYMRK* and *LaCCamK* remained present in WL genome and expressed in AMIGA roots (Supplementary Note 6). This suggests that WL adapted to environments where fungi partners may not thrive and instead favored a new type of root adaptive mechanism towards phosphate acquisition while maintaining the ability to symbiotically fix atmospheric nitrogen; conferring tremendous adaptation abilities to this plant^19^.

Despite the importance of cluster roots in nutrient acquisition, no gene controlling cluster root development has been described to date. We therefore generated a detailed transcriptomic dataset of WL cluster root developmental zones. Our RNA-seq survey (mRNA and miRNA) covered 8 sections of mature clusters that mimic the temporal stages of their development (Fig. 3a and Supplementary Note 6). Mature rootlets (S6 and S7) showed the highest number of up-regulated genes, compared to an ordinary lateral root that does not form cluster roots (Fig. 3b). This set of genes have a strong enrichment in GO terms associated with membrane components linked with their highly active physiology required to remobilize and acquire phosphate efficiently (Supplementary Figs. 14-17). Interestingly, a list of 42 genes overexpressed in all cluster root sections (Fig. 3b), showed a strong enrichment in transcription factors (43%). Nine of these belong to the AP2/EREBP family^20^, key regulators of several developmental processes^21–24^ (Fig. 3c). Among this family, 14 genes are miRNA targets, including 5 transcription factors (Supplementary Table 15), suggesting a tight regulation of their expression during cluster root formation.

**Figure 3:**
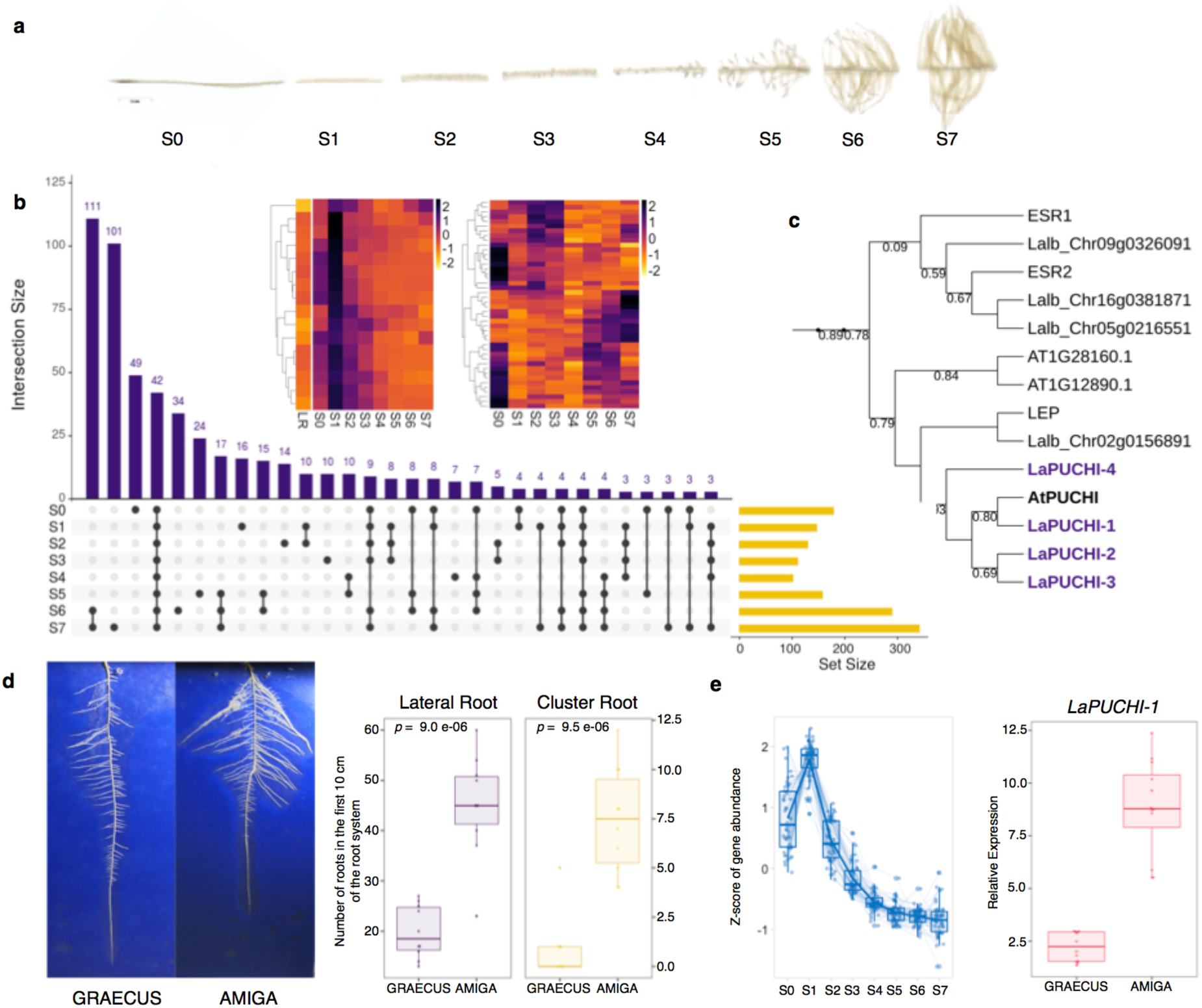
Molecular events associated with early root system establishment in white lupin. **(a)** 8 developmental stages of cluster root development used for transcriptomic studies, showing the formation of numerous rootlets. **(b)** Matrix layout for all intersections of up-regulated genes in the 8 CR segments. For each comparison, the number on top of each bar represents the number of differentially expressed genes (intersection size). The comparison in question is indicated by the dots or connected dots below its respective bar. Set size indicates the total number of genes for each comparison. In detail the heatmap of the 16 genes over expressed in S1 region (left) and the miRNA expressed in the entire CR **(c)** Partial phylogenetic tree of AP2/EREBP subfamily B-1 of Arabidopsis and white lupin orthologs, highlighting the 4 PUCHI orthologs in lupin. **(d)** Left. Visualization of white lupin root system in 2D. Right. Number of lateral roots and cluster roots in GRAECUS (wild) vs. AMIGA (modern) varieties in the first 10 cm of the root system (n=10). **(e)** Left. Expression pattern of 63 genes that are overexpressed in S1 region, which include *LaPUCHI-1*. Right. Relative expression of *LaPUCHI-1* in AMIGA and GRAECUS in top lateral roots of 11-day-old plants.

The impact of domestication on these candidate genes and WL soil exploration capacity was determined using a 2D-phenotyping platform. We identified that the root system architecture of AMIGA develops earlier than the wild-relative GRAECUS as a result of a strong increase in lateral and cluster root number (Fig. 3d). This difference was correlated with an increased level of expression of the regulatory gene *LaPUCHI-1* (Fig. 3c and 3e), an ortholog of a gene that is required for morphogenesis of the lateral root primordium in Arabidopsis^23^. A list of candidate genes selected on their high induction level at the S1 stage (*LaCLE1, LaMYB1, LaPEP1, LaPME41* and *LaSTART*), also showed a higher expression level in AMIGA compared to its wild relative GRAECUS (Extended Data Fig. 6). This suggests that activation of key regulatory genes can trigger the early establishment of the root system, a trait that has been characterized in other crops to be key for more efficient phosphate acquisition (*e.g.* the *pup1* QTL in rice where the *PSTOL1*^25^ genes controls early root development).

The striking ability of plants from 10 different botanical families^7^ to produce cluster roots (including monocots from the Cyperaceae family) questions whether these developmental structures appeared independently several times during evolution due to the lack of mycorrhizal associations in these species or whether they were present in a common ancestor and subsequently lost in most plants. The high quality genome sequence of WL, the only annual legume crop producing cluster roots and showing a reduced need for phosphate fertilizers, will help to understand the molecular mechanisms behind these adaptations. Since phosphate is a limited resource^26^, it could represent an important trait for the future improvement of nutrient acquisition in other crops. Our reference genome sequence and companion resources will also help reinforce breeding programs of WL aimed at reducing seed allergenicity, improving yield stability and maintaining a low content of anti-nutritional alkaloids.

## ACKNOWLEDGMENTS

This project has received funding from the European Research Council (ERC) under the European Union’s Horizon 2020 research and innovation program (Starting Grant LUPINROOTS - grant agreement No 637420 to B.P.). We acknowledge Lucie Combes-Soia and Valerie Labas (Physiologie de la Reproduction et des Comportements, UMR85, INRA, CNRS, Université de Tours, France), Olivier Bouchez (GeT-PlaGe, INRA Auzeville, Castellan Toulosan, France), and Charles Poncet and Veronique Gautier (Genotyping Platform Gentyane, Clermont-Ferrand, France). This work was partly performed in the LRSV and LIPM, which belong to the TULIP LABEX (ANR-10-LABX-41). This work was also partly performed in IPS2, which belong to the SPS LABEX (ANR-10-LABX-40). We also thank the VILLUM Foundation for the Young Investigator grant awarded to F.G.-F. (Project 15476) and funding from Innovate UK to M.N.

## AUTHORS CONTRIBUTIONS

W.M., S.A. and H.B. performed DNA extraction for single-molecule sequencing and optical mapping analysis. J. G. assembled the AMIGA genome. E.S. and S.C performed genome annotation. A.M., R.G. and V.S performed identification and analysis of repeated elements. A. S. developed bioinformatic resources and performed GRAECUS and P27174 assembly. C.H. and J.S. performed paleogenomic analysis. K.G.-G. and D.A. performed seed protein analysis. D. M., F. G.-F., J.T. and M.N. performed analysis of alkaloid pathway. T.B ad M.C. performed miRNA analysis.P.D. and M.L. performed cluster root studies and analysis. L.M. and F.D. performed expression and root architecture studies. J.K. and P.-M. D. performed the identification of symbiotic genes. B.H. performed experiments, data collection and analysis. B.H. and B.P. designed experiments and wrote the article.

## METHODS

### Genome assembly and annotation of *Lupinus albus* L. cv. AMIGA

A meta-assembly strategy similar to the one developed to assemble the Rosa genome^6^ was applied. The Supplementary Table 1 provides details of the different steps of the process including data, software and the evolution of the metrics of the assembly. Firstly, three assemblies were performed with CANU^1^ using different level of stringency (errorRate=default, 0.015 and 0.025m respectively). Corrected reads generated by CANU were also used to run FALCON^7^. The graph of overlaps of FALCON was filtered using three different sets of parameters of the program til-r ^6^, in order again to generate alternative assemblies with different level of stringency.

The N50 metrics of the primary assemblies ranged from 1.6 to 7.1Mb. The sequences of these six primary assemblies were first transformed in pseudo long reads of 100kb with an overlap of 50kb. Then, the pseudo long reads were assembled with CANU 1.6 in the mode –trim-assemble to enable the trimming of sequence ends specific to a single primary assembly.

The meta-assembly result displays a N50 of 8.9Mb in only 129 contigs. The Bionano hybridScaffold.pl software was run in order to scaffold the contigs of the meta-assembly using the Bionano Optical map (N50 2.3Mb). 15 putative breakpoints were identified and corrected by the scaffolder. The scaffolds were polished twice, firstly using arrow and the pacbio raw data mapped with blasr, then with Pilon^8^ using 100x of illumina data mapped with glint software (http://lipm-bioinfo.toulouse.inra.fr/download/glint/). Finally the pseudo-chromosomes were obtained with ALLMAPS^9^ by scaffolding the polished scaffolds with the high density genetic map^10^. A total 96.2% of the data were anchored on the linkage map and 95.3% were oriented (Extended Data Fig. 1). Detailed information about the genome annotation is presented in Supplementary Note 1.

### Evaluation of AMIGA heterogeneity

In order to evaluate the heterogeneity of cv. AMIGA, a bulk of 90 AMIGA plants was resequenced using Illumina HiSeq300, with paired-end 2×150 bp reads. This produced 193,734,276 clean reads corresponding to a total of 64.47x depth. Cutadapt^1^ has been used to remove Illumina Truseq adapter from the sequencing data and to remove bases with a quality score lower than 30,in both 5’ and 3’ end of the reads. Reads with a length lower than 35 have been discarded. We used BWA-MEM version 0.7.17^2^ to map the resequencing reads to the white lupin reference genome. Picard tools (https://github.com/broadinstitute/picard/issues) have been used to detect and remove PCR and Optical duplicates. We then used GATK 4.0^3^ HaplotypeCaller tool to call variants. This identified *ca*. 300,000 SNPs without filtering the data. All the SNPs are evenly distributed on the 25 chromosomes and contigs. We generated a VCF file with this information, available in the white lupin Genome Browser.

### Assembly of chloroplast genome

A *de novo* assembly protocol was used to assemble both cytoplasmic genomes. The chloroplast genome was generated using NOVOPlasty 3.2^4^, by using the aforementioned Illumina reads, after adapter-removing step. The assembled chloroplast genome (plastome) resulted in a single circularized contig of 151,915 bp. Annotation of the plastome was made based on the already available *L. albus* plastome (GenBank accession NC_026681).

### Annotation of repeats

Identification and characterization of moderately to highly repeated genomic sequences was achieved by graph-based clustering of genomic Illumina reads using RepeatExplorer2 pipeline ^5^. A total of 1,144,690 of 150bp paired reads, representing approx. 0.5× genome coverage, were used for the clustering and the 145 largest clusters with genome proportions of at least 0.01% were examined in detail. Clusters containing satellite DNA (satDNA) repeats were identified based on the presence of tandem sub-repeats within their read or assembled contig sequences with TAREAN^6^. Genome-wide TE repeat annotation was performed using the DANTE (Domain-based ANnotation of Transposable Elements) tool^6^. Consensus sequences of satDNA repeats and rDNA genes were used to perform genome-wide annotation of satDNA and rDNA arrays using the Geneious v. 9.1.8 annotation tool (https://www.geneious.com). The generated GFF3 files were further incorporated on the *L. albus* genome browser.

### Chromosome preparation for *in situ* hybridization

Chromosome preparations for *in situ* hybridization analysis were conducted as described previously^7^ with modifications. First, young roots (pre-treated with 8-hydroxyquinoline 2mM for 3-5 h at room temperature) and anthers were fixed in 3:1 (ethanol:acetic acid) for 2-24 h. The fixed tissues were treated with an enzyme mixture (0.7% cellulase R10, 0.7% cellulase,1.0% pectolyase, and 1.0% cytohelicase in 1× citric buffer) for 1h at 37 °C. Material was then washed twice in water and fragmented in 7 μl of 60% freshly prepared acetic acid into smaller pieces with the help of a needle on a slide. Another 7 μl of 60% acetic acid was added, and the specimen was kept for 2 min at room temperature. Next, a homogenization step was performed with an additional 7 μl 60% acetic acid and the slide was placed on a 55 °C hot plate for 2 min. The material was spread by hovering a needle over the drop without touching the hot slide. After spreading of cells, the drop was surrounded by 200 μl of ice-cold, freshly prepared 3:1 (ethanol:acetic acid) fixative. More fixative was added and the slide was briefly washed in fixative, then dipped in 60% acetic acid for 10 min and dehydrated in 96% ethanol. The slides were stored until use in 96% ethanol at 4 °C.

### Probe preparation and fluorescence *in situ* hybridization

FISH probes were obtained as 5′-Cy3 or 5′-FAM-labeled oligonucleotides (Eurofins MWG Operon, http://www.eurofinsdna.com), or were PCR-amplified as described below. All DNA probes, except oligonucleotides, were labeled with Cy3- or Alexa 488-dUTP (Jena Bioscience) by nick translation, as described previously^8^. The sequences of all oligonucleotides and primers are listed in Supplementary Note Table 5. FISH was performed as described previously^7^. Probes were then mixed with the hybridization mixture (50% formamide and 20% dextran sulfate in 2× SSC), dropped onto slides, covered with a cover slip and sealed. After denaturation on a heating plate at 80°C for 3 min, slides were hybridized at 37 °C overnight. Post-hybridization washing was performed in 2× SSC for 20 min at 58°C. After dehydration in an ethanol series, 4′,6–diamidino-2–phenylindole (DAPI) in Vectashield (Vector Laboratories, http://www.vectorlabs.com) was applied. Microscopic images were recorded using a Zeiss Axiovert 200M microscope equipped with a Zeiss AxioCam CCD. Images were analyzed using the ZEN software (Carl Zeiss GmbH). Primer and oligo-probes information is presented in Supplementary Note 2.

### PCR amplification of tandem repeat and retroelement fragments for probe labeling

Fragments for probe labeling were amplified using genomic DNA from L. albus using the forward and reverse primers as supplied on Supplementary Note Table 5. Eight PCR reactions for each target repeat were performed in 50 μL reaction volume containing 100 ng of gDNA, 1 μM primers, 1× PCR buffer, 0.2 mM dNTPs, and 1U of Taq polymerase (Qiagen). Thirty-five amplification cycles with proper conditions for each set of primers were run. PCR reactions were sampled, purified and concentrated using Wizard® SV Gel and PCR Clean-Up System (Promega). Sanger sequencing confirmed correct amplification of PCR fragments. After confirmation, the PCR products containing the same class of repeat were collected and used for probe labeling by nick translation as described above.

### LalbCENH3-ChIP and ChIP-seq analysis

Chromatin immunoprecipitation experiments were done with Abcam ChIP Kit - Plants (ab117137) following the manufacturers instructions. First, 1 g of young *L. albus* cv. AMIGA leaves were collected and cross-linked with formaldehyde 1% for 15 min on ice. Leaves were then ground in liquid nitrogen and sonicated using a Diagenode Sonicator. Sonicated chromatin-DNA ranging from 200-1000 bp was immunoprecipitated using anti-LalbCENH3. Immunoprecipitated DNA and, as a control, an input chromatin DNA samples (3-7ng for each sample) were sent for ChIPseq at BGI. The original ChIPseq sample data are available at White Lupin Genome Website (http://www.whitelupin.fr). To identify repeats associated with CENH3-containing chromatin, reads from the ChIPseq experiment obtained by sequencing DNA from isolated chromatin prior to (the input control sample) and after immunoprecipitation with the CENH3 antibody (the ChIP sample) were separately mapped to the repeat clusters. The mapping was based on read similarities to contigs representing individual clusters, using BLASTn (22) with parameters “-m 8 -b 1 -e 1e-20 -W 9 -r 2 -q -3 -G 5 -E 2 -F F” and custom Perl scripts for parsing the results. Each read was mapped to a maximum of one cluster, based on its best similarity detected among the contigs. Ratio of ChIP/input reads assigned to individual clusters was then used to identify repeats enriched in the ChIP sample as compared to the input.

### White lupin diversity analysis

We selected 14 white lupin accessions, covering a wide range of diverse material, to evaluate a broader range of the genetic diversity and determine population structure and linkage disequilibrium. More information about these accessions can be found in Supplementary Note 3.

#### Data generation with short-reads technology

Young leaves of 30 plants were used to extract genomic DNA of each accession using the QIAGEN Genomic-tip 100/G kit following the supplier’s recommendations. The accessions were sequenced using Illumina technology using paired-end 2×150 bp short-reads. It was generated a total of 310.95 Gb of data with average sequencing depth of 45.99x (Supplementary Note Table 6).

#### Mapping and SNP detection

Cutadapt^1^ was used to remove Illumina Truseq adapters from the sequencing data and to remove bases with a quality score lower than 30, in both 5’ and 3’ end of the reads. Reads with a length lower than 35 were discarded. We then used BWA-MEM version 0.7.17^2^ to map the resequencing reads from all 15 genotypes to the white lupin reference genome. PCR and Optical duplicates have been detected and removed using Picard Tools. After that, GATK 4 HaplotypeCaller tool have been used in emit-ref-confidence GVCF mode to produce one gvcf file per sample. These files have been merged using GATK CombineGVCFs. Finaly, GATK GenotypeGVCFs have been used to produce a vcf file containing variants from all the 15 samples. This identified a total of 6,620,353 SNPs/indel. After filtering for minimum allele frequency of 0.15 and heterozygosity frequency of 0–0.2, 2,659,837 SNPs were retained to further analysis.

#### Phylogenetic analysis, population structure and linkage disequilibrium

The genetic distance matrix was calculated based on identity-by-state similarity method and an average cladogram constructed using neighbour-joining algorithm implemented on TASSEL 5.2.51^9^. Then, a phylogenetic tree was prepared using the iTOL v 4.3^10^. A principal component analysis (PCA) was also performed in R^11^ (http://www.R-project.org/) function “prcomp”. A Bayesian model-based clustering method implemented with STRUCTURE v2.3.4^12,13^ was used to investigate the population structure using all the filtered SNPs. The program was run 10 times for each K value, ranging from 1 to 5, with a 1,000 burn-in time and 1,000 iterations. The optimal K value was determined based on the ΔK from the Structure Harvester v0.6.94^14^ program, through Evanno’s test^15^.

### Long-read sequencing and *de novo* assembly of GRAECUS and P27174

#### Sequencing

Long-read sequencing was realized using Oxford Nanopore technology, using a GridION 18.04.1-0, with a software Minknow 1.10.24-1 at platform at Get-PlaGe core facility (INRA, Toulouse, France). Briefly, DNA was used to prepare a library with the ONT It was used the Ligation Sequencing Kit 1D (sqk-lsk109). The DNA was sequenced using a single ONT MinION R9.4 flowcell (FLO-MIN106) for 48h. Base-calling was performed using Albacore 2.1.10-1. This produced 1,280,206 sequences for GRAECUS, corresponding to 12.45 Gb of data with a N50 length of 13.6 Kb (27.6 × of sequencing depth). For the accession P27174 this produced a total of 1,738,579 reads corresponding to 14.59 Gb of data with N50 length of 11.8 Kb (32.36 × of sequencing depth).

#### De novo assembly of GRAECUS and P27174

The *de novo* assembly of the two genotypes were performed using CANU^16^. For P27174-4, two round of correction have been made prior to the assembly step, using the parameters correctedErrorRate=0.16 anc corMaxEvidenceErate=0.15. For GRAECUS, only one round of correction have been made, using minOverlapLength=400, correctedErrorRate=0.16 and corMaxEvidenceErate=0.15. The Illumina paired-end data described in 3.1 were used to polish two times the two genome assemblies using Pilon^17^. BUSCO v 3.0.0^18^ was run on the set of predicted transcripts. The assessment software detected for GRAECUS 96.8% of complete gene models (1142 complete single copy and 188 duplicated respectively) plus 9 additional fragmented gene models. For P27174 97.8% of complete gene models (1125 complete single copy and 220 duplicated respectively) plus 4 additional fragmented gene models. Structural variation of these two accession were performed in Assemblytics software^19^ based on whole genomes alignments generated with MUMmer^20^. Details of the *de novo* genome assembly and analysis of structural variation of these two accessions are provided in Supplementary Note 3.3.

### Evolutionary analysis of Legume genomes

The proposed evolutionary scenario was obtained following the method described in^21^ based on synteny relationships identified between *L. albus* and other 11 legume species. Briefly, the first step consists of aligning the investigated genomes to define conserved/duplicated gene pairs on the basis of alignment parameters (Cumulative Identity Percentage and Cumulative Alignment Length Percentage). The second step consists of clustering or chaining groups of conserved genes into synteny blocks (excluding blocks with less than 5 genes) corresponding to independent sets of blocks sharing orthologous relationships in modern species. In the third step, conserved gene pairs or conserved groups of gene-to-gene adjacencies defining identical chromosome-to-chromosome relationships between all the extant genomes are merged into Conserved Ancestral Regions (CARs). CARs are then merged into protochromosomes based on partial synteny observed between a subset (not all) of the investigated species. The ancestral karyotype can be considered as a ‘median’ or ‘intermediate’ genome consisting of protochromosomes defining a clean reference gene order common to the existent species investigated. From the reconstructed ancestral karyotype an evolutionary scenario was then inferred taking into account the fewest number of genomic rearrangements (including inversions, deletions, fusions, fissions, translocations), which may have operated between the inferred ancestors and the modern genomes. Additional information are provided in Supplementary Note 4.3.

### Genome synteny and intragenomic collinearity

To identify intragenomic colinearity blocks inside the white lupin genome we used SynMap (CoGe, www.genomevolution.org) using homologous CDS pairs using the following parameters: Maximum distance between two matches (-D): 20; Minimum number of aligned pairs (-A): 10; Algorithm “Quota Align Merge” with Maximum distance between two blocks (-Dm): 500.

### Gene family identification

We used a comparative analysis to examine the conservation of gene repertoires among orthologs in the genomes of white lupin, narrow-leafed lupin (v1.0) *M. truncatula* (Mt4.0) and *Arabidopsis thaliana* (TAIR10). First, we aligned all-to-all proteins using BLASTP (e-value of 1e^-5^). Genes were then clustered using OrthoMCL (1.4) implemented in OrthoVenn ^22^ with a Markov inflation index of 1.5 and a minimum e-value of 1e-15.

### Candidate genes controlling cluster root formation

#### Spatial transcriptome for mRNA and small RNA

Ten cluster roots coming from four grown plants were harvested after 12 days of culture and dissected in eight parts of 0.5-cm from the apex of the lateral root that carries the cluster root (Supplementary Note 6.2). As control, 1-cm of lateral roots without cluster roots, sampled 1-cm away from the primary root, were collected. Four biological replications were produced for each experiment. Total RNA was extracted from all frozen samples using the Direct-zol RNA MiniPrep kit (Zymo Research, Irvine, CA) according to the manufacturer’s recommendations.

For mRNA sequencing, 36 independent root RNA-seq libraries were constructed using Illumina TruSeq Stranded mRNA Sample Preparation Kit (Illumina Inc.), according to the manufacturer’s protocol. The samples were sequenced using paired-end sequencing was performed generating paired-ended 2 × 150 bp reads using TruSeq SBS kit v3 sequencing chemistry (Illumina Inc.) in one lane of Illumina NovaSeq instrument according to the manufacturer’s instructions. A total of 2,048,118,650 paired-end reads of 150 pb were sequenced using an Illumina NovaSeq6000 Sequencer. To remove low quality sequences, the RNA-seq reads were checked and trimmed using Cutadapt^1^ with a minimum quality score of 30 in both 3’ and 5’ end, with the nextseq-trim option enabled. Illumina TruSeq adapter sequences have also been removed. The resulting reads shorter than 35 pb have been discarded. The quality checked RNA-seq reads were then mapped on white lupin reference genome using Hisat2^23^ software. Transcripts were assembled and quantified using Stringtie software. Gene counts were extracted and imported in the R package DESeq2^24^. These counts have been normalized according to the size factor computed by DESeq2.

For small RNA sequencing, 24 independent root RNA-seq libraries were constructed using NEXTflex™ Small RNA-Seq kit according to the manufacturer’s protocol. All small RNA libraries were sequenced on an Illumina NextSeq 500 sequencing platform, using a single-end, 75 nt read metric instrument according to the manufacturer’s instructions. A total of 460,506,072 reads of 75 nt were sequenced. Small RNA-seq reads were trimmed using Cutadapt version 1.11^1^ to remove remnants of the following 3’-adapter sequence. Details on the trimming, assembly, differential expression analysis and miRNA family identification can be found in Supplementary Note 6.2.

#### AMIGA and GRAECUS root sampling and expression analysis of cluster root initiation genes

We sampled 2-3 cm of lateral roots 1-cm away from the primary root in the top 5 cm (cluster root region, CRR) and at 10 cm from the top (regular lateral root region, NLR) of the root system of AMIGA and GRAECUS plants, 11 days after germination. Three CRR and 3 NLR independent samples were collected for each accession. Total RNA from these samples was extracted using the Direct-zol RNA MiniPrep kit (Zymo Research, Irvine, CA) according to the manufacturer’s recommendations. RNA concentration was measured on a NanoDrop (ND1000) spectrophotometer. Poly(dT) cDNA were prepared from 2 μg total RNA using the revertaid First Strand cDNA Synthesis (Thermo Fisher). Gene expression was measured by quantitative Real Time - Polymerase Chain Reaction (qRT-PCR) (LightCycler 480, Roche Diagnostics, Basel, Switzerland) using the SYBR Premix Ex Taq (Tli RNaseH, Takara, Clontech, Mountain View, CA) in 384-well plates (Dutscher, Brumath, France). Target quantifications were performed with specific primer pairs described on the Supplementary note table 9. Expression levels were normalized to LaHelicase (Lalb_Chr13g0304501). All qRT-PCR experiments were performed in technical quadruplicates. Relative gene expression levels were calculated according to the ΔΔCt method^25^, using as a calibrator the NLR samples. All experiments were performed as three biological replicates.

## Data availability

The integrative web interface white lupin genome portal: www.whitelupin.fr contains a Genome Browser, Expression tools, a Sequence retriever tool as well as all raw data available for download.

---

**Extended Data Table 1.** Statistics of the white lupin genome and gene models prediction

|  | Number | Size |
| --- | --- | --- |
| **Assembly feature** |  |  |
| Assembled sequences | 89 | 450.972 Mb |
| N50 | 12 | 17.35 Mb |
| N90 | 23 | 14.55 Mb |
| GC content (%) | 33.71 |  |
| **Genome annotation** |  |  |
| TE proportion (%) | 60 |  |
| Annotated protein-coding genes | 38 258 |  |
| Annotated non-coding proteins genes | 3 129 |  |
| Complete BUSCOs | 1 331 (97.7%) |  |
| Fragmented BUSCOs | 3 (0.2%) |  |
| Missing BUSCOs | 29 (2.1%) |  |

**Extended Data Figure 1.**
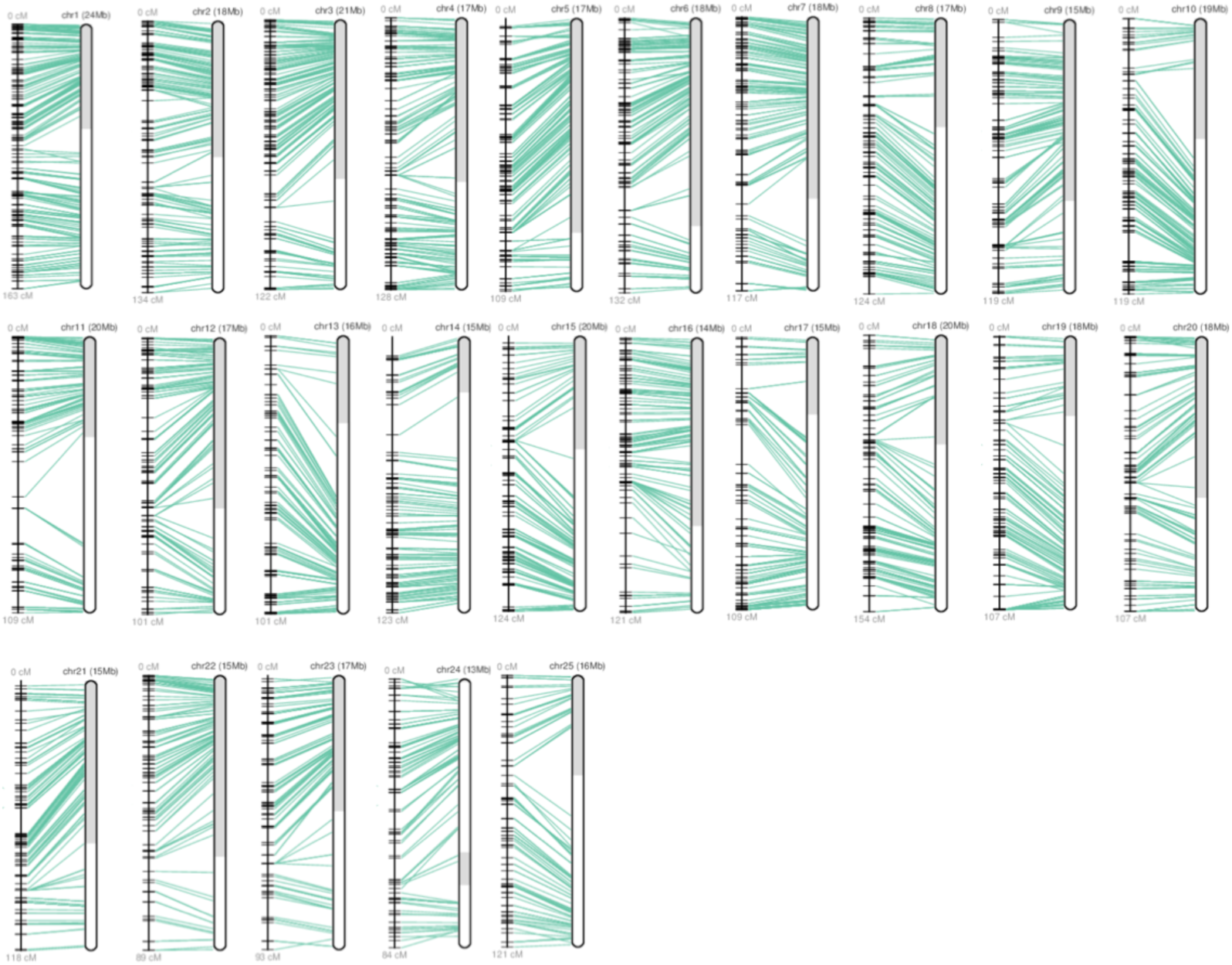
Integration of sequence data and genetic map as provided by ALLMAPS. Each chromosome is represented by its linkage group (right) and chromosome arms (left).

**Extended Data Figure 2.**
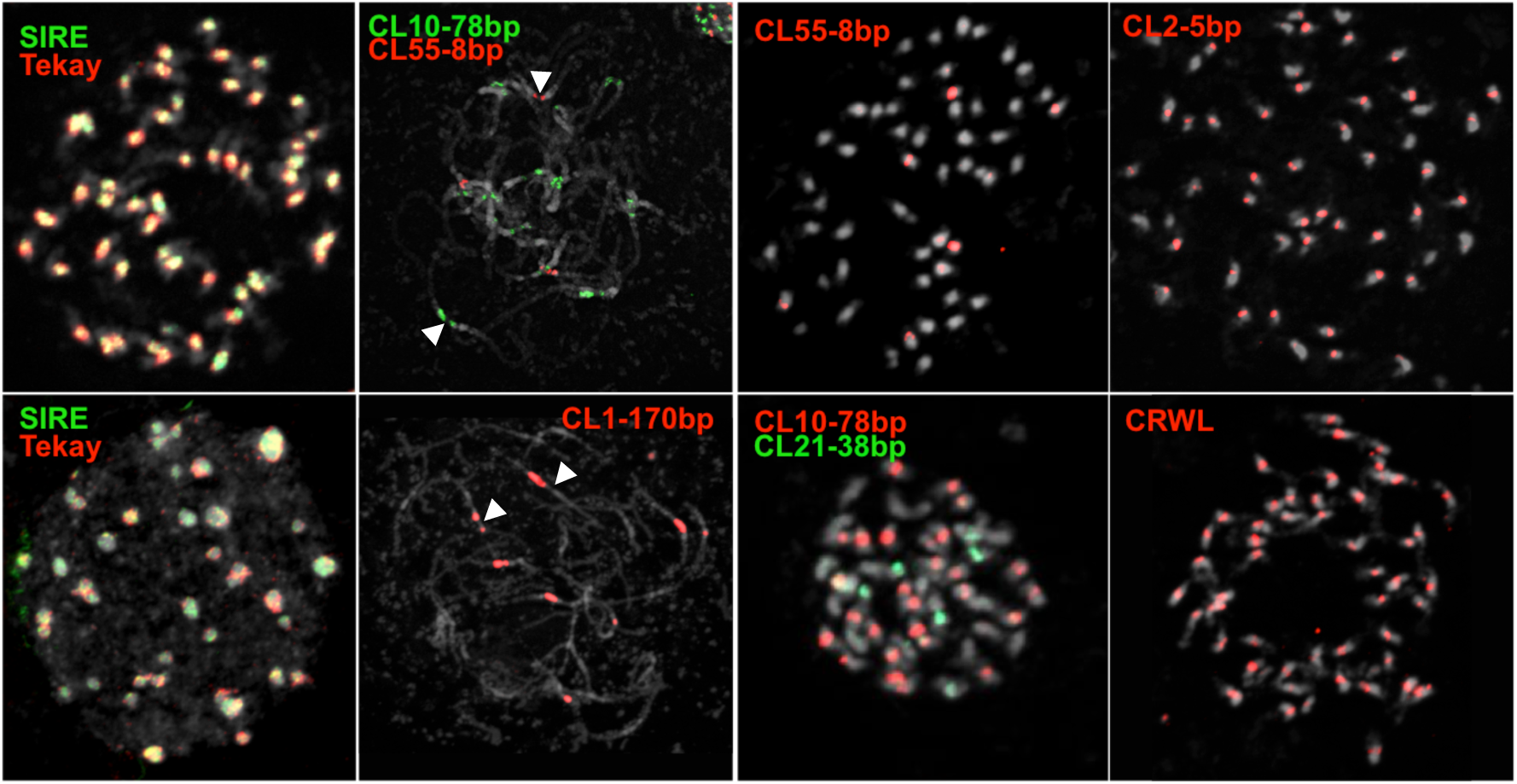
FISH with main repeats of WL genome showing the distribution of pericentromeric and centromeric repeats. Arrowheads point to core centromeres. CRWL, CL2-5bp, CL10-78bp, CL21-38bp and CL55-8bp repeats localize specifically to core centromeres, while CL1-170bp repeat localizes aside core centromeres.

**Extended Data Figure 3.**
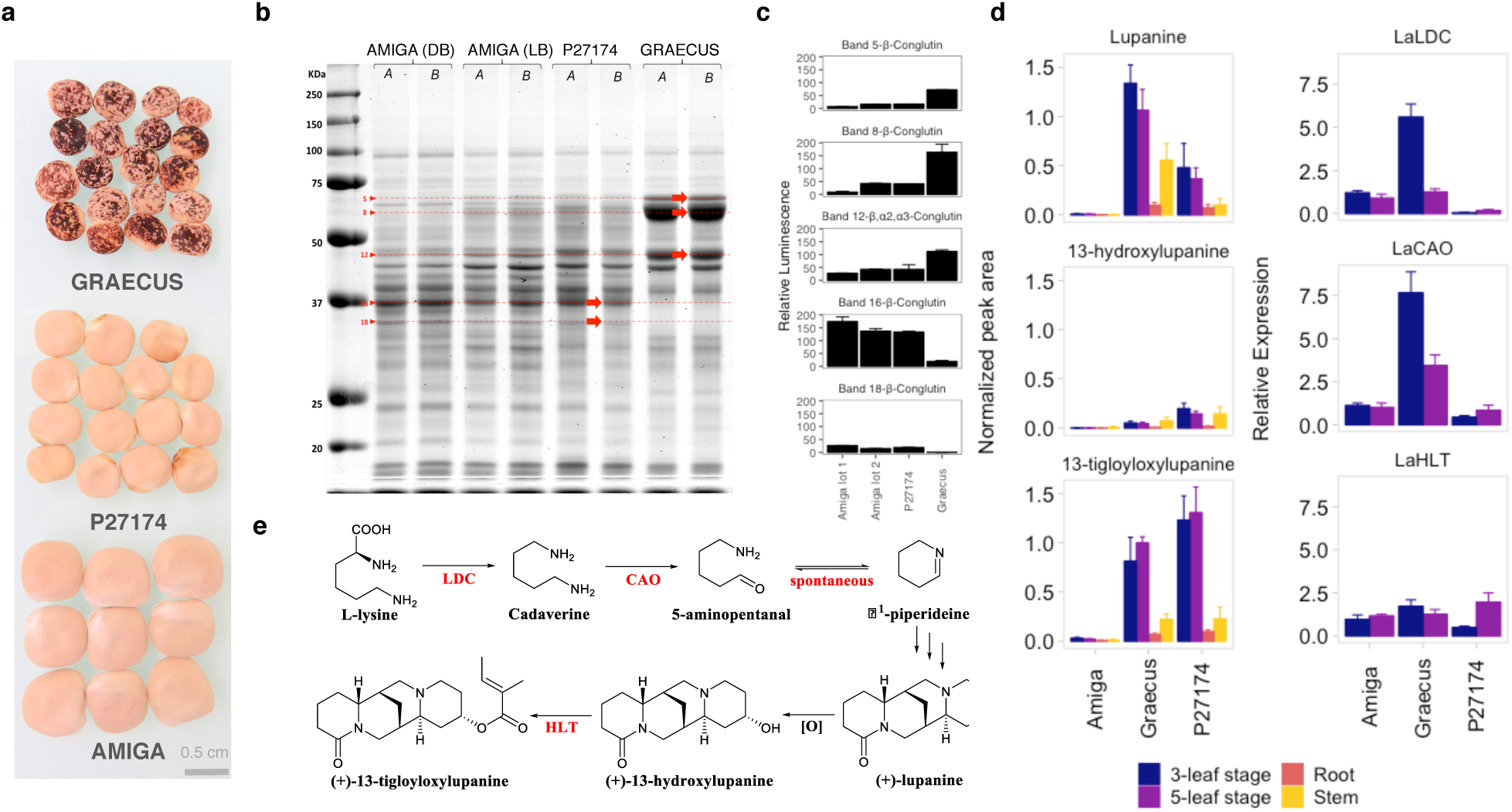
Analysis of white lupin seed protein and leaf alkaloid content. (a) Grain appearance in GRAECUS (wild-type), Ethiopian landrace P27174 and *cv.* AMIGA. **(b)** Seed protein composition of AMIGA (DB, dark brown seeds; LB, light brown seeds), P27174 and GRAECUS (A and B: two independent extractions of proteins using Tris-SDS, separation in 12% SDS-PAGE). The bands extracted for MS/ MS analysis are highlighted with red arrows. **(c)** Protein content quantification of each band extracted for each variety expressed in normalized gel volume. **(d)** Abundance of the three major alkaloids in young leaf (3-and 5-leaf growth stage), stem and root tissues of the three varieties of *L. albus* as measured by LC-MS and expression of the known enzyme-coding alkaloid biosynthesis genes in young leaves as measured by qRT-PCR. All data are expressed as means ± standard deviation (n = 5 for AMIGA and n = 4 for P27174 and GRAECUS). *LDC: lysine decarboxylase; CAO: copper amine oxidase; HLT: 13-hydroxylupanine O-tigloyltransferase*. (e) The putative biosynthetic pathway of tetracyclic quinolizidine alkaloids in lupins. Characterized steps and respective enzyme names are marked in red.

**Extended Data Figure 4.**
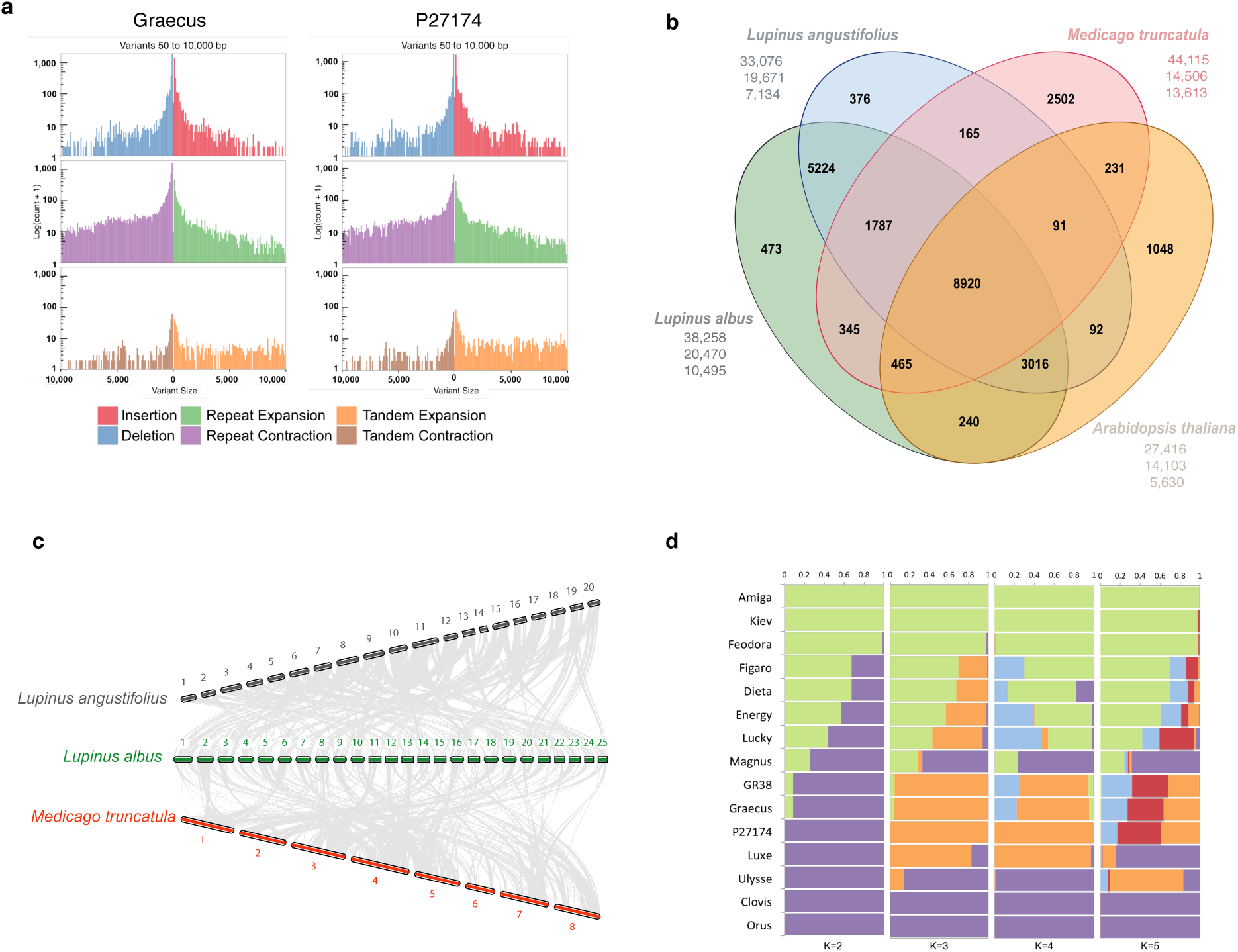
**(A)** Structural variants between *L. albus* cv. AMIGA and the *de novo* assembly of GRAECUS (left) and P27174 (right). The biggest proportion of variants are the repeated elements. **(b)** OrthoMCL clustering of white lupin genes with those of *L. angustifolius, M. truncatula* and *A. thaliana*. Numbers in the sections of the diagram indicate the number of clusters (gene groups). The first number below each species name is the total number of genes of the species, the second number is the number of genes in clusters and the third number is the number of genes that did not cluster. **(c)** Synteny blocks shared between white lupin, its close relative *L. angustifolius* and the legume model *Medicago truncatula.* **(d)** Admixture representation of the 15 accessions with population clustering for K=2-5. Each individual is represented by a horizontal bar and each color represents a subpopulation. The color of each individual accession represents their proportional membership in the different populations.

**Extended Data Figure 5.**
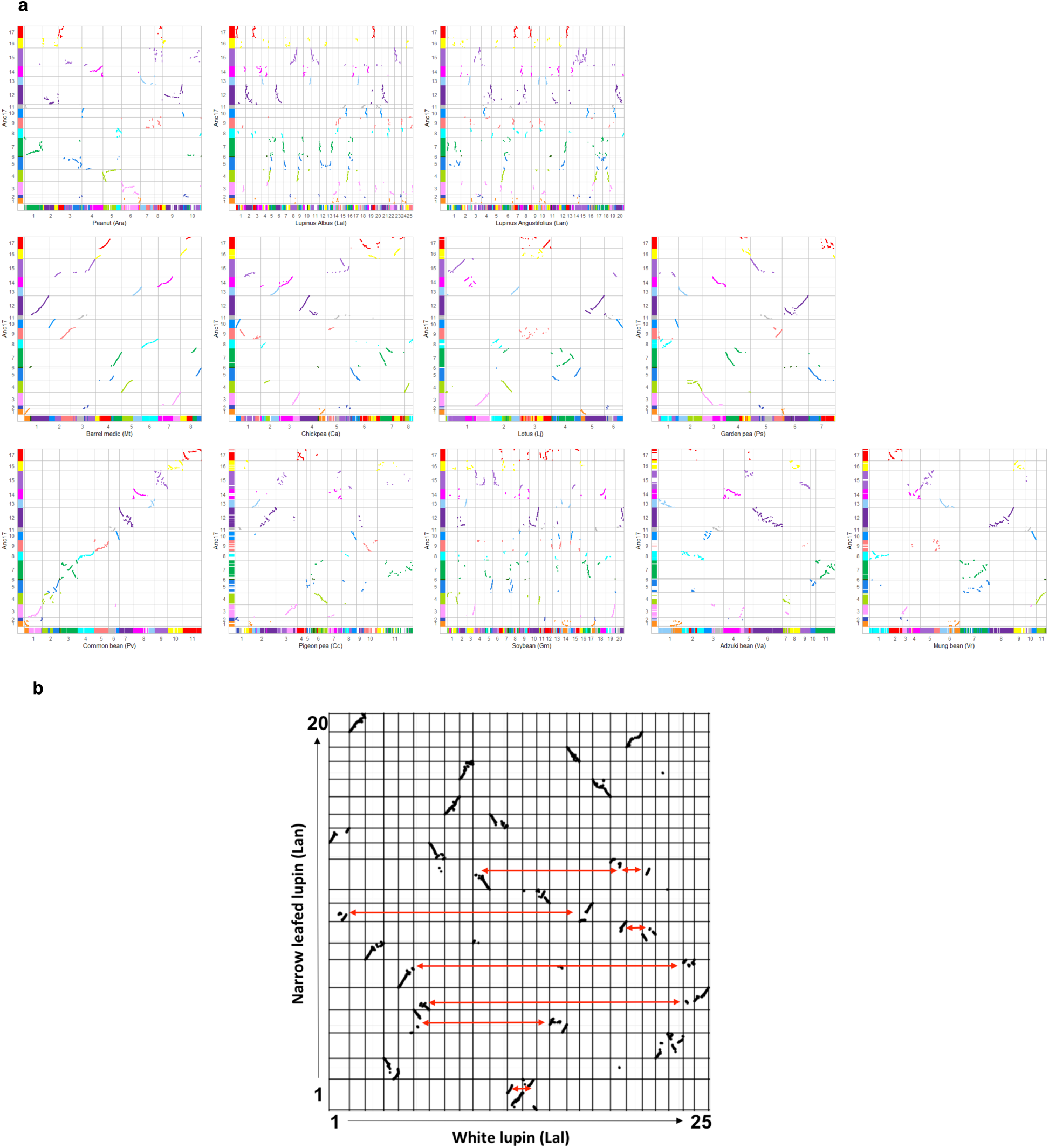
Legume genome synteny. **(a)** Dotplot-based deconvolution of the synteny relationships between Ancestral Legume Karyotype (y-axis) and the 12 legume genomes *(x-axis).* The chromosomes are depicted as a mosaic of a 17 color-codes reflecting the 17 inferred CARs. The synteny relationships identified between the ancestral genome and the modern species are illustrated with colored diagonals in the dotplot. **(b)** Dotplot-based deconvolution of the synteny relationships between white and narrow leafed lupin highlighting 8 major chromosomal shuffling events (red arrows).

**Extended Data Figure 6.**
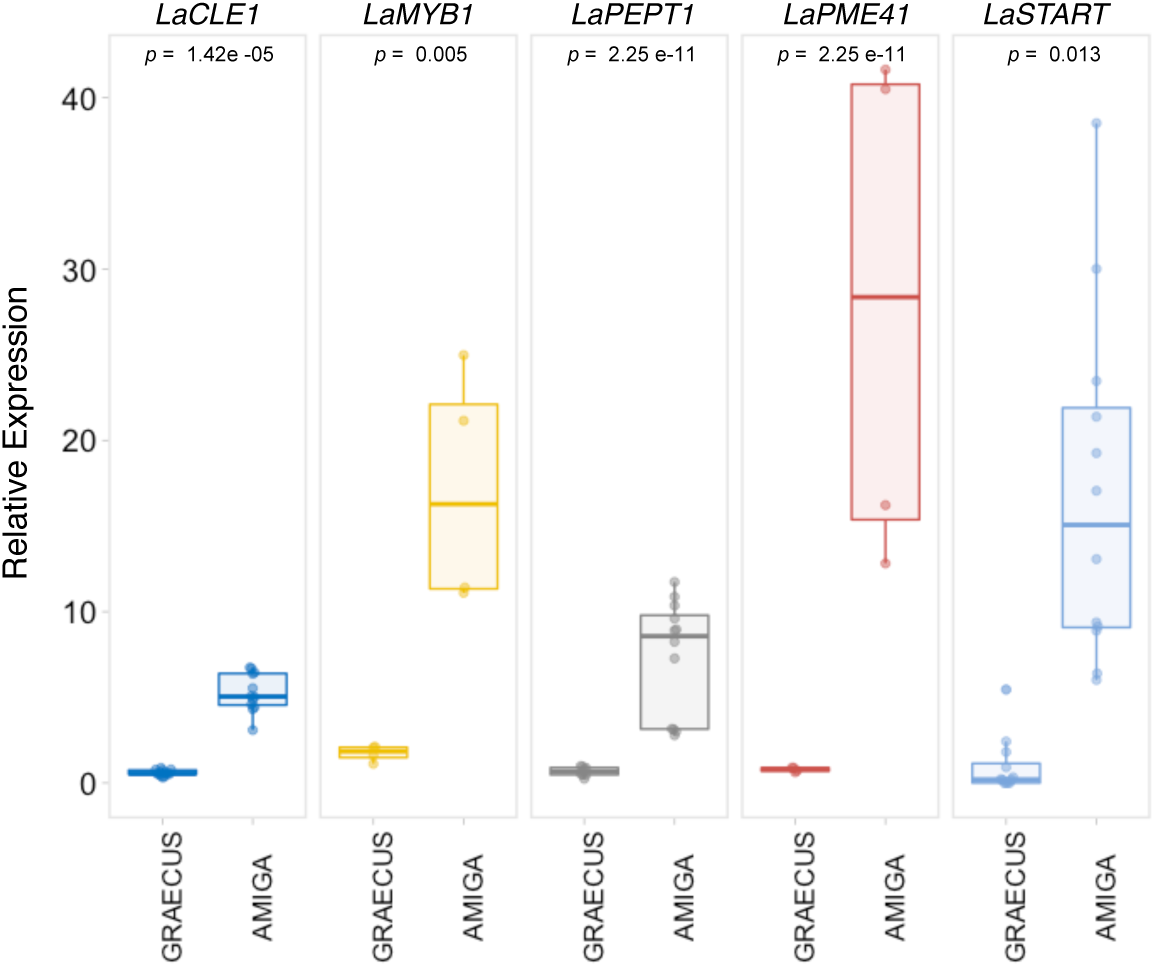
Relative expression level in top lateral roots of AMIGA and GRAECUS of 5 genes overexpressed in the cluster roots S1 developmental zone. The *L. albus* genes LaSTART (Lalb _ Chr20g0121101), LaCLE-1(Lalb_Chr19g0133921), LaPEPT-1 (Lalb_Chr16g0388291), LaPME41(Lalb_Chr06g0166531) and LaMYB1 (Lalb_Chr20g0122341) are overexpressed in the in the top lateral roots of the cultivated variety AMIGA. This suggests that activation of key regulatory genes can trigger the early establishment of the CR and the root system.

