## Supplementary Notes and Figures for "Genome sequence of the cluster root forming white lupin"

Hufnagel *et al.*

#### SUPPLEMENTARY NOTE

Table of Contents

**1 Genome sequencing and assembly** 3
**1.1 Single-molecule real-time and short-read sequencing** 3
**1.2 Optical maps** 3
**1.3 K-mer genome size estimation** 4
**1.4 Genome assembly** 4
**1.5 Assembly of chloroplast and mitochondrial genomes** 5
**1.6 Genome annotation** 5
1.6.1 Generation of Illumina RNA-seq data from 10 white lupin tissues 5
1.6.2 Prediction of gene models 8
1.6.3 Functional annotation of protein-coding genes 8
**1.7 Evaluation of AMIGA heterogeneity** 11
**1.8 White lupin genome portal** 11
**2 Transposable elements and annotation of repeated sequences** 11
**2.1 Material and Methods** 11
2.1.1 Plant material for cytological analysis 11
2.1.2 Chromosome preparation for in situ hybridization 12
2.1.3 Repeat identification and genome-wide repeat annotation 12
2.1.4 Probe preparation and fluorescence in situ hybridization. 13
2.1.5 PCR amplification of tandem repeat and retroelement fragments for probe labeling 14
2.1.6 Identification and generation of LalbCENH3 antibody 15
2.1.7 Immunostaining 15
2.1.8 Super-resolution microscopy 15
2.1.9 CENH3-ChIP and ChIP-seq analysis 16
**2.2 Results** 16
**3 White lupin diversity analysis** 20
**3.1 Methods** 21
3.1.1 Data generation with short-reads technology 21
3.1.2 Mapping and SNP detection 21
3.1.3 Phylogenetic analysis, population structure and linkage disequilibrium 23
**3.2 Results** 23
3.2.1 Results 23
**3.3 Long-read sequencing and *de novo* assembly of GRAECUS and P27174** 24
3.3.1 Methods 24
3.3.2 Results 26
**4 Seed Quality** 26
**4.1 Identification and characterization of seed storage protein** 26
4.1.1 Methods 26
4.1.2 Results 30

### 1 Genome sequencing and assembly

#### 1.1 Single-molecule real-time and short-read sequencing

Genomic DNA was extracted from young leaves of a one single seed descendent (S2 generation) derived plant of *Lupinus albus* cv. AMIGA (from Florimond Desprez, France) using the QIAGEN Genomic-tip 100/G kit following the supplier's recommendations. Sequencing was sub-contracted at Gentyane core facility (INRA, Clermont-Ferrand, France), using a PacBio Sequel E406 with 30 SMRT Cells (V2 chemistry, 6h run), yielding a total of 94.2 Gbp of data and estimated genome coverage of 164x (Supplementary Notes Table 1).

**Supplementary Note Table 1. PacBio White lupin genome sequencing data.** Maximal and mean values (from 30 SMRT cells) relative to sequencing efficiency are indicated. The N50 value indicates that 50% of the sequenced nucleotides are contained in reads equal to or larger than this value.

| | Number of reads | Max read Length (bp) | N50 (bp) | Mean read length (bp) | Median read length (bp) | Sequence data per SMRT cell (Gb) |
|---|---|---|---|---|---|---|
| **Max** | 733 203 | 18 3920 | 17 575 | 11 825 | 10 795 | 6.67 |
| **Mean** | 345 841 | 89 407 | 15 047 | 9 394 | 7 842 | 3.14 |

DNA genomic from the same extraction described above was used to generate short-reads using one lane of Illumina HiSeq300, with paired-end 2x150 bp reads at Get-PlaGe core facility (INRA, Toulouse, France). It produced 794,235,810 clean reads corresponding to a total of 119 Gb of reads (208x depth).

#### 1.2 Optical maps

Leaves were collected from two white lupin plants (three-leaf seedling stage) grown in the dark for 3 days and originating from the same parent as the plant used for long and short read sequencing. High Molecular Weight DNA was extracted with the IrysPrep® Plant Tissue DNA Isolation kit (BioNano Genomics) from 2 g of fresh leaves following the supplier's recommendations. To preserve the DNA integrity and obtain long DNA fragments, nuclei were embedded in agarose before DNA extraction. A first draft assembly generated from the PacBio reads using CANU[1] was analysed with the Label Density CalculatorTM 1.3.0 software (Bionano Genomics) to assess the frequency of recognition sites for different nicking enzymes. We chose the Nb.BssSI nicking enzyme (New England Biolabs), which had a theoretical label density of 10 labels for 100 kb. DNA molecules were nicked with Nb.BssSI and labelled according to the instruction provided with the IrysPrep® NLRS Labelling Kit (BioNano Genomics). The labelled DNA molecules were stretched on IrysChip® V2 (BioNano Genomics) and pictured on the Irys® System (BioNano Genomics). Raw image data were converted by the AutoDetectTM 2.1.4.9159 software (BioNano Genomics) into .bnx files corresponding to digital representations of the DNA molecules carrying the motif-specific label pattern. The *de novo* assembly algorithm from BioNano SolveTM data analysis software aligned the digital molecules for the construction of a consensus genome map. 293 maps with a N50 of 2.3 Mb were produced using a selection of molecules above 150 kb. The consensus genome optical map size is 455Mb.

#### 1.3 K-mer genome size estimation

We used the 119 Gb data from the 150bp paired-end Illumina library to perform genome size estimation. The library was submitted to 17-mer frequency distribution analysis using Jellyfish[2] and GCE (Genomic Character Estimator - v1.0.0)[3]. A total of 10,4,671,784,125 17-mers were counted. Given the unique k-mer depth of 174, we calculated that the genome size = KmerCount/Depth = 573.769 Mb. This value is close to the 567 ± 21 Mb described previously based on flow cytometry[4,5].

#### 1.4 Genome assembly

A meta-assembly strategy similar to the one developed to assemble the Rosa genome[6] was applied. The Supplementary Table 1 provides details of the different steps of the process including data, software and the evolution of the metrics of the assembly. Firstly, three assemblies were performed with CANU[1] using different level of stringency (errorRate=default, 0.015 and 0.025 respectively). Corrected reads generated by CANU were also used to run FALCON[7]. The graph of overlaps of

FALCON was filtered using three different sets of parameters of the program til-r [6], in order again to generate alternative assemblies with different level of stringency.

The N50 metrics of the primary assemblies ranged from 1.6 to 7.1Mb. The sequences of these six primary assemblies were first transformed in pseudo long reads of 100kb with an overlap of 50kb. Then, the pseudo long reads were assembled with CANU 1.6 in the mode –trim-assemble to enable the trimming of sequence ends specific to a single primary assembly.

The meta-assembly result displays a N50 of 8.9Mb in only 129 contigs. The Bionano hybridScaffold.pl software was run in order to scaffold the contigs of the meta-assembly using the Bionano Optical map (N50 2.3Mb). 15 putative breakpoints were identified and corrected by the scaffolder. The scaffolds were polished twice, firstly using arrow and the pacbio raw data mapped with blasr, then with Pilon[8] using 100x of illumina data mapped with glint software (http://lipm-bioinfo.toulouse.inra.fr/download/glint/).

Finally the pseudo-chromosomes were obtained with ALLMAPS[9] by scaffolding the polished scaffolds with the high density genetic map[10]. A total 96.2% of the data were anchored on the linkage map and 95.3% were oriented (Extended Data Fig. 1).

#### 1.5 Assembly of chloroplast and mitochondrial genomes

A *de novo* assembly protocol was used to assemble both cytoplasmic genomes. The chloroplast and mitochondrial genomes were assembled using NOVOPlasty[11], by using the aforementioned raw Illumina reads. The assembled chloroplast genome (plastome) resulted in a single circularized contig of 151,915 bp. Annotation of the plastome was made based on the already available *L. albus* plastome (GenBank accession NC_026681).

#### 1.6 Genome annotation

##### *1.6.1 Generation of Illumina RNA-seq data from 10 white lupin tissues*

White lupin material

Plants of *L. albus* cv. AMIGA were grown in growth chambers under controlled conditions (16 h light/8 h dark, 25°C day/20°C night, 65% relative humidity and PAR intensity 200 µmol $m^{-2}$ $s^{-1}$) at B&PMP Institute facilities, Montpellier, France. Tissue was flash frozen in liquid nitrogen and stored at -80 °C. In total, 10 white lupin different samples were isolated for generation of RNA-seq data:

- *Root tissues*: Root tissues were sampled from plants grown in hydroponics conditions. The hydroponic solution was modified from previous medium[12] with (+P, 400 $KH_2PO_4$ µM) or without phosphate (-P, $K_2SO_4$ 200 µM), according to the following composition: $MgSO_4$, 54 µM; $Ca(NO_3)_2$ 400 µM; Na-Fe-EDTA 10 µM; $H_3BO_3$ 2.4 µM; $MnSO_4$ 0.24 µM; $ZnSO_4$ 0.1 µM; $CuSO_4$ 0.018 µM and $Na_2MoO_4$ 0.03 µM. Six different types of root tissues were sampled:
    - AMIGA Roots –P: entire root system of AMIGA grown under –P conditions 20 days after germination;
    - AMIGA Roots +P: entire root system of AMIGA grown under +P conditions 20 days after germination;
    - AMIGA Lateral Roots: mixture of lateral roots of AMIGA (without cluster roots), grown under –P and +P conditions 20 days after germination;
    - AMIGA Primary Roots: mixture of AMIGA primary roots grown under –P and +P conditions 20 days after germination;
    - AMIGA Cluster Roots: mixture of entire cluster roots of AMIGA grown under –P condition 20 days after germination;
    - AMIGA nodulated roots: entire root system of AMIGA in different stages of nodulation after inoculation with bacteria *Bradyrhizobium lupini* strain LL13. Plants were sampled 21 days after germination (18 days after inoculation.
- *Vegetative tissues*:
    - AMIGA Leaves: mixture of AMIGA leaves in different stages of development sampled for plants grown in hydroponic solution (as described in the above section) with +P and –P conditions;
    - AMIGA Flower: mixture of entire flowers of 6 AMIGA plants grown in 2 L pots;
    - AMIGA Pods: mixture of pods of 3 AMIGA plants grown in 2 L pots in controlled conditions.

- *Seeds:*
  - AMIGA seeds: mixture of tissue from 10 mature seeds of AMIGA.

RNA extraction

Total RNA was extracted from all frozen samples using using the Direct-zol RNA MiniPrep kit (Zymo Research, Irvine, CA) according to the manufacturer's recommendations.

Library preparation and Illumina RNA sequencing

RNA-seq libraries were constructed and sequenced at Montpellier GenomiX facilities. The Illumina TruSeq Stranded mRNA Sample Preparation Kit (Illumina Inc.) was used according to the manufacturer's protocol. One lane of Illumina HiSeq 2500 instruments according to the manufacturer's instructions. Paired-end sequencing was performed generating paired-ended 2 x 125 bp reads using TruSeq SBS kit v3 sequencing chemistry (Illumina Inc.). This produced 69.25 Gb of data (276.99 M reads) in an average of 6.9 Gb of data per sample (Supplementary Notes Table 2).

**Supplementary Note Table 2. Summary of data generated in the RNA sequencing of 10 samples of white lupin tissues.**

| Sample | # Filtered reads | Sequence data (Gb) |
|---|---|---|
| AMIGA Root -P | 28 699 273 | 7.17 |
| AMIGA Root +P | 38 270 774 | 9.57 |
| AMIGA Cluster Root | 24 488 390 | 6.12 |
| AMIGA Pods | 22 075 950 | 5.52 |
| AMIGA Flower | 27 047 695 | 6.76 |
| AMIGA Seeds | 24 521 334 | 6.13 |
| AMIGA Lateral Roots | 35 512 338 | 8.88 |
| AMIGA Primary Root | 24 777 784 | 6.19 |
| AMIGA Nodulated Root | 26 467 897 | 6.62 |
| AMIGA Leaves | 25 128 577 | 6.28 |
| *Total* | 276 990 012 | 69.25 |
| *Mean* | 27 699 001 | 6.92 |

##### *1.6.2 Prediction of gene models*

Protein and non-protein coding gene models were predicted using the integrative EuGene pipeline[13] (http://eugene.toulouse.inra.fr/Downloads/egnep-Linux-x86_64.1.4.tar.gz). Three protein databases were aligned to contribute to translated regions detection: i) Swiss-Prot – October 2016 ii) a plant subset of Uniprot proteins – October 2016 and iii) the proteome of *Medicago truncatula*, release 5.1.6[14]. Proteins with similarities to REPBASE[15] were removed from datasets prior alignment. Chained alignments spanning less than 50% of the length of the database protein were removed.

Illumina-based RNAseq datasets described in 1.6.1 were assembled with an iterative *k*-mer strategy based on velvet [16], parameters: -cov_cutoff 4 -read_trkg yes -exp_cov 100 -min_contig_lgth 150 -max_divergence 0.05 -long_mult_cutoff 0) and used as transcriptional evidences by the EuGene pipeline. Spliced alignments spanning at least 30% of the RNA-seq contig length at a minimum of 97% identity were retained. The gene-modelling algorithm used the standard EuGene 4.2a parameters, except that non-canonical GC/donor sites were allowed.

Regions spanned either by Medicago protein hits or by transcript alignments were preserved from the integrated repeat masking process.

After removal of redundant ncRNA predictions, 38,258 protein-coding genes and 3,129 non protein-coding genes were retained. A large proportion of 5-prime and 3-prime untranslated regions of protein-coding gene models were predicted on the basis of transcript alignments (84% and 83% respectively).

The BUSCO v 3.1.0[17] was run on the set of predicted transcripts. The assessment software detected 97.7% of complete gene models (1,142 complete single copy and 201 duplicated respectively) plus 3 additional fragmented gene models.

##### *1.6.3 Functional annotation of protein-coding genes*

Protein coding genes were annotated by integrating five sources of information. Results were successively integrated depending on the expected accuracy of the source of information. Priority was successively given to: i) a BLASTp search of reciprocal best hits with the 1,938 Fabaceae proteins tagged as "reviewed" in the Uniprot database (90% span, 80% identity) as of June 2017 (Uniprot C); ii) 1,523 EC numbers assigned to 10,791 putative enzymes by using the described protocol [18] with BLAST e-value cutoff lowered to $1.e^{-5}$ and pathway-prediction-score set to 0.3 in pathway-tools to increase stringency; iii) the transcription factors and kinases identified (3,284 and 1,358 respectively) by ITAK release 1.7[19] iv) the 4,851 transcription factors identified by PlantTFCat[20] v) the Interpro (release 64.0) search matching 36,298 proteins[21].

The EC numbers were tested against the ENZYME[22] database (November 2017), updated when deprecated and then used to get the description of the enzymes. At each step, the description provided by the source of information was edited when necessary to ensure compliance with a submission to Genbank (Benson DA). Finally, the protein annotations were validated by the tbl2asn software (https://www.ncbi.nlm.nih.gov/genbank/tbl2asn2, November, 2017). A putative function was assigned to 28,019 proteins and 7,625 proteins were tagged as "hypothetical proteins". GO terms were assigned using the BLAST2GO pro software integrating blastp similarities with Genbank NR (May, 2016, e-value < 1e-5, 20 best-scoring hits) and Interpro release 64 results. The final metrics of the assembly and annotation are described in the Supplementary Note Table 3.

**Supplementary Note Table 3. Summary of final assembly and annotation of white lupin cv. AMIGA.**

| | | |
|---|---|---|
| **Total assembly** | | |
| **Total number of nucleotides** | **450,969,408** | |
| Number of nucleotides (without 'N') | 448,704,169 | |
| | Per cent GC | 33.71 |
| **Total number of genes** | **41,387** | |
| | Total nucleotides (bp) | 132,508,751 |
| **Protein coding genes** | | |
| **Number of protein coding genes** | **38,258** | |
| | Mean gene length (bp) | 3,411 |
| | Coding nucleotides (bp) | 42,471,860 |
| | Per cent genes with introns | 78.00 |
| | Per cent genes with 5'-UTR | 84.00 |
| | Per cent genes with 3'-UTR | 83.00 |
| Exons | | |
| | Mean number per gene | 5.13 |
| | Mean length (bp) | 309.47 |
| | GC per cent | 39.28 |
| Introns | | |
| | Mean number per gene | 4.13 |
| | Mean length (bp) | 441.78 |
| | GC per cent | 32.19 |
| CDS | | |
| | Mean length (bp) | 1,110.14 |
| | Min length (bp) | 123.00 |
| | Max length (bp) | 15,306.00 |
| | GC per cent | 41.86 |
| five_prime_UTR | | |
| | Mean length (bp) | 229.85 |
| | GC per cent | 34.44 |
| three_prime_UTR | | |
| | Mean length (bp) | 340.45 |
| | GC per cent | 32.50 |
| **Non protein coding genes** | | |
| **Number of non-protein coding genes** | **3,129** | |
| | Mean ncRNA gene length (bp) | 640.55 |
| | Min length (bp) | 23 |
| | Max length (bp) | 14,112 |
| | GC percent | 41.57 |
| | Pecent ncRNA genes with introns | - |
| | Mean exon number per ncRNA gene | 1.00 |
| **Intergenic (inter protein-coding genes)** | | |
| | Mean length | 7,695 |
| | GC per cent | 32.68 |

##### 1.7 Evaluation of AMIGA heterogeneity

In order to evaluate the heterogeneity of cv. AMIGA, a bulk of 90 AMIGA plants was resequenced using Illumina HiSeq300, with paired-end 2x150 bp reads at Get-PlaGe core facility (INRA, Toulouse, France). This produced 193,734,276 clean reads corresponding to a total of 64.47x depth.

Cutadapt[23] has been used to remove Illumina Truseq adapter from the sequencing data, and to remove bases with a quality score lower than 30, in both 5' and 3' end of the reads. Reads with a length lower than 35 have been discarded. We used BWA-MEM version 0.7.17[24] to map the resequencing reads to the white lupin reference genome. Picard tools (https://github.com/broadinstitute/picard/issues) have been used to detect and remove PCR and Optical duplicates. We then used GATK 4.0[25] HaplotypeCaller tool to call variants. This identified *ca*. 300,000 SNPs without filtering the data. All the SNPs are evenly distributed on the 25 chromosomes and contigs. We generated a VCF file with this information, available in the white lupin genome browser.

##### 1.8 White lupin genome portal

All information regarding white lupin genome sequence is accessible to the scientific community on the white lupin genome portal: www.whitelupin.fr. This website contains a Genome Browser, Expression tools, a Sequence retriever tool as well as all raw data available for download.

### 2 Transposable elements and annotation of repeated sequences

#### 2.1 Material and Methods

##### *2.1.1 Plant material for cytological analysis*

Seeds of *L. albus* cv. AMIGA were germinated on petri dishes with moist paper. After germination root tips were collected. For meiotic tissue, young flower buds were collected from the same variety grown in growth chambers under controlled conditions (16 h light/8 h dark, 25°C day/20°C night, 65% relative humidity and PAR intensity 200 µmol $m^{-2}$ $s^{-1}$) at B&PMP Institute facilities, Montpellier, France. Pre-treatment and fixation of tissues were performed as described below.

##### *2.1.2 Chromosome preparation for in situ hybridization*

Chromosome preparations for *in situ* hybridization analysis were conducted as described in [26] with modifications. First, young roots (pre-treated with 8-hydroxyquinoline 2mM for 3-5 h at room temperature) and anthers were fixed in 3:1 (ethanol:acetic acid) for 2-24 h. The fixed tissues were treated with an enzyme mixture (0.7% cellulase R10, 0.7% cellulase, 1.0% pectolyase, and 1.0% cytohelicase in 1× citric buffer) for 1h at 37 °C. Material was then washed twice in water and fragmented in 7 µl of 60% freshly prepared acetic acid into smaller pieces with the help of a needle on a slide. Another 7 µl of 60% acetic acid was added, and the specimen was kept for 2 min at room temperature. Next, a homogenization step was performed with an additional 7 µl 60% acetic acid and the slide was placed on a 55 °C hot plate for 2 min. The material was spread by hovering a needle over the drop without touching the hot slide.

After spreading of cells, the drop was surrounded by 200 µl of ice-cold, freshly prepared 3:1 (ethanol:acetic acid) fixative. More fixative was added and the slide was briefly washed in fixative, then dipped in 60% acetic acid for 10 min and dehydrated in 96% ethanol. The slides were stored until use in 96% ethanol at 4 °C.

##### *2.1.3 Repeat identification and genome-wide repeat annotation*

Identification and characterization of moderately to highly repeated genomic sequences was achieved by graph-based clustering of genomic Illumina reads using RepeatExplorer2 pipeline [27]. A total of 1,144,690 of 150bp paired reads, representing approx. 0.5× genome coverage, were used for the clustering and the 145 largest clusters with genome proportions of at least 0.01% were examined in detail. Clusters containing satellite DNA (satDNA) repeats were identified based on the presence of tandem sub-repeats within their read or assembled contig sequences with TAREAN [28]. Genome-wide TE repeat annotation was performed using the DANTE (Domain-based ANnotation of Transposable Elements) tool[28]. Consensus sequences of satDNA repeats (Extended Data Fig. 2) and rDNA genes were used to perform genome-wide annotation of satDNA and rDNA arrays using the Geneious v. 9.1.8 annotation tool (https://www.geneious.com). The generated GFF3 files were further incorporated on the *L. albus* genome browser.

##### *2.1.4 Probe preparation and fluorescence in situ hybridization.*

FISH probes were obtained as 5′-Cy3 or 5′-FAM-labeled oligonucleotides (Eurofins MWG Operon, http://www.eurofinsdna.com), or were PCR-amplified as described below. All DNA probes, except oligonucleotides, were labelled with Cy3- or Alexa 488-dUTP (Jena Bioscience) by nick translation, as described by [29] . The sequences of all oligonucleotides and primers are listed in Supplementary Note Table 4. FISH was performed as described in [26]. Probes were then mixed with the hybridization mixture (50% formamide and 20% dextran sulfate in 2× SSC), dropped onto slides, covered with a cover slip and sealed. After denaturation on a heating plate at 80°C for 3 min, slides were hybridized at 37 °C overnight. Post-hybridization washing was performed in 2× SSC for 20 min at 58°C. After dehydration in an ethanol series, 4′,6–diamidino-2–phenylindole (DAPI) in Vectashield (Vector Laboratories, http://www.vectorlabs.com) was applied. Microscopic images were recorded using a Zeiss Axiovert 200M microscope equipped with a Zeiss AxioCam CCD. Images were analyzed using the ZEN software (Carl Zeiss GmbH).

**Supplementary Note Table 4. List of primers and direct-labeled oligo-probes for white lupin studied repeats.**

| Primer name | Sequence | Obs.: |
|---|---|---|
| CL1-170bp-F1 | GTGACATATGCATTTGGGGTAG | |
| CL1-170bp-R1 | ACGTCCAATTCTGATATACGGAA | |
| CL10-78bp-F1 | AAAASTAGCCACAAARAAGT | Centromeric DNA |
| CL10-78bp-R1 | MAGCACAAATAACTCCCAA | Centromeric DNA |
| INT-Tekay-SC1-F1 | CAGAAAGCCAAGATTGAGCATC | |
| INT-Tekay-SC1-R1 | CCCAATATCAAGTTCTCCCCAG | |
| RT-Tekay-SC1-F1 | GTGCGCCTGTTTTGTTGGTT | |
| RT-Tekay-SC1-R1 | TGTTCCTCTCGGCTTTTGGAG | |
| INT-CRM-SC5-F1 | CATGGGCTTTACACGCCCTT | Centromeric DNA |
| INT-CRM-SC5-R1 | GTAAGTCCAGCGGGGTCAAA | Centromeric DNA |
| RT-CRM-SC5-F1 | CAGAGCATGAGCCCTTGTG | Centromeric DNA |
| RT-CRM-SC5-R1 | TGATCCACGCAAAAGGAACA | Centromeric DNA |
| RT-CRM-SC5-F2 | CAACAGAGCATGAGCCCTTG | Centromeric DNA |
| RT-CRM-SC5-R2 | TGACATGATCCACGCAAAAGG | Centromeric DNA |
| RT-SIRE-SC3-F1 | ATTGAGGATGTGCACTGGGG | |
| RT-SIRE-SC3-R1 | CCAAGCCTCTTTCGAACCCT | |
| INT-SIRE-SC3-F1 | CGCGTTGCACACATCCATAT | |
| INT-SIRE-SC3-R1 | TTTGTTGTGGAGTGCGAGGA | |
| **Oligo-probe name** | **Sequence and fluorescent dye** | |
| CL1-sat170bp | [Cy3]GTATATCMGAATTGGACGTRTGACATATGM | |
| CL10-sat78bp | [Cy3]MAAATWACTCCCAAACTTSTTTGTGGC | Centromeric DNA |
| CL21-sat38bp | [FAM]GAAAAGTAGCCAAACAAACAAAAG | Centromeric DNA |
| CL2-microsat5bp | [Cy3]GGATAGGATAGGATAGGATAGGATA | Centromeric DNA |
| CL55-sat8bp | [Cy3]CTAAACTCCTAAACTCCTAAACTCCT | Centromeric DNA |

##### *2.1.5 PCR amplification of tandem repeat and retroelement fragments for probe labeling*

Fragments for probe labeling were amplified using gDNA from *L. albus* using the forward and reverse primers as supplied on Supplementary Note Table 4. Eight PCR reactions for each target repeat were performed in 50 µL reaction volume containing 100 ng of gDNA, 1 µM primers, 1× PCR buffer, 0.2 mM dNTPs, and 1U of Taq polymerase (Qiagen). Thirty-five amplification cycles with proper conditions for each set of primers were run. PCR reactions were sampled, purified and concentrated using Wizard® SV Gel and PCR Clean-Up System (Promega). Correct amplification of PCR fragments was confirmed by Sanger sequencing. After confirmation, the PCR products containing the same class of repeat were collected and used for probe labeling by nick translation as described above.

##### *2.1.6 Identification and generation of LalbCENH3 antibody*

The *CENH3* gene was identified *in silico* by BLAST search from the transcriptome data of *L. albus*. The peptide GRVKHTPASLGGPHRRKAKS was used to generate *L. albus* CENH3-specific (*LalbCENH3*) polyclonal antibodies. Peptide synthesis, immunization of rabbits, and peptide affinity purification of antisera were performed by LifeTein (http://www.lifetein.com).

##### *2.1.7 Immunostaining*

Immunostaining for CENH3 (Abcam-ab1012) were performed as described before [30] with some modifications. Roots were treated with 8-hydroxyquinoline 2mM for 3-5 h at room temperature and fixed in ice-cold 4% paraformaldehyde in 1×PBS buffer pH 7.5 (1.3 M NaCl, 70 mM $Na_2HPO_4$, 30 mM $NaH_2PO_4$) for 1 h and squashed in a drop of the same buffer. Then, the slides were washed in 1×PBS and blocked with 3% BSA for 30 min at 37 °C. The antibodies used were rabbit anti-LalbCENH3 diluted 1:500 in 1% BSA in 1xPBS. The detection of anti-LalbCENH3 and anti-H3K4me3 were done with goat anti-rabbit CF488 (Sigma, SAB4600389), or alternatively goat anti-rabbit Alexa-594 (JacksonImmunoResearch-111-585-008), and donkey anti-mouse Fluorescein (JacksonImmunoResearch-715-097-003), respectively, diluted 1:200 in 1×PBS containing 1% BSA. The slides were counterstained with 2 μg/ml 4',6-diamidino-2-phenylindole (DAPI) in Vectashield H-1000.

##### *2.1.8 Super-resolution microscopy*

To analyze the structures and spatial arrangement of FISH signals and the global chromatin at a lateral optical resolution of ~120 nm (super-resolution, achieved with a 488 nm laser), 3D structured illumination microscopy (3D-SIM) was applied using a C-Apo 63×/1.2W Korr objective of an Elyra PS.1 microscope system and the software ZEN (Carl Zeiss GmbH). Image stacks were captured separately for each fluorochrome using the 561, 488, and 405 nm laser lines for excitation and appropriate emission filters, and then merged using the ZEN software[31] .

##### 2.1.9 CENH3-ChIP and ChIP-seq analysis

Chromatin immunoprecipitation experiments were done with Abcam ChIP Kit - Plants (ab117137) following the manufacturers instructions. First, 1 g of young *L. albus* cv. AMIGA leaves were collected and cross-linked with formaldehyde 1% for 15 min on ice. Leaves were then ground in liquid nitrogen and sonicated using a Diagenode Sonicator. Sonicated chromatin-DNA ranging from 200-1000 bp was immunoprecipitated using anti-LalbCENH3. Immunoprecipitated DNA and, as control, Input chromatin DNA samples (3-7ng for each sample) were sent for ChIPseq at BGI. The original ChIPseq sample data are available at White Lupin Genome Website (http://www.whitelupin.fr)

To identify repeats associated with CENH3-containing chromatin, reads from the ChIPseq experiment obtained by sequencing DNA from isolated chromatin prior to (the input control sample) and after immunoprecipitation with the CENH3 antibody (the ChIP sample) were separately mapped to the repeat clusters. The mapping was based on read similarities to contigs representing individual clusters, using BLASTn (22) with parameters "-m 8 -b 1 -e 1e-20 -W 9 -r 2 -q -3 -G 5 -E 2 -F F" and custom Perl scripts for parsing the results. Each read was mapped to a maximum of one cluster, based on its best similarity detected among the contigs. Ratio of ChIP/input reads assigned to individual clusters was then used to identify repeats enriched in the ChIP sample as compared to the input.

#### 2.2 Results

Tandem repeat-rich regions create a technical challenge to genome assembly26. By means of in silico analysis with RepeatExplorer2, we identified 14 tandem repeat clusters (CL1, CL2, CL10, CL21, CL52, CL53, CL55, CL68, CL77, CL85, CL114, CL118, CL121, CL127) accounting for ~15% of the genome (Supplementary Note Table 5 & 6; Fig. 1a). These tandem-repeats varied from short monomer length with

5 bp consensus sequence (CL2) to very long monomers up to 918bp (CL121). After further characterization of the consensus sequence of the identified tandem repeats we observed that CL52 and CL127 are very similar to CL1 and was grouped within a single supercluster, thus representing a subset of the same repeat comprising the most abundant tandem repeat family of WL. CL10 (78bp) and CL21 (38bp) are similar satellites comprising different subfamilies. CL10 represents a dimer organization of CL21. CL118 (182bp) also showed sequence similarity to CL10 and CL21. Among the identified tandem repeats CL1-CL52-CL127 (170bp) and CL2 (5bp) were highly abundant comprising both together ~12% of the genome (Supplementary Note Table 5). Further characterization of satellite sequences is provided on Supplementary Note Table 6.

**Supplementary Note Table 5. Detailed repetitive DNA composition of the *Lupinus albus* genome**

<table>
<tr><th colspan="3">Class</th><th>Genomic abundance [%]</th></tr>
<tr><td colspan="2" rowspan="6">LTR retroelements Ty1/copia</td><td>Ale</td><td>0.18</td></tr>
<tr><td>Ikeros</td><td>1.21</td></tr>
<tr><td>Ivana</td><td>0.20</td></tr>
<tr><td>SIRE</td><td>6.26</td></tr>
<tr><td>TAR</td><td>0.43</td></tr>
<tr><td>Tork</td><td>0.96</td></tr>
<tr><td rowspan="4">LTR retroelements Ty3/gypsy</td><td rowspan="2">non-chromovirus</td><td>Athila</td><td>1.24</td></tr>
<tr><td>Ogre</td><td>0.27</td></tr>
<tr><td rowspan="2">chromovirus</td><td>CRM</td><td>3.40</td></tr>
<tr><td>Tekay</td><td>16.68</td></tr>
<tr><td colspan="2" rowspan="4">DNA transposons</td><td>EnSpm CACTA</td><td>0.34</td></tr>
<tr><td>hAT</td><td>0.04</td></tr>
<tr><td>MuDR Mutator</td><td>0.27</td></tr>
<tr><td>Helitron</td><td>0.11</td></tr>
<tr><td colspan="3">Other TEs</td><td>11.64</td></tr>
<tr><td colspan="2" rowspan="2">rDNA</td><td>35S rDNA</td><td>2.43</td></tr>
<tr><td>5S rDNA</td><td>0.15</td></tr>
<tr><td colspan="3">Tandem repeats</td><td>14.71</td></tr>
<tr><td colspan="3">Total</td><td>60.52</td></tr>
</table>

**Supplementary Note Table 6.Satellite DNA genomic abundance, monomer length and features.**

| SatDNA clusters | Genomic abundance [%] | Monomer length (bp) | Obs.: |
|---|---|---|---|
| CL1, CL52, CL127 | 6,129 | 170 | Most abundant satDNA; No enrichment with CENH3-immunoprecipitated DNA |
| CL2 | 5,473 | 5 | Relative enrichment with CENH3-immunoprecipitated DNA |
| CL10 | 1,290 | 78 | Enrichment with CENH3-immunoprecipitated DNA |
| CL21 | 0,766 | 38 | Enrichment with CENH3-immunoprecipitated DNA |
| CL53 | 0,303 | 24 | No enrichment with CENH3-immunoprecipitated DNA |
| CL55 | 0,285 | 8 | High enrichment with CENH3-immunoprecipitated DNA |
| CL68 | 0,190 | 7 | Telomere motif |
| CL77 | 0,117 | 36 | No enrichment with CENH3-immunoprecipitated DNA |
| CL85 | 0,085 | 76 | No enrichment with CENH3-immunoprecipitated DNA |
| CL114 | 0,027 | 247 | No enrichment with CENH3-immunoprecipitated DNA |
| CL118 | 0,024 | 182 | Similarity to CL10 and CL21; No enrichment with CENH3-immunoprecipitated DNA |
| CL121 | 0,023 | 918 | No enrichment with CENH3-immunoprecipitated DNA |
| Total | 14,713 | | |

Chromosomal scale genome-wide annotation of repetitive sequences revealed the *in silico* distribution of the main classes of repeats (Fig. 1b). LTR retroelements showed an increased density towards the central regions of chromosome assemblies along presumed pericentromeric regions. A narrow peak for the distribution of CRM (Centromeric Retrotransposon of Maize) clade retroelements was observed in all assembled chromosomes. Giving the known association of CRM clade with centromeres in different plant species we presumed that the observed peak defines the centromeric regions of WL chromosomes. Thus, we refer to this element as CRWL for Centromeric Retrotransposon of While Lupin. Remarkably, a high association of CRWL and satellite DNA peaks was observed, suggesting a more specific distribution of these repeats compared to other TEs.

To further characterize the repeat portion of WL genome we performed *in situ* hybridization of the most abundant repeat clusters identified by the RepeatExplorer analysis. Given the abundance of class I TEs and diversity of tandem repeats, we studied the *in situ* distribution of the three most abundant classes of LTRs found, the Ty1/Copia SIRE and of the Ty3/Gypsy chromoviruses Tekay and CRWL, as well as the main tandem repeats (CL1-170bp, CL2-5bp, CL10-78bp, CL21-38bp and CL55-8bp). As expected CRWL FISH signals were observed as narrow distributed signals at the centromeric regions of most WL chromosomes. Remarkably, a precise co-localization of signals of CRWL and CL2-5bp satDNA was observed (Fig. 1b-1). A more wide distribution towards the pericentromere of WL chromosomes was observed for the other two retrotransposons studied, SIRE and Tekay. FISH signals of CL1-170bp repeat were observed in 8 chromosome pairs, mostly seen as large blocks apparently located in pericentromeric heterochromatin knobs (Extended Data Fig. 2); CL10-78bp repeat-signals were observed in 14 chromosome pairs with a narrow distribution in centromeric regions; CL21-38bp and CL55-8bp repeat-signals were also observed as narrow centromeric signals in three different chromosome pairs each. FISH mapping of the main repeats are presented on Fig. 1c and Extended Data Fig. 2.

The association of CRWL with functional centromeres was analyzed by CENH3-ChIPseq. We have then characterized the CENH3 gene and developed a specific antibody which is able to recognize all WL centromeres (Supplementary Fig. 1). Analysis of ChIPseq reads demonstrated that CRWL elements (CL13, CL20, CL34, CL48 and CL49) are among the clusters that showed the highest levels of association in the immunoprecipitated fraction (Fig. 1d). Although CRWL is highly abundant on centromeric regions of WL chromosomes (Fig. 1b-c), detailed analysis of ChIPseq data revealed that CENH3-containing chromatin is also associated with at least four families of centromeric tandem repeats: CL2-5bp, CL10-78bp, CL21-38bp and CL55-8bp (Fig. 1c-d). In contrast, the most abundant satellite repeat CL1-170bp did not show significant enrichment with the immunoprecipitated DNA. Remarkably, the CL55-8bp satellite, which is specifically located in three chromosome pairs, showed the highest ratio of enrichment (Fig. 1c-d). Indeed super-resolution microscopy of pachytene and somatic chromosomes confirmed a centromere-specific localization for CRWL, CL2, CL10, CL21 and CL55 repeats, while CL1 repeat localizes aside core centromeres (Fig. 1c, Extended Data Fig. 2, arrowheads). Summing up the total amount of repeat classes showing enrichment with CENH3-immunoprecipitated chromatin we have about 11% (49.55 Mb) of the genome. Thus, the functional centromeres of WL are preferentially associated with CRWL and with different families of tandem repeats in a chromosome-specific pattern. Our results confirm a high centromeric DNA diversity for WL centromeres.

### 3 White lupin diversity analysis

We selected 14 white lupin accessions, covering a wide range of diverse material, to evaluate a broader range of the genetic diversity and determine population structure and linkage disequilibrium. The 14 accessions selected for this work, together with the variety AMIGA - used to construct the white lupin reference genome - are described in the Supplementary Note Table 7. This group of accession comprises cultivated varieties with different pattern of sowing, one landraces and two wild accessions.

**Supplementary Note Table 7. White lupin accessions used in this study.**

| Genotype | Accession type | Sowing Time | Seed color | Country of origin |
|---|---|---|---|---|
| Amiga | Cultivated | Spring | White | Chile |
| Clovis | Cultivated | Winter | White | France |
| Dieta | Cultivated | Spring | White | Ukraine |
| Energy | Cultivated | Spring | White | France |
| Feodora | Cultivated | Spring | White | Germany |
| Figaro | Cultivated | Spring | White | France |
| GR38 (Megalopolis) | Non-domesticated | Winter | Marble | Greece |
| GRAECUS | Non-domesticated | Winter | Dark Marble | Greece |
| Kiev | Cultivated | Spring | White | Ukraine |
| Lucky | Cultivated | Spring | White | France |
| Luxe | Cultivated | Winter | White | France |
| Magnus | Cultivated | Winter | White | France |
| Orus | Cultivated | Winter | White | France |
| P27174 | Landrace | Winter | White | Ethiopia |
| Ulysse | Cultivated | Winter | Pink | France |

#### 3.1 Methods

##### *3.1.1 Data generation with short-reads technology*

Young leaves of 30 plants were used to extract genomic DNA of each accession using the QIAGEN Genomic-tip 100/G kit following the supplier's recommendations. The accessions were sequenced using Illumina technology using paired-end 2x150 bp short-reads. It was generated a total of 310.95 Gb of data with average sequencing depth of 45.99x (Supplementary Note Table 8).

**Supplementary Note Table 8. Summary of re-sequencing of white lupin accession using short reads.**

| Acession | Number of reads | Sequencing depth (x) |
|---|---|---|
| Clovis | 85072762 | 28.31 |
| Dieta | 88382474 | 29.41 |
| Energy | 120147660 | 39.98 |
| Feodora | 111688474 | 37.17 |
| Figaro | 88274518 | 29.38 |
| GR38 | 123360898 | 41.05 |
| GRAECUS | 148837522 | 49.53 |
| Kiev | 247673360 | 82.42 |
| Lucky | 107121722 | 35.65 |
| Luxe | 89069200 | 29.64 |
| Magnus | 216658716 | 72.10 |
| Orus | 93626420 | 31.16 |
| P27174 | 256870228 | 85.48 |
| Ulysse | 102510626 | 34.11 |

##### *3.1.2 Mapping and SNP detection*

Cutadapt[23] has been used to remove Illumina Truseq adapter from the sequencing data, and to remove bases with a quality score lower than 30, in both 5' and 3' end of the reads. Reads with a length lower than 35 have been discarded. We then used BWA-MEM version 0.7.17[24] to map the resequencing reads from all 15 genotypes to the white lupin reference genome. PCR and Optical duplicates have been detected and removed using Picard Tools. After that, GATK 4 HaplotypeCaller tool have been used in emit-ref-confidence GVCF mode to produce one gvcf file per sample. These files have been merged using GATK CombineGVCFs. Finaly, GATK GenotypeGVCFs have been used to produce a vcf file containing variants from all the 15 samples. This identified a total of 6,620,353 SNPs/indel. After filtering for minimum allele frequency of 0.15 and heterozygosity frequency of 0–0.2, 2,659,837 SNPs were retained to further analysis. All the SNPs are evenly distributed on the 25 chromosomes with a mean of SNP density of 10.74 SNPs per Kb of sequence (Fig. 2A, Supplementary Note Table 9).

**Supplementary Note Table 9. Summary of SNP distribution along the 25 white lupin chromosomes.**

| Chromosome | Chr. size (Mb) | SNPs numbers | SNP density (per Kb) |
|---|---|---|---|
| Lalb_Chr01 | 23.521 | 251648 | 10.70 |
| Lalb_Chr02 | 17.964 | 189793 | 10.57 |
| Lalb_Chr03 | 21.293 | 209454 | 9.84 |
| Lalb_Chr04 | 16.824 | 178403 | 10.60 |
| Lalb_Chr05 | 17.322 | 166236 | 9.60 |
| Lalb_Chr06 | 17.860 | 180397 | 10.10 |
| Lalb_Chr07 | 17.716 | 173366 | 9.79 |
| Lalb_Chr08 | 16.982 | 195090 | 11.49 |
| Lalb_Chr09 | 15.103 | 178221 | 11.80 |
| Lalb_Chr10 | 19.347 | 202619 | 10.47 |
| Lalb_Chr11 | 20.058 | 178485 | 8.90 |
| Lalb_Chr12 | 17.349 | 206723 | 11.92 |
| Lalb_Chr13 | 16.178 | 185099 | 11.44 |
| Lalb_Chr14 | 14.546 | 168640 | 11.59 |
| Lalb_Chr15 | 20.045 | 236105 | 11.78 |
| Lalb_Chr16 | 14.402 | 152366 | 10.58 |
| Lalb_Chr17 | 15.015 | 160319 | 10.68 |
| Lalb_Chr18 | 20.222 | 168727 | 8.34 |
| Lalb_Chr19 | 18.232 | 189237 | 10.38 |
| Lalb_Chr20 | 18.249 | 217106 | 11.90 |
| Lalb_Chr21 | 15.132 | 149269 | 9.86 |
| Lalb_Chr22 | 15.014 | 182937 | 12.18 |
| Lalb_Chr23 | 16.633 | 205286 | 12.34 |
| Lalb_Chr24 | 12.487 | 121927 | 9.76 |
| Lalb_Chr25 | 16.406 | 195986 | 11.95 |

###### *3.1.3 Phylogenetic analysis, population structure and linkage disequilibrium*

The genetic distance matrix was calculated based on identity-by-state similarity method and an average cladogram constructed using neighbour-joining algorithm implemented on TASSEL 5.2.51 [32]. Then, a phylogenetic tree was prepared using the iTOL v 4.3 [33]. A principal component analysis (PCA) was also performed in R [34] (http://www.R-project.org/) function "prcomp" .

A Bayesian model-based clustering method implemented with STRUCTURE v2.3.4 [35,36] was used to investigate the population structure using all the filtered SNPs. The program was run 10 times for each K value, ranging from 1 to 5, with a 1,000 burn-in time and 1,000 iterations. The optimal K value was determined based on the ΔK from the Structure Harvester v0.6.94 [37] program, through Evanno's test [38].

Linkage Disequilibrium (LD) was estimated with the $r^2$ statistics using the R package snpStats [39] (version 1.32.0) using a subset comprising 46,783 SNPs, distributed evenly on all chromosomes (average of 1871 SNPs per chromosome). Heatmaps of pairwise LD between markers were plotted using the R package LDheatmap (version 0.99-5) [40].

#### 3.2 Results

###### *3.2.1 Results*

The genetic distance ranged from 0.103 to 0.409, with an average of 0.296 (Supplementary Table 2). The smallest genetic distance (0.103) was observed between accessions AMIGA and FEODORA, both of which are cultivars from the French breeding program.

A neighbor-joining phylogenetic tree consisting of 15 white lupin accessions was constructed based on the 4,643,439 high-quality SNPs (Fig. 2b). The tested white lupin accessions were clearly divided into three clusters, reflecting their domestication pattern: accessions adapted for sowing in spring (vernalization needed, slow growth, cold adapted), accessions with sowing during before winter (no vernalization needed, fast growth, strong vigor, reduced life-cycle) and non-domesticated varieties, which included wild varieties and landraces.

To further verify the clustering observed in the phylogenetic tree, a principal component analysis (PCA) was conducted using the same samples and SNP set. More than half of total genetic variance (58.2%) could be explained by the two first components (Supplementary Fig. 2a). The scatter plots of these first two principal components (Supplementary Fig. 2b) clearly revealed that the accessions in the three clusters of the phylogenetic tree were grouped separately. It also highlighted that the accession LUXE is in an overlap of the winter accessions group and the non-domesticated accessions group.

The population structure was explored with the same set of SNPs using STRUCTURE[35,36]. We tested for a population structure ranging from 2 subpopulations (K=2) up to 5 subpopulations (K=5, Extended Data Fig. 4d). Additionally, we applied Evanno's test [38] to determine the optimal K value according to the maximum ΔK value. The highest ΔK value (68,117) was observed when K = 2. This result indicated that these 15 white lupin accessions might be divided into two subpopulations, one formed by the spring accessions and the other with the winter and non-domesticated accessions.

We estimated the level of LD using the $r^2$ parameter between all pairwise SNP comparisons by using these 15 accessions in the 25 chromosomes (Supplementary Fig. 3). We used a subset of the high-quality filtered SNPs, containing 46,783 markers. LD decay distance showed apparent variation, indicating that selection pressure in different chromosomal regions was varied, probably due to different selection goals in breeding.

#### 3.3 Long-read sequencing and *de novo* assembly of GRAECUS and P27174

##### *3.3.1 Methods*

We selected the wild accession GRAECUS and the landrace P27174 to a further investigation of the impact of domestication on the white lupin genome. Genomic DNA of both accessions was extracted as described on the section 3.1. Long-read sequencing was realized using Oxford Nanopore technology, using a GridION 18.04.1-0, with a software Minknow 1.10.24-1 at platform at Get-PlaGe core facility (INRA, Toulouse, France). Briefly, DNA was used to prepare a library with the

ONT It was used the Ligation Sequencing Kit 1D (sqk-lsk109). The DNA was sequenced using a single ONT MinION R9.4 flowcell (FLO-MIN106) for 48h. Base-calling was performed using Albacore 2.1.10-1. This produced 1,280,206 sequences for GRAECUS, corresponding to 12.45 Gb of data with a N50 length of 13.6 Kb (27.6 x of sequencing depth, Supplementary Fig. 4a). For the accession P27174 this produced a total of 1,738,579 reads corresponding to 14.59 Gb of data with N50 length of 11.8 Kb (32.36 x of sequencing depth, Supplementary Fig. 4b).

The *de novo* assembly of the two genotypes were performed using CANU[1]. For P27174-4, two round of correction have been made prior to the assembly step, using the parameters correctedErrorRate=0.16 anc corMaxEvidenceErate=0.15. For Graecus, only one round of correction have been made, using minOverlapLength=400, correctedErrorRate=0.16 and corMaxEvidenceErate=0.15. The Illumina paired-end data described in 3.1 were used to polish two times the two genome assemblies using Pilon[8]. BUSCO v 3.0.0[17] was run on the set of predicted transcripts. The assessment software detected for GRAECUS 96.8% of complete gene models (1,131 complete single copy and 200 duplicated respectively) plus 9 additional fragmented gene models. For P27174 97.8% of complete gene models (1,125 complete single copy and 220 duplicated respectively) plus 4 additional fragmented gene models. Structural variation of these two accession were performed in Assemblytics software [41] based on whole genomes alignments generated with MUMmer [42]. The final assembly of GRAECUS has 428.06 Mb and P27174 458.23 Mb (Supplementary note table 10).

**Supplementary Note Table 10. Statistics of GRAECUS and P27174 *de novo* genome assembly.**

| **Data acquisition** | | | |
|---|---|---|---|
| | Total number of Reads | Total sequencing length (Gb) | N50 reads length (Kb) |
| GRAECUS | 1 280 690 | 12.45 | 13.6 |
| P27174 | 1 739 129 | 14.59 | 11.8 |

| **Assembly Features** | | | |
|---|---|---|---|
| | Total number of Contigs | Total size assembled (Mb) | N50 (Kb) |
| GRAECUS | 5385 | 428.06 | 251.63 |
| P27174 | 5869 | 458.23 | 207.64 |

###### *3.3.2 Results*

Analysis of genome structural variation

In order to detect existing structural variation (SVs) between two *de novo* assembled genotypes and the reference genome AMIGA we used Assemblytics software [41] based on whole genomes alignments generated with MUMmer [42]. This software classified SVs into insertions, deletions, tandem expansions, tandem contractions, repeat contraction, and repeat expansion. P27174 assembly has a total length of 18.67 Mb of SVs affected whereas GRAECUS has a slightly smaller size of the genome affected by SVs (18.08 MB – Extended Fig. 4a, Supplementary Table 3).

The repeat expansion and contraction are the majority of SVs sequences in GRAECUS, accounting for 53% of the total SVs and 12.3 Mb of bases affected. The following most abundant SVs are the insertion and deletion in the genome that accounts for 3.33 Mb of SVs. In P27174 the majority of variants are the indels and they affect a smaller portion of the genome compared to GRAECUS (11.44 Mb). P27174 presented more tandem SVs (expansion and contraction) than GRAECUS, either in number of variants and the size total size of variants.

### 4 Seed Quality

#### 4.1 Identification and characterization of seed storage protein

###### *4.1.1 Methods*

White Lupin protein extracts and electrophoresis

Total soluble proteins were extracted from mature seeds of AMIGA, P27174 and GRAECUS using 500 µl of the Tris-HCl/SDS lysis buffer for 10 mg of seed powder (two independent extractions, A and B, realized). For AMIGA, seeds were splitted in two samples based on seed color (dark brown, DB; light brown, LB). The protein extracts were stirred for 30 min at room temperature and then centrifuged (35,000g) for 10 min. The supernatant was subjected to a second clarifying centrifugation step as above. Protein concentration was determined using the Protein assay from Bio-

Rad (Marnes-la-Coquette, France). For each seed sample, 10 µg proteins were separated by one-dimensional electrophoresis (1D) in a SDS polyacrylamide gel (SDS-PAGE) using the XCell4 *Surelock*™ Midi-Cell system (Life Technology) [Resolution gel: 4.1 mM Tris-HCl pH 8.8, 40% (v/v) acrylamide/bisacrylamide (30%/0.8%), 1% (g/v) SDS, 0.05% (g/v) ammonium persulfate, 0.05% (v/v) TEMED; Concentration gel: 0.6 mM Tris-HCl pH 6.8, 13% (v/v) acrylamide/bisacrylamide (30%/0.8%), 1% SDS, 0.05% ammonium persulfate, 0.05% TEMED; Electrophoresis buffer: 50 mM Trizma base, 380 mM glycine, 0.1% (g/v) SDS]. Gels were stained with Coomassie Blue R250 (Bio-Rad) and scanned using an Odyssey Infrared Imaging System (LI COR BIOSCIENCES GMBH, Bad Homburg, Germany), with an intensity of 7.5 and a resolution of 84 µm. Protein band detection and quantification were performed using Phoretix 1D (v11.2, Total Lab Limited, Newcastle, UK). The quantitative data of each band was normalized by dividing its volume by total band volume in the well, and the molecular weight (kDa) of each band was calculated using a low-range protein ladder (Bio-Rad).

Mass spectrometry analysis

Gel pieces were washed in water: acetonitrile solution (1:1, 5 min) followed by 100% acetonitrile (10 min). Reduction and cysteine alkylation was performed by successive incubation with 10 mM dithiothreitol in 50 mM $NH_4HCO_3$ (30 min, 56 °C), then 55 mM iodoacetamide in 50 mM $NH_4HCO_3$ (20 min, RT, in dark). Pieces were then incubated with 50 mM NH4HCO3 and acetonitrile (1:1, 10 min) followed by acetonitrile (15 min). Proteolytic digestion was carried out overnight using 25 mM $NH_4HCO_3$ with 12.5 ng/µl Trypsin (Sequencing grade, Roche diagnostics, Paris, France). Resultant peptides were extracted by incubation in 5% formic acid (sonicated) with the supernatant removed and saved, followed by incubation in acetonitrile and 1% formic acid (1:1, 10 min) and a final incubation with acetonitrile (5 min), again supernatant was removed and saved. These two peptide extractions were pooled and dried using a SPD1010 speedvac system (Thermosavant, Thermofisher Scientific, Bremen, Germany) and the resultant peptide mixture was analyzed by nanoflow liquid chromatography tandem mass spectrometry (Nano LC-MS/MS).

All experiments were performed on a dual linear ion trap Fourier Transform Mass Spectrometer (FT-MS) LTQ Orbitrap Velos (Thermo Fisher Scientific, Bremen, Germany) coupled to an Ultimate® 3000 RSLC Ultra High Pressure Liquid Chromatographer (Dionex, Amsterdam, The Netherlands) controlled by Chromeleon Software (version 6.8 SR11; Dionex, Amsterdam, The Netherlands).

Samples were desalted and concentrated for 10 min at 5μL/min on an LCPackings trap column (Acclaim PepMap 100 C18, 75 μm inner diameter x 2cm long, 3μm particles, 100Å pores).

The peptide separation was conducted using a LCPackings nano-column (Acclaim PepMap C18, 75μm inner diameter x 50cm long, 2μm particles, 100Å pores) at 300nL/min.

Gradient profile:

- Columns equilibration with 96% solvent A (0.1% formic acid, 97.9 % water, 2% acetonitrile (v/v/v)) and 4% solvent B (0.1% formic acid, 15.9 % water, 84% acetonitrile (v/v/v)).
- Gradient of 4-60% solvent B for 90min.
- Stage at 99% solvent B for 10min.

Data were acquired in positive mode in data-dependent mode to automatically switch between high resolution full-scan MS spectra (R 60 000) collected in profile mode and low resolution CID-MS/MS in centroid mode (m/z 300-1800). The 20 most intense peptide ions with charge states ≥ 2 were sequentially isolated and fragmented in the high pressure linear ion trap by low energy CID (collision energy 35%, activation time 10ms, Qz 0.25). Dynamic exclusion is activated during 30 seconds with a repeat count of 1.

For protein identification, MS/MS ion searches were performed using Mascot search engine version 2.3.2 (Matrix Science, London, UK) via Proteome Discoverer 2.1 software (ThermoFisher Scientific, Bremen, Germany) against *L .albus*_protein database and NCBI prot_viridiplantae. The search parameters included trypsin as a protease with two allowed missed cleavages and carbamidomethylcysteine, methionine oxidation and acetylation of N-term protein as variable modifications. The tolerance of the ions was set to 5 ppm for parent and 0.8 Da for-fragment ion matches. Peptides and proteins identified by MASCOT were validated using « Peptid

Prophet » and « Protein Prophet » algorithm with Scaffold software (version 4.8.7, Proteome Software, Portland, USA). Protein identifications were accepted if they contained at least two identified peptide.

Identification of conglutin protein genes in the *L. albus* genome and phylogenetic relationship

Conglutin cDNA sequences of *L. angustifolius* described previously [45] were used to identify orthologs on *L. albus* genome using BLAST tool search. The sequences used as query were: Lang_ALPHA1 [Genbank:HQ670406], Lang_ALPHA2 [Genbank:HQ670407], Lang_ALPHA3 [Genbank:HQ670408], Lang_BETA1 [Genbank:HQ670409], Lang_BETA2 [Genbank:HQ670410], Lang_BETA3 [Genbank: HQ670411], Lang_BETA4 [Genbank:HQ670412], Lang_BETA5 [Genbank:HQ670413], Lang_BETA6 [Genbank:HQ670414], Lang_BETA7 [Genbank:HQ670415], Lang_GAMMA1 [Genbank: HQ670416], Lang_GAMMA2 [Genbank:HQ670417], Lang_DELTA1 [Genbank: HQ670418], Lang_DELTA2 [Genbank:HQ670419], Lang_DELTA3 [Genbank:HQ670420] and Lang_DELTA4 [Genbank:HQ670421].

A phylogenetic tree with all the *L. angustifolius* conglutin aminoacid sequences and the amino acid ortholog sequences identified on *L. albus* genome were constructed using the pipeline on the Phylogeny.fr platform [46]. Sequences were aligned with MUSCLE (v3.7) conFig.d for highest accuracy (MUSCLE with default settings). The phylogenetic tree was reconstructed using the maximum likelihood method implemented in the PhyML program (v3.0). The WAG substitution model was selected assuming an estimated proportion of invariant sites and 4 gamma-distributed rate categories to account for rate heterogeneity across sites. The gamma shape parameter was estimated directly from the data. Reliability for internal branch was assessed using the aLRT test (SH-Like).

###### *4.1.2 Results*

*L. albus* conglutin identification

We identified a total of 15 genes coding for conglutin proteins in the white lupin genome (Supplementary Table 4), a similar number (16) to the one described for the narrow-leafed lupin genome [45]. We identified 3 distinct α-conglutin – also known as 11S globulin – orthologs one for each narrow-leafed lupin α-conglutin. These proteins were already described in the literature [45], and in this work we could localize the genes coding for each one of them that were positioned on the chromosomes 02, 21 and 25 (Supplementary Fig. 5 and Supplementary Table 4). We identified 6 genes that are orthologs of β-conglutin (or 7S globulin), one less than found in *L. angustifolius*. One of them is located on chromosome 22 and the other five are positioned on chromosome 6. The β-conglutin genes on chromosome 6 are tandem duplications and form a cluster of 87.5 kb positioned from 6,684,640 bp to 6,772,112 bp. They showed highest homology among themselves than to β-conglutins of narrow-leafed lupin (Supplementary Fig. 5). The same pattern was shown for the 4 δ-conglutin identified (Supplementary Table 4): they form a paralogs cluster on chromosome 24 and with a high conservation level. We also identified 2 basic 7S proteins, that are specific proteins for lupin seeds [47], that have a high level of homology with the 2 *L. angustifolius* γ-globulin (Supplementary Fig. 10). They are single genes on chromosomes 4 and 15, respectively.

*L. albus* seed storage protein characterization

We analyzed seed protein composition of the three white lupin accessions explored in this study (domesticated, landrace, and wild accessions, Supplementary Extended Fig. 3a-c). The seed protein profile of GRAECUS was highlight contrasted to that of the other accessions. Five protein bands showing striking differences in their abundance between the three accessions were selected for mass spectrometry analyses (Extended Data Fig. 3b, red arrows). With the availability of the full-length sequences of white lupin, the most abundant proteins in these bands were identified as corresponding to β-conglutins from the gene cluster on chromosome 6 (Supplementary Table 5).

It was previously described that β-conglutins are the major component of lupin seed storage protein and account for about 43.4% [47], followed by α-conglutins (33%). β-conglutins are relevant in human nutrition since they have been identified as being hypocholesterolemic [48] and preventing or reducing hypertension [49]. According to Duranti et al. [47], β-conglutins have pronounced structural variation, being composed of over 20 polypeptide chains with no disulfide bridges, with a broad range of molecular masses ($M_r$ 15–65 kDa). This revealed that domestication was associated with the disappearance of high molecular weight storage proteins (~60 kDa, Extended Fig. 3b-c). These corresponded to ß-conglutin precursors that can be proteolytically processed during seed development to give rise to polypeptides of lower molecular weights. The accumulation of these processed forms at the expense of precursor forms in seeds of the domesticated variety AMIGA could be linked to an increased germination capacity. Importantly, the unprocessed ß-conglutins of high molecular weight have high allergenicity[50], a trait that might have been counter-selected during domestication.

#### 4.2 Alkaloid characterization

##### *4.2.1 Methods*

*Tissue sampling for alkaloid analysis*

Seeds of WL accessions. AMIGA, GRAECUS and P27174, were germinated on germination paper during 4 days. Seedlings were cultivated in growth chambers under controlled conditions (16 h light – from 8:00 to 23:59 / 8 h dark – from 00:00 to 7:59, 25°C day/20°C night, 65% relative humidity and PAR intensity 200 µmol $m^{-2}$ $s^{-1}$). After germination, seedlings were transferred to 200-L tanks. The hydroponic solution was modified from previously described medium [12] without phosphate, according to the following composition: $MgSO_4$, 54 µM; $Ca(NO_3)_2$ 400 µM; $K_2SO_4$ 200 µM; Na-Fe-EDTA 10 µM; $H_3BO_3$ 2.4 µM; $MnSO_4$ 0.24 µM; $ZnSO_4$ 0.1 µM; $CuSO_4$ 0.018 µM and $Na_2MoO_4$ 0.03 µM. The nutrient solution was continuously aerated and was renewed every 7 days.

At the 3-leaf stage, young leaf material was collected from each plant at 11:00 h. At 5-leaf stage young leaf, stem, root system and the rest of the shoot material were also collected at 11:00 h. All tissues were placed immediately in liquid nitrogen and stored at –50 °C until further analyses. Each plant represents a biological replicate. There were 4 biological replicates for each genotype.

*Alkaloid quantification*

Frozen tissue samples were pulverized in liquid nitrogen using metal balls and a ball grinder. Around 50 mg of frozen powder were mixed with 500 μl of 60% methanol containing 0.06% formic acid and 5 mg/l caffeine as internal standard. The mixtures were agitated vigorously for 2 h at room temperature. Following a brief centrifugation to separate the solids, the extracts were diluted 1:5 (v:v) with water, cleared through a 0.22-μm filter, and analyzed by reversed-phase LC-MS.

The diluted, cleared extracts were analyzed using a Thermo Fisher Dionex UltiMate 3000 RS HPLC/UHPLC system coupled to a Bruker compact QqTOF mass spectrometer via an ESI source. Analyte separations were performed on a Kinetex XB-C18 column (100 x 2.1 mm, 1.7 μm, 100Å, Phenomenex). The eluent flow rate was 0.3 ml/min and the column temperature was kept constant at 40°C. Mobile phases A and B consisted of 0.05% formic acid in water and 0.05% formic acid in acetonitrile, respectively. The elution profile was 0 – 1 min, 2% B (constant); 1 – 16 min, 2 – 25% B (linear); 16 – 24 min, 25 – 65% B (linear), 24 – 26 min, 65 – 100% B (linear); 26 – 27 min, 100% B (constant); 27 – 27.5 min, 100 – 2 % B (linear); and 27.5 – 33 min, 2% B (constant). ESI mass spectra were acquired in positive ionization mode with the following parameters: capillary voltage of 4500 V; end plate offset of -500 V; source temperature of 250°C; desolvation gas flow of 8.0 l/min; and nebulizer pressure of 2.5 bar. $N_2$ was used as desolvation, nebulizer and collision cell gas.

The identity of lupanine was confirmed by comparison with a standard (Sigma-Aldrich), while that of the other alkaloids was inferred from their exact masses and their ESI(+) fragmentations at different collision energies. For quantification, alkaloid peaks were normalized to the signal of the internal standard and the fresh weight of the samples.

*Expression analysis*

Total RNA from leaves harvested on 3-leaf and 5-leaf stage plants was extracted using the Direct-zol RNA MiniPrep kit (Zymo Research, Irvine, CA) according to the manufacturer's recommendations. RNA concentration was measured on a NanoDrop (ND1000) spectrophotometer. Poly(dT) cDNA were prepared from 1.5 µg total RNA using the revertaid First Strand cDNA Synthesis (Thermo Fisher). Gene expression was measured by quantitative Real Time - Polymerase Chain Reaction (qRT-PCR) (LightCycler 480, Roche Diagnostics, Basel, Switzerland) using the SYBR Premix Ex Taq (Tli RNaseH, Takara, Clontech, Mountain View, CA) in 384-well plates (Dutscher, Brumath, France). Target quantifications were performed with specific primer pairs designed using Universal Probe Library software (Roche Diagnostics, Basel, Switzerland). The primers pairs used in the parallel PCR reaction are described in the Supplementary Table 6. Expression level was normalized to LaEIF-4 (Lalb_Chr07g0195211). All qRT-PCR experiments were performed in two technical replicates as well as a negative control were included in each assay and the values presented represent means ± SD. Relative gene expression levels were calculated according to the ΔΔCt method [51]. All experiments were performed as four biological replicates.

*Delineating the* pauper *region*

Wild type and landraces of white lupin all contain high levels of quinolizidine alkaloids while modern cultivars incorporate one or more mutations conferring low alkaloid content [52] . The most commonly used mutation is *pauper*, a single gene locus that was mapped to linkage group 18[10]. We re-examined marker segregation data from [10] and, taking in account the small degree of marker order ambiguity present in the data, we identified a 6.2 cM map interval flanked by markers TP1572 and TP70046 that we conservatively predicted must contain the *pauper* locus. Sequence tags for TP1572 and TP70046 were aligned to the AMIGA reference genome and physical coordinates defined.

###### *4.2.2 Results*

Measurement of the abundance of key alkaloids in various organs confirmed that the modern variety AMIGA contains low levels of lupanine, 13-hydroxylupanine and 13-tigloyloxylupanine (Extended Fig. 3d). We were not able to match this lower amount of alkaloids with significantly a reduced expression of the known genes coding for enzymes involved in the biosynthetic pathway (Extended Fig. 3e). Next, we explored the gene content of the *pauper* region on Chr18 that is responsible for the low-alkaloid trait in most modern WL varieties, including AMIGA [22–24]. Based on a reappraisal of mapping data presented by [10], we predicted conservatively that pauper is contained within a 958 Kb region containing 66 genes (Supplementary Table 7), including genes coding for transcription factors, transporters, and enzymes. Functional characterization of these genes will lead to a better understanding of the alkaloid content reduction observed in modern varieties and provide the genetic mechanisms underlying the *pauper* locus.

### 5 Genome comparison and evolution

#### 5.1 Evolutionary analysis of Legume genomes

###### *5.1.1 Materials and Methods*

The proposed evolutionary scenario was obtained following the method described in [53] based on synteny relationships identified between *L. albus* (current manuscript), *L. angustifolius* [54] , *Pisum sativum* (under publication), diploid peanut (*Arachis duranensis*, [55]), Lotus (*Lotus japonicus*, [56]), barrel medic (*M. truncatula*, [57]), chickpea (*Cicer arietinum*, [58]), pigeonpea (*Cajanus cajan*, [59]), soybean (*Glycine max*, [60]), common bean (*Phaseolus vulgaris*, [61]), mungbean (*Vigna radiata*, [62]), and adzuki bean (*Vigna angularis*, [63]). Briefly, the first step consists of aligning the investigated genomes to define conserved/duplicated gene pairs on the basis of alignment parameters (CIP for Cumulative Identity Percentage and CALP for Cumulative Alignment Length Percentage). The second step consists of clustering or chaining groups of conserved genes into synteny blocks (excluding blocks with less than 5 genes) corresponding to independent sets of blocks sharing orthologous relationships in modern species. In the third step, conserved gene pairs or conserved groups of gene-to-gene adjacencies defining identical chromosome-to-chromosome relationships between all the extant genomes are merged into Conserved Ancestral Regions (CARs). CARs are then merged into protochromosomes based on partial synteny observed between a subset (not all) of the investigated species. The ancestral karyotype can be considered as a 'median' or 'intermediate' genome consisting of protochromosomes defining a clean reference gene order common to the existent species investigated. From the reconstructed ancestral karyotype an evolutionary scenario was then inferred taking into account the fewest number of genomic rearrangements (including inversions, deletions, fusions, fissions, translocations), which may have operated between the inferred ancestors and the modern genomes.

##### *5.1.2 Results*

We performed a comparative genomic investigation of 12 legume genomes including white lupin (*Lupinus albus*) and covering the Galegoid and Millettoid families. Identification of independent blocks of synteny (see method section) allowed us to propose an ancestral legume karyotype of 17 conserved ancestral regions (CARs), Supplementary table 8. The ancestral genome consists in a minimal shared ancestral genome which lack components of the 'real' (unknown) ancestral genome that were either lost from all of the investigated descendants and/or retained by only one modern species. From the ancestral legume karyotype (ALK) the modern legume genomes experienced lineage-specific rearrangements (mainly ancestral chromosome fusions and fissions) to reach their modern karyotypes as well as specific polyploidization events in the case of soybean and lupin (white and blue), so that their genomes are illuminated as a mosaic of 17 shuffled ancestral CARs (Fig. 2a). From ALK blue and white lupin experience a common triplication event illuminated with a 1-3 chromosomal relationships between ALK and lupin genomes, with the white lupin differentiated from blue lupin though 8 major species-specific chromosomal shuffling events (fusion and fission), Extended Data Fig. 5. From the 17 ancestral chromosomes the modern white Lupin genome experienced a whole genome triplication (with a potential 51-chromosomes intermediate) followed by a massive chromosome number reduction through fusions to reach the 25 modern chromosomes. Our comparative genomics-based evolutionary scenario unravels the legume paleohistory from the reconstructed ALK delivering the complete catalog of paralogous and orthologous gene relationships between 12 modern legume genomes as well as the first reconstructed ancestor of this major botanical family[64]. For that purpose the comparative genomics data described here are made available in the public web tool https://urgi.versailles.inra.fr/synteny/legumes.

#### 5.2 Genome synteny, intragenomic collinearity and gene family identification

##### *5.2.1 Methods*

To identify intragenomic colinearity blocks inside the white lupin genome we used SynMap (CoGe, www.genomevolution.org) using homologous CDS pairs using the following parameters: Maximum distance between two matches (-D): 20; Minimum number of aligned pairs (-A): 10; Algorithm "Quota Align Merge" with Maximum distance between two blocks (-Dm): 500.

Chromosome collinearity between white lupin and narrow-leafed lupin (*Lupinus angustifolius,* v1.0) and *M. truncatula* (Mt4.0) was carried out with MCscan (phyton version) [65,66]. The following parameters were used to detect syntenic blocks: alignment similarity e $\leq 10^{-05}$; average intergenic distance (u) = 40; number of genes required to call synteny, (s) = 5; gap penalty (g) = 2.

We used a comparative analysis to examine the conservation of gene repertoires among orthologs in the genomes of white lupin, narrow-leafed lupin (v1.0) *M. truncatula* (Mt4.0) and *Arabidopsis thaliana* (TAIR10). First, we aligned all-to-all proteins using BLASTP (e-value of $1e^{-5}$). Genes were then clustered using OrthoMCL (1.4) implemented in OrthoVenn [67] with a Markov inflation index of 1.5 and a minimum e-value of $1e^{-15}$.

##### *5.2.2 Results*

Intragenomic analysis for segmental duplications provided evidence of 928 syntenic blocks bigger than 10kb, which retain from 5 to 240 genes. These blocks have an average size of 65 kb and the largest duplication consists of a 4.1 Mb block shared between Chr18 and Chr20 (Fig. 2d, Supplementary Table 9). The results of this analysis showed pinpointing a triplication feature that can be observed in several chromosomes segments (*i.e.* Chr07 that has two homolog regions with Chr12 and one with Chr13).

The WL genome shared highly conserved syntenic blocks with the genome of narrow-leafed lupin (NLL), its closest relative in the Fabaceae family with a sequenced genome[17] and *Medicago truncatula*, the reference genome within this family (Extended Data Figure 4c). We identified *ca*. 420 syntenic blocks between white Lupin *M. truncatula* genomes, with a conservation of 91% of the genes. A lower conservation was detected with narrow-leafed lupin genome, 78%. It is probably due to the incomplete status of the assembly of *L. angustifolius* genome. We could 250 synteny blocks between the two lupin species, and the biggest block has 2.1Mb.

We analyzed coding-sequence deduced protein families shared between white lupin, aforementioned legume species and *Arabidopsis thaliana* (Extended Data Figure 4b). A total of 24,975 protein clusters were generated, corresponding to 77.14% of the deduced protein. The three legume species share 1,787 clusters and WL shares 5,224 clusters with its closest NLL relative. *L. albus* has 473 clusters and 10,495 proteins that did not cluster (Supplementary Table 10). Gene Ontology[20] terms representation revealed an enriched annotation of serine-type carboxypeptidase activity proteins (GO:0004185), however most of the clusters (58%) did not have any GO annotation.

### 6 Soil Exploration

#### 6.1 White Lupin Symbiosis

##### *6.1.1 Material & Methods*

*In silico identification of White Lupin Symbiotic genes*

Symbiotic genes were searched in 96 plant genomes covering different Angiosperm orders and symbiotic abilities (mycorrhization and/or root nodule symbiosis, no symbiosis) as indicated in Supplementary Table 11. Reference proteins previously identified in *M. truncatula* were used as queries for tBLASTn 2.7.1+ [68] and hits with a e-value < 1e-10 were selected for each investigated plant species. Selected sequences were aligned using MAFFT v7.380 [69] with default parameters. When needed, alignments were cleaned using GBlocks [70] implemented in SeaView v4.7 [71]. Alignments were then subjected to phylogenetic analysis using IQ-TREE v1.6.7 [72]. First, the best-fitting evolutionary model was determined for each alignment using ModelFinder [73] before performing maximum likelihood analysis. Branch supports were tested using 10,000 replicates of UltraFast Bootstraps [74]. Tree visualization was performed through the iTOL v4.2.3 platform [33].

*Molecular cloning and hairy-root transformation of white lupin*

The primers for *pLaSYMRK*(Lalb_Chr09g0325781; F-5'-TCGGGTATAAGCAAGCCAAC-3'; R-5'- CTATTTCTGCATCCTTGCTAAGC -3') and *pLaCCamK* (Lalb_Chr19g0132601; F-5'- GGTGTGTGTTAGATAGATGCCTTTG -3'; R-5'-GATGGCTACATGGATTTTCTCAC-3') were designed using Primer3Plus [75]. They were used to amplify a total of 1 963 and 2 185 bp upstream of the start codon of *LaSYMRK* and *LaCCamK*, respectively, from white lupin genomic DNA with the addition of the attb1 (5'-ACAAGTTTGTACAAAAAAGCAGGCT-3') and attb2 (5'-ACCACTTTGTACAAGAAAGCTGGGT-3') adapters. Amplified fragments were subsequently cloned into the pDONR221 by Gateway reaction (Thermo Fisher). The promoters were then cloned into the binary plasmid pKGW-FS7[76] containing a green fluorescent protein-glucuronidase (GFP-GUS) fusion by Gateway cloning. Hairy root transformation of white lupin plants as well as histochemical and microscopic analysis were performed as described before [77].

###### *6.1.2 Results*

The presence of genes known to be specifically required for arbuscular mycorrhizal symbiosis (AMS), root nodule symbiosis (RNS) or both associations was investigated (Supplementary Table 12). Phylogenetic analyses with seven other angiosperm genomes covering the diversity of symbiotic abilities [78,79] (revealed that RNS specific genes (such as NFP, EPR3, NIN, RPG, PUB1) along with gene required for both symbiosis (such as SYMRK, CYCLOPS, CCamK, VAPYRIN…) are retained in *Lupinus albus* and are expressed (Supplementary Fig. 6). By contrast, AMS specific genes (PT4, RAM1, RAM2, FatM, HA1, STR1, STR2 and Exo70) were not retrieved in the *Lupinus albus* genome (Supplementary Figs. 16-42). This result expands previous findings on the draft genome *of Lupinus angustifolius* and supports the hypothesis that loss of AMS occurred early during the evolution of the *Lupinus* genus or their genistoid progenitors.

##### 6.2 Candidate genes controlling cluster root formation

###### *6.2.1 Material and Methods*

*Spatial transcriptome for mRNA and small RNA*

Seeds of white lupin (*L. albus* L. cv. AMIGA) were germinated for 4 days on vermiculite soaked with water. Thereafter, the seedlings were transferred to an aerated nutrient solution in the absence of phosphate in growth chambers under controlled conditions (16h light/8h dark, 25°C day/20°C night, 65% relative humidity and PAR intensity 200 µmol.$m^{-2}$.$s^{-1}$).

After 12 days of culture, ten cluster roots coming from four grown plants were harvested and dissected in eight parts of 0.5-cm from the apex of the lateral root that carries the cluster root. As control, 1-cm of lateral roots without cluster roots, sampled 1-cm away from the primary root, were collected. Four biological replications were produced for each experiment.

Total RNA was extracted from all frozen samples using the Direct-zol RNA MiniPrep kit (Zymo Research, Irvine, CA) according to the manufacturer’s recommendations.

For mRNA sequencing, 36 independent root RNA-seq libraries were constructed and sequenced at Get-PlaGe core facility (INRA, Toulouse, France). The Illumina TruSeq Stranded mRNA Sample Preparation Kit (Illumina Inc.) was used according to the manufacturer's protocol. The samples were sequenced using Paired-end sequencing was performed generating pair-ended 2 x 150 bp reads using TruSeq SBS kit v3 sequencing chemistry (Illumina Inc.) in one lane of Illumina NovaSeq instrument according to the manufacturer's instructions. A total of 2,048,118,650 paired-end reads of 150 pb were sequenced using an illumina NovaSeq6000 Sequencer. To remove low quality sequences, the RNA-seq reads were checked and trimmed with a minimum quality score of 30 in both 3' and 5' end. The resulting reads shorter than 35 pb have been discarded. The quality checked RNA-seq reads were then mapped on white lupin reference genome using Hisat2 software. Transcripts were assembled and quantified using Stringtie software. Gene counts were extracted and imported in the R package DESeq2. These counts have been normalized according to the size factor computed by DESeq2.

For small RNA sequencing, 24 independent root RNA-seq libraries were constructed and sequenced at Transcriptomic Platform IPS2 (IPS2, Paris, France). The NEXTflex™ Small RNA-Seq kit was used for generation of small RNA-seq libraries according to the manufacturer's protocol. All small RNA libraries were sequenced on an Illumina NextSeq 500 sequencing platform, using a single-end, 75 nt read metric instrument according to the manufacturer's instructions. A total of 460,506,072 reads of 75 nt were sequenced. Small RNA-seq reads were trimmed using cutadapt version 1.11 [23] to remove remnants of the following 3'-adapter sequence. Only sequences that had the adapter were kept, insuring that they are small fragments. The adapter trimming was performed based on the adapter sequence: TGGAATTCTCGGGTGCCAAGGAACTCCAGTCAC of 33 nucleotides long, removing the adapter if the length of the match is at least 10 nts. A second round of trimming was performed with cutadapt to remove the four nts on the 5' and 3' ends of the reads (which corresponded to the four degenerated nts on each "High Definition" (HD) adapter). Empty reads (adapter dimer) and all the sequences shorter than 15 nucleotides were discarded. A total of 424,385,998 trimmed reads were kept for final analysis. The sequence files produced were collapsed to include only non-redundant reads and the number of occurrences of each. To quantify the level of expression of Lupin miRNAs we used the high conservation in sequence of the main miRNA families among plant species. We therefore use the publicly available mature miRNA of three well studied species in the Fabaceae clade max present in miRBase v22[80]: *Medicago truncatula, Lotus japonicas* or *Glycine max*. Only sequences corresponding exactly to the published mature miRNA sequence were retained and counted.

*AMIGA and GRAECUS root sampling and expression analysis of cluster root initiation genes*

We sampled 2-3 cm of lateral roots 1 cm away from the primary root in the top 5 cm (cluster root region, CRR) and at 10 cm from the top (regular lateral root region, NLR) of the root system of AMIGA and GRAECUS plants 9 days after germination. Three CRR and 3 NLR independent samples were collected for each accession. Total RNA from these samples was extracted using the Direct-zol RNA MiniPrep kit (Zymo Research, Irvine, CA) according to the manufacturer's recommendations. RNA concentration was measured on a NanoDrop (ND1000) spectrophotometer. Poly(dT) cDNA were prepared from 2 µg total RNA using the revertaid First Strand cDNA Synthesis (Thermo Fisher). Gene expression was measured by quantitative Real Time - Polymerase Chain Reaction (qRT-PCR) (LightCycler 480, Roche Diagnostics, Basel, Switzerland) using the SYBR Premix Ex Taq (Tli RNaseH, Takara, Clontech, Mountain View, CA) in 384-well plates (Dutscher, Brumath, France). Target quantifications were performed with specific primer pairs designed using Universal Probe Library software (Roche Diagnostics, Basel, Switzerland). The primer pairs used in the parallel reactions are described on the Supplementary note table 11. Expression levels were normalized to *LaHelicase* (Lalb_Chr13g0304501). All qRT-PCR experiments were performed in technical quadruplicates. Relative gene expression levels were calculated according to the ΔΔCt method[51], using as a calibrator the NLR samples. All experiments were performed as three biological replicates.

**Supplementary Note Table 11. Genes and primer pairs used to study relative expression of cluster root initiation genes in AMIGA and GRAECUS**

| Gene | Id | Primer forward | Primer reverse |
|---|---|---|---|
| LaPUCHI-1 | Lalb_Chr07g0177601 | gagggagaagaaagcaagca | gcagcatatctaccccaagg |
| LaSTART | Lalb_Chr20g0121101 | caaccaaaagaaggggttga | tgaggcttcttccctatcagc |
| LaCLE-1 | Lalb_Chr19g0133921 | caacaataggcaaaagggttagt | cctagcattgacctctttgaga |
| LaPEPT-1 | Lalb_Chr16g0388291 | atgcattcagacccacatca | tttcttctgcatatttgtgacca |
| LaPME41 | Lalb_Chr06g0166531 | ggacgggttaccatgtgatt | ggtgtaagtaactcccgtctgtg |
| LaMYB1 | Lalb_Chr20g0122341 | aaagagaggccggtttacct | gccaagaatgctatgtagctga |
| LaHelicase | Lalb_Chr13g0304501 | caattctgaaggctatactatgtgc | tcaccagctgctacattaggg |

##### *6.2.2 Results*

The capacity of WL to produce cluster roots (CR) prompted us to perform a detailed spatial RNA-seq survey (mRNA and miRNA) into the 8 developmental sections of mature clusters of the AMIGA variety (Fig. 3a). To check the quality among the experimental replicates, we verified the clustering of the samples in a principal component analysis (PCA), Supplementary Fig. 7. The first two principal components are explaining 79% of the total variance among the samples. The PCA plot of the samples in the 2D plane spanned by their first two principal components shows that they are grouped by root region (LR-S7).

We performed a differential expression analysis of genes from each CR part (S0-S7, Fig. 3a) comparing to regular lateral root (LR - lateral root that does not produce CR). We identified 591 up-regulated genes (Supplementary Table 13) in the CR parts and 669 down-regulated genes (Supplementary Table 14). S6 and S7, the two zones containing the mature rootlets, shown the major number of overexpressed genes (261 in total, Fig. 3b). A GO enrichment analysis of the individual parts showed a strong enrichment in GO terms associated with membrane components in these two parts, linked with their highly active physiology[81,82]. This is confirming their strong capacity to remobilize and acquire phosphate efficiently (Supplementary Figs. 9-11).

We produced a matrix representing all intersections of up-regulated (Fig. 3b) and down-regulated (Supplementary Fig. 8) genes in the CR parts. Interestingly, a list of 42 genes overexpressed in all cluster roots parts (Supplementary Table 15,

Supplementary Fig. 12) showed a strong enrichment in transcription factors (43%) and 9 of them belong to the AP2/EREBP family[83]. This is a large multigene family, and they are key regulators of several developmental processes, like floral organ identity determination, control of leaf epidermal cell identity and control of lateral root development. In white lupin, we identified 217 genes in this family.

Similarly, a list containing only the genes overexpressed in the S1 region, where the early time events of the CR development happens, is also enriched with transcription factors (6 out of 16) and there is an overexpression of 3 genes of AP2/EREBP family that are orthologs of *AtPUCHI*, a gene that is required for morphogenesis in the early lateral root primordium of Arabidopsis[84]. We performed an identification of all the orthologs genes of the AP2/EREBP subfamily B-1, which the gene *AtPUCHI* belongs. We identified 20 orthologs in the white lupin genome and 4 orthologs of the gene *PUCHI* (Fig. 3c, Supplementary Fig. 13).

In another approach, we identified all mature microRNAs that are expressed in the CR. We identified 103 miRNA cluster families in the CR, which 29 are predictions (Fig. 3b detail, Supplementary Table 15 and Supplementary Figure 14). Some of the known miRNA families that we identified were already described as related with P-deficiency response, as miRNA156, miRNA166 and miRNA2111[85]. For the first time it was detected the presence of miRNA399 family, a key regulator for the phosphate starvation response[86,87], in CR of white lupin[88,89] (Supplementary Fig. 14a, Supplementary Table 16).

We identified that 14 genes out of the group of 42 overexpressed in all CR zones are miRNA targets, including 5 transcription factors (Supplementary Table 15). Likewise, in the group of 16 genes that are only overexpressed in the region S1, we identified 5 genes that are targets of the miRNAs detected, comprising transcription factors *LaWRKY* (Lalb_Chr07g0182001) and *LaPUCHI-3* (Lalb_Chr18g0055601).

Comparing the root system of AMIGA and GRAECUS, we identified that AMIGA has an earlier establishment of CRs than the wild-relative GRAECUS, with a strong increase in lateral and cluster root production in the top part o the root system (Fig. 3d, Supplementary Fig. 15). Therefore, we checked how some of the genes with an overexpression in S1 (63 genes, Fig. 3e) are expressed in the top of the root system of both accessions. There is an increased level of expression of potential regulator genes such as *LaPUCHI-1* (Fig. 3e). A list of candidate genes selected on their high induction level at the S1 stage (*LaCLE1, LaMYB1, LaPEP1, LaPME41* and *LaSTART*), also showed a higher expression level in AMIGA compared to its wild relative GRAECUS (Extended Data Fig. 6).

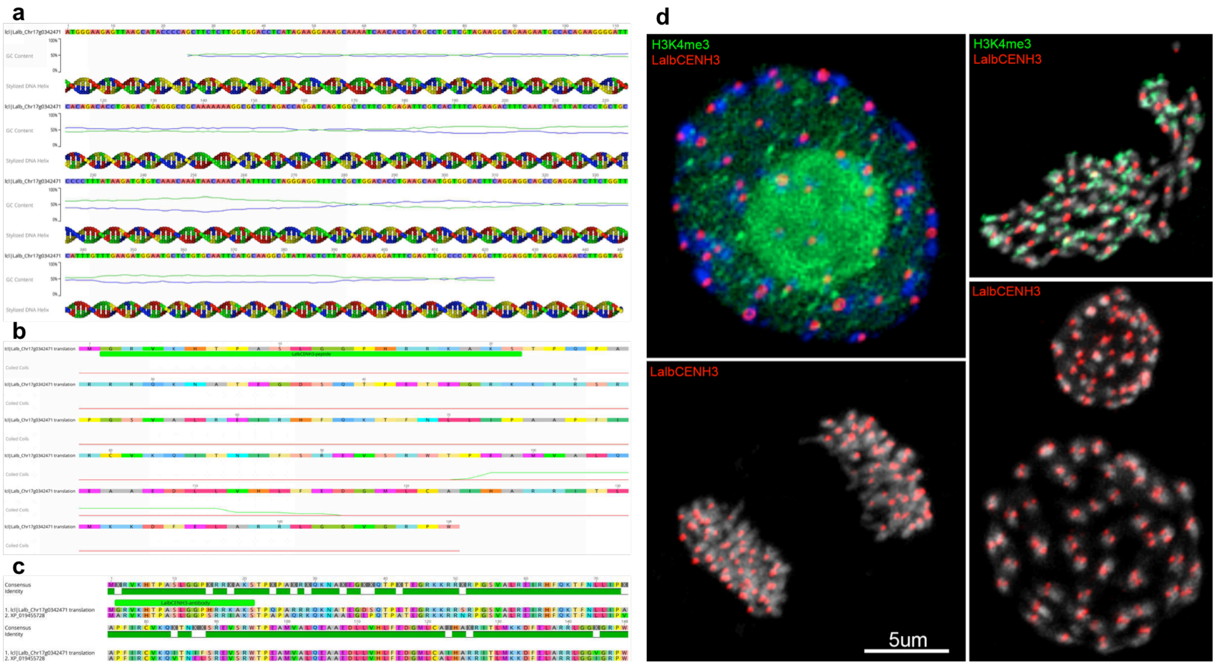

**Supplementary Figure 1. Characterization of *Lupinus albus* CENH3. (a**) Coding DNA sequence; **(b)** Deduced amino acid sequence; **(c)** Alignment of LalbCENH3 and target peptide site for specific antibody generation. XP_019455728 refers to the Genbank accession number of the *Lupinus angustifolius* CENH3 sequence. **(d)** Immunostaining of *L. albus* chromosomes with LaCENH3 (red) and H3K4me3 (green).

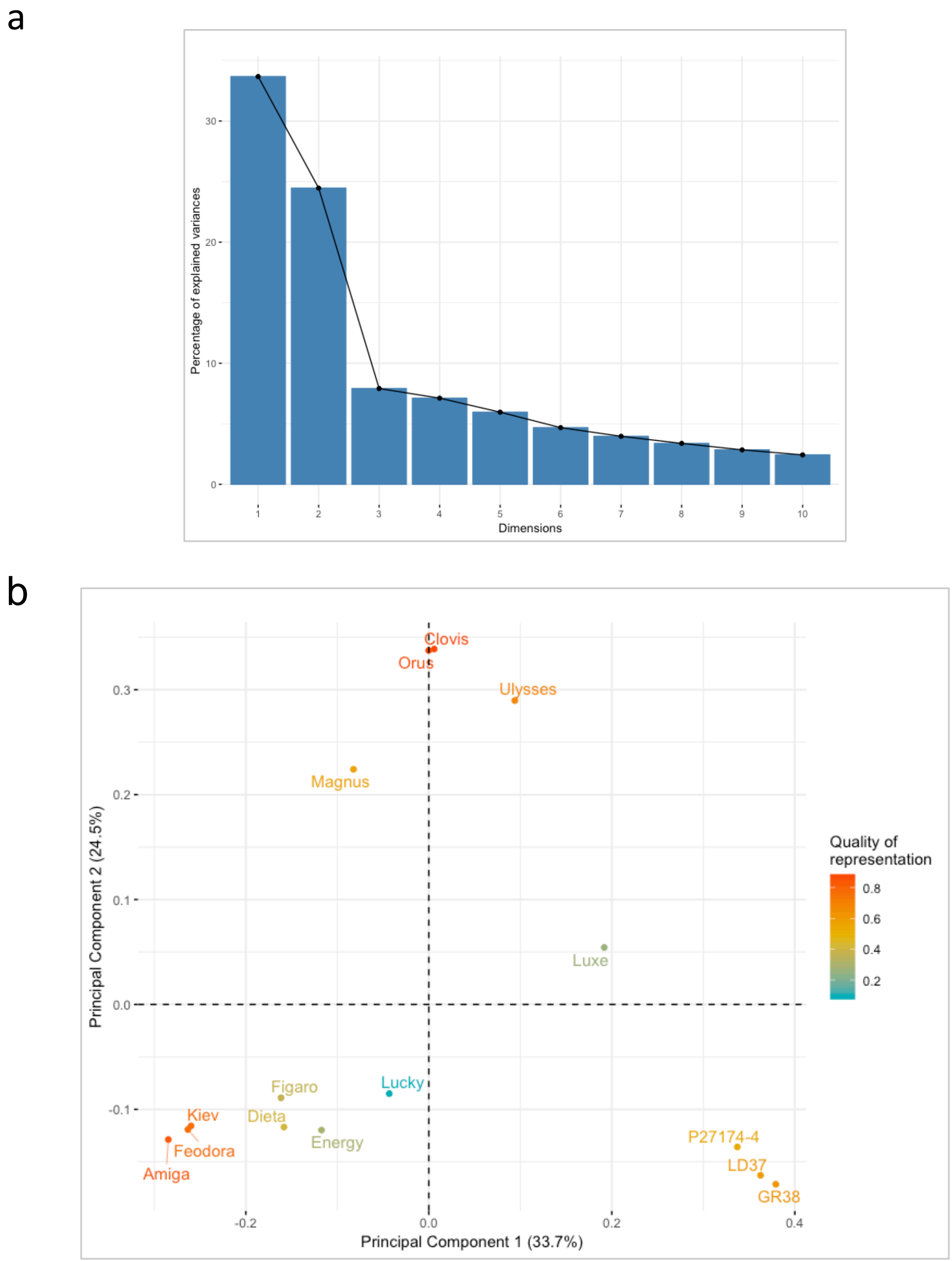

**Supplementary Figure 2. Principal component analysis of 15 white lupin accessions. (a)** Screen plot showing the percentage of variances explained by the 10 first principal components. **(b)** Scatter plot of the distribution of the 15 accessions in the two first principal components, explaining 58.2% of the total variance. The distribution of the accessions clearly reflects the pattern observed in the phylogenetic tree.

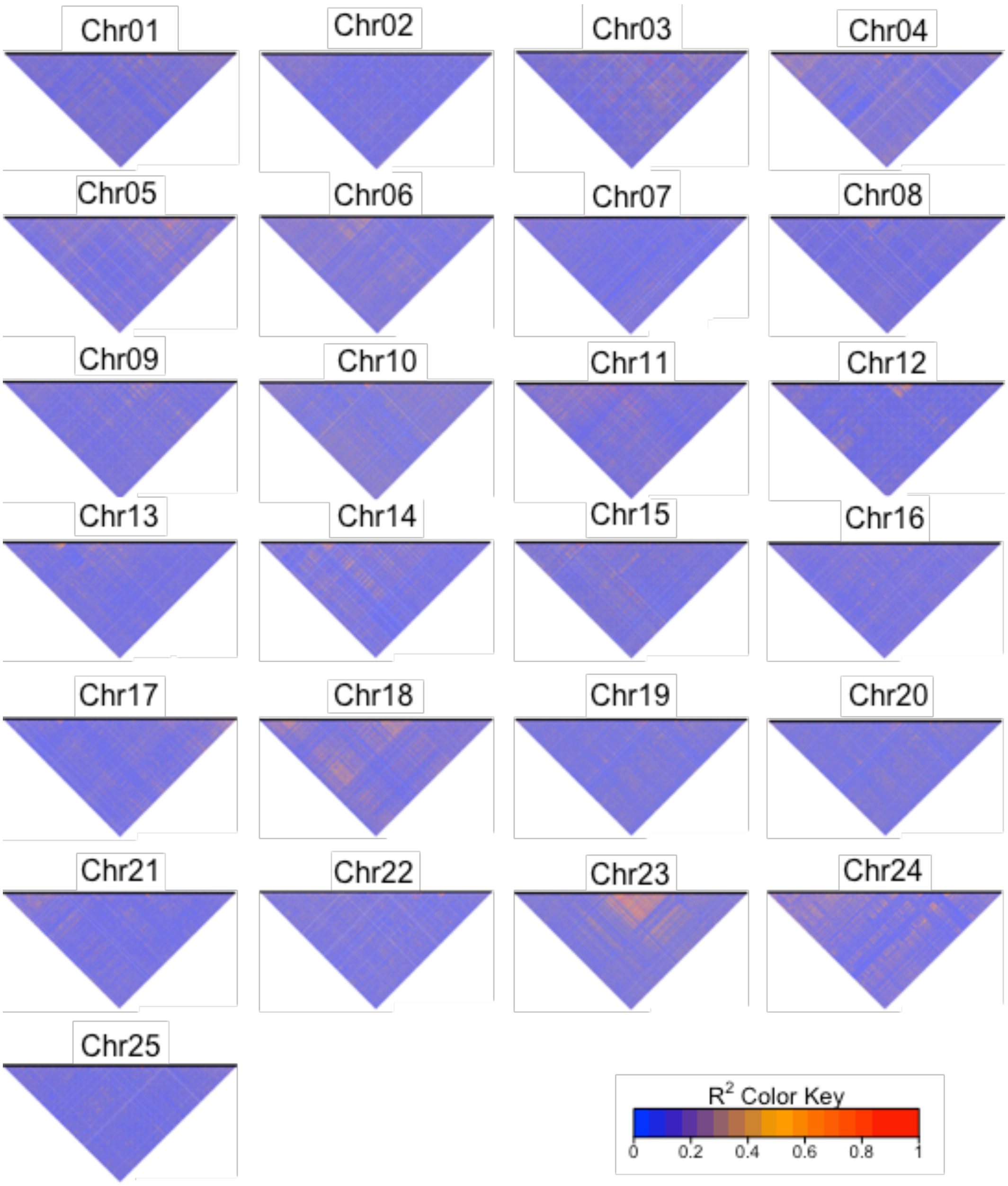

**Supplementary Figure 3. Linkage disequilibrium analysis for the 25 white lupin chromosomes.** Genotypic linkage disequilibrium (LD, $r^2$) represented by heatmap for all white lupin chromosomes. The black bar in the top of the figure is a graphical representation of physical location of the SNPs on the chromosomes. The color coding represents high LD (red) and low LD (blue).

a

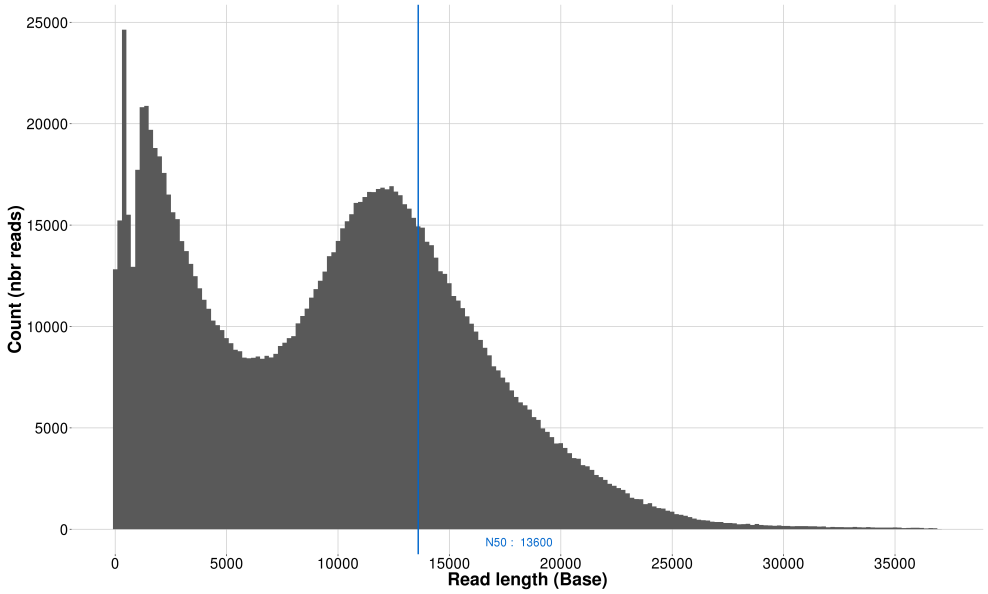

b

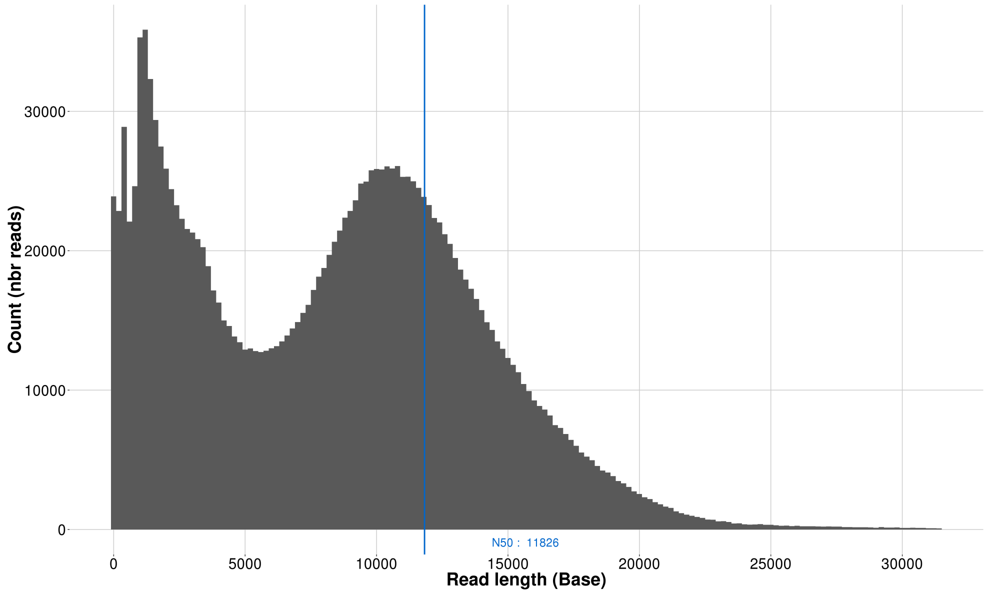

**Supplementary Figure 4. Reads length distribution of GRAECUS and P27174 genome sequencing using Oxford Nanopore technology.** (a) Results presented for genotype GRAECUS, with a N50 size of 13.6 Kb and (b) P27174 with a N50 size of 11.8 Kb.

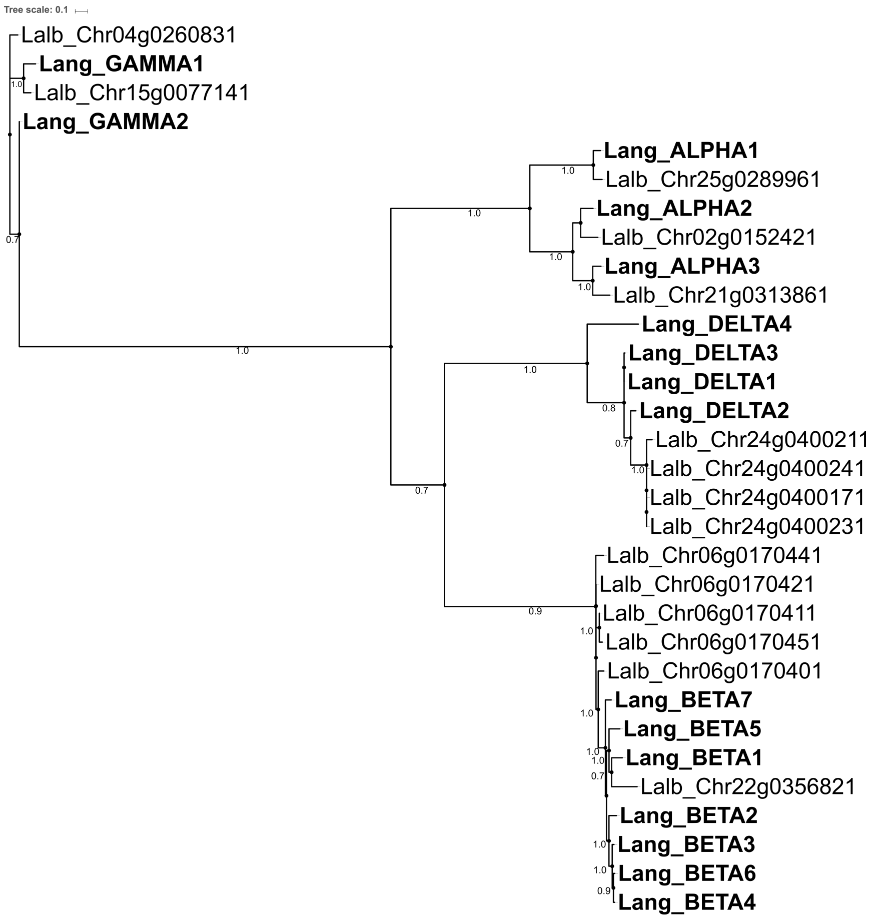

**Supplementary Figure 5. White lupin conglutins.** Maximum likelihood phylogenetic tree of conglutin orthologs of *L. albus* and *L. angustifolius* (in bold).

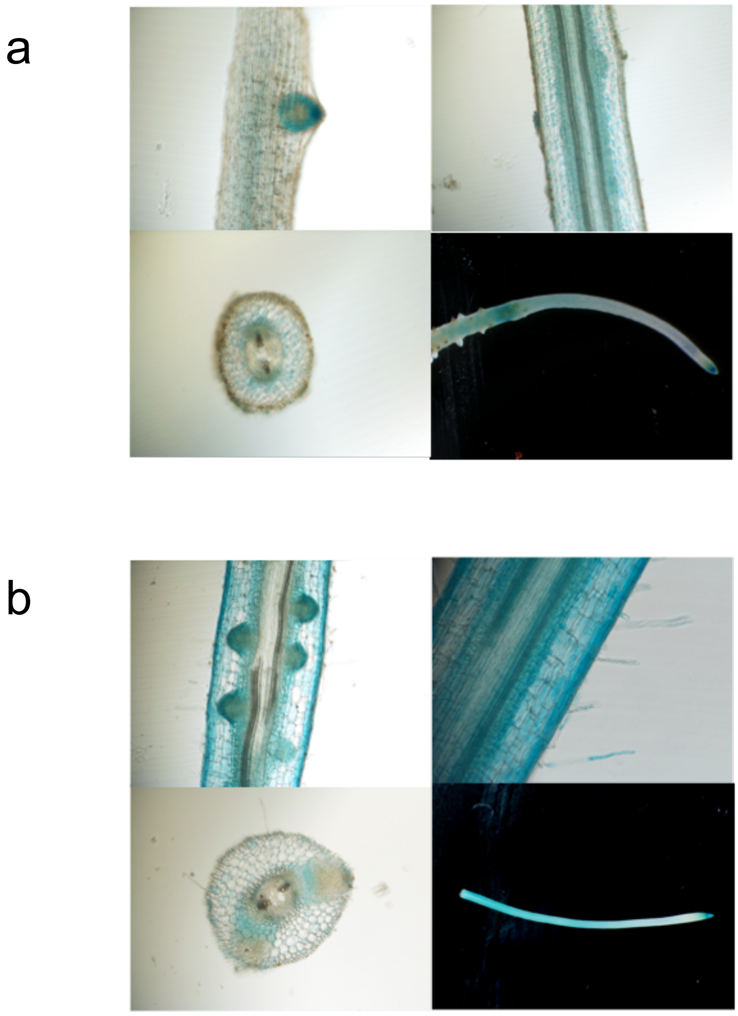

**Supplementary Figure 6.** Expression pattern of *pLaSYMRK:GUS* **(a)** and *pLaCCamK:GUS* **(b)** in *hairy roots* of 4-week-old transformed plants grown in low nitrate condition and presence of *Bradyrhizobium lupini LL13.*

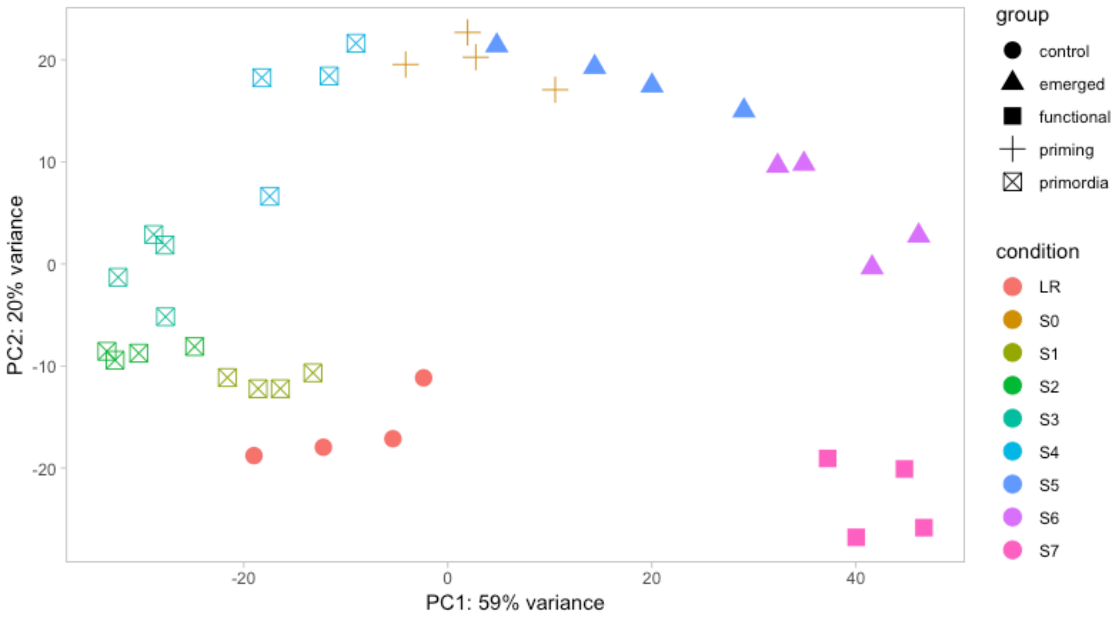

**Supplementary Figure 7. Scatterplot of the two first principal components over counts matrices of cluster root detailed development RNAseq.**

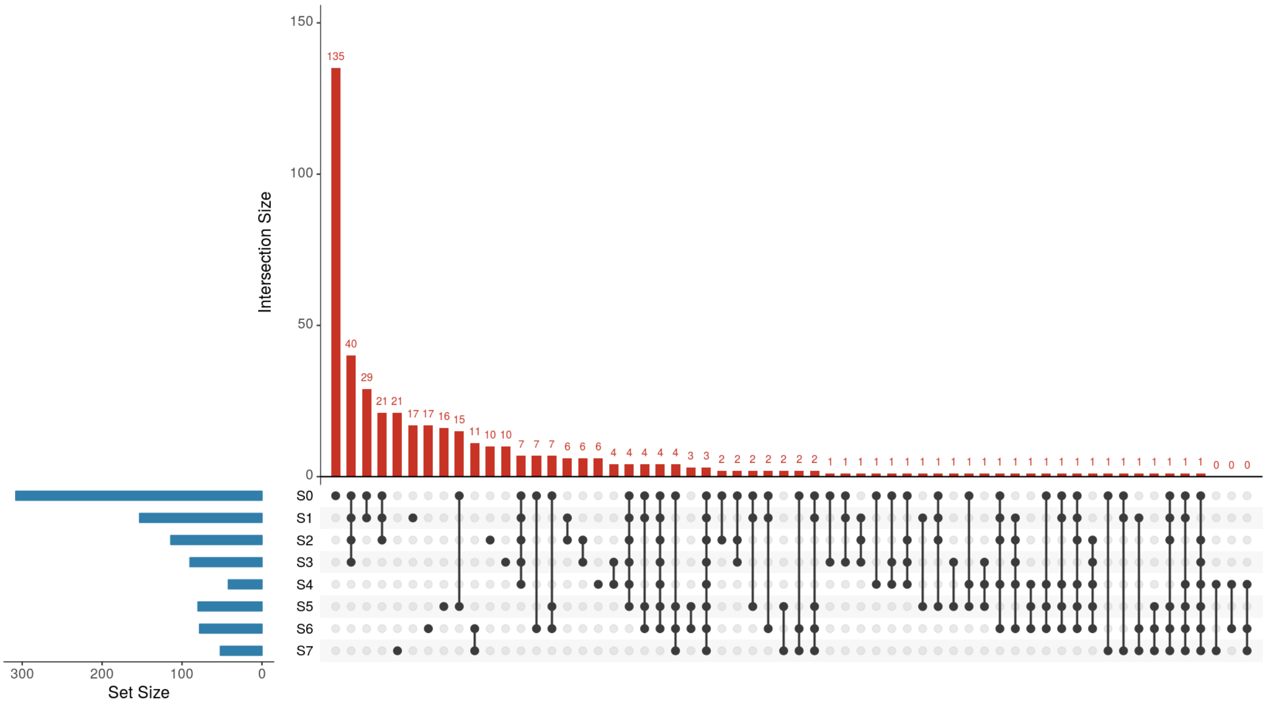

**Supplementary Figure 8. UpSet graph of genes down regulated in cluster root parts (S0-S7) in comparison with lateral roots (LR) .** The UpSet graph displays the number of down-regulated expressed genes among the CR parts. For each comparison, the number on top of each bar represents the number of differentially expressed genes (intersection size). The comparison in question is indicated by the dots or connected dots below its respective bar. Set size indicates the total number of genes for each comparison.

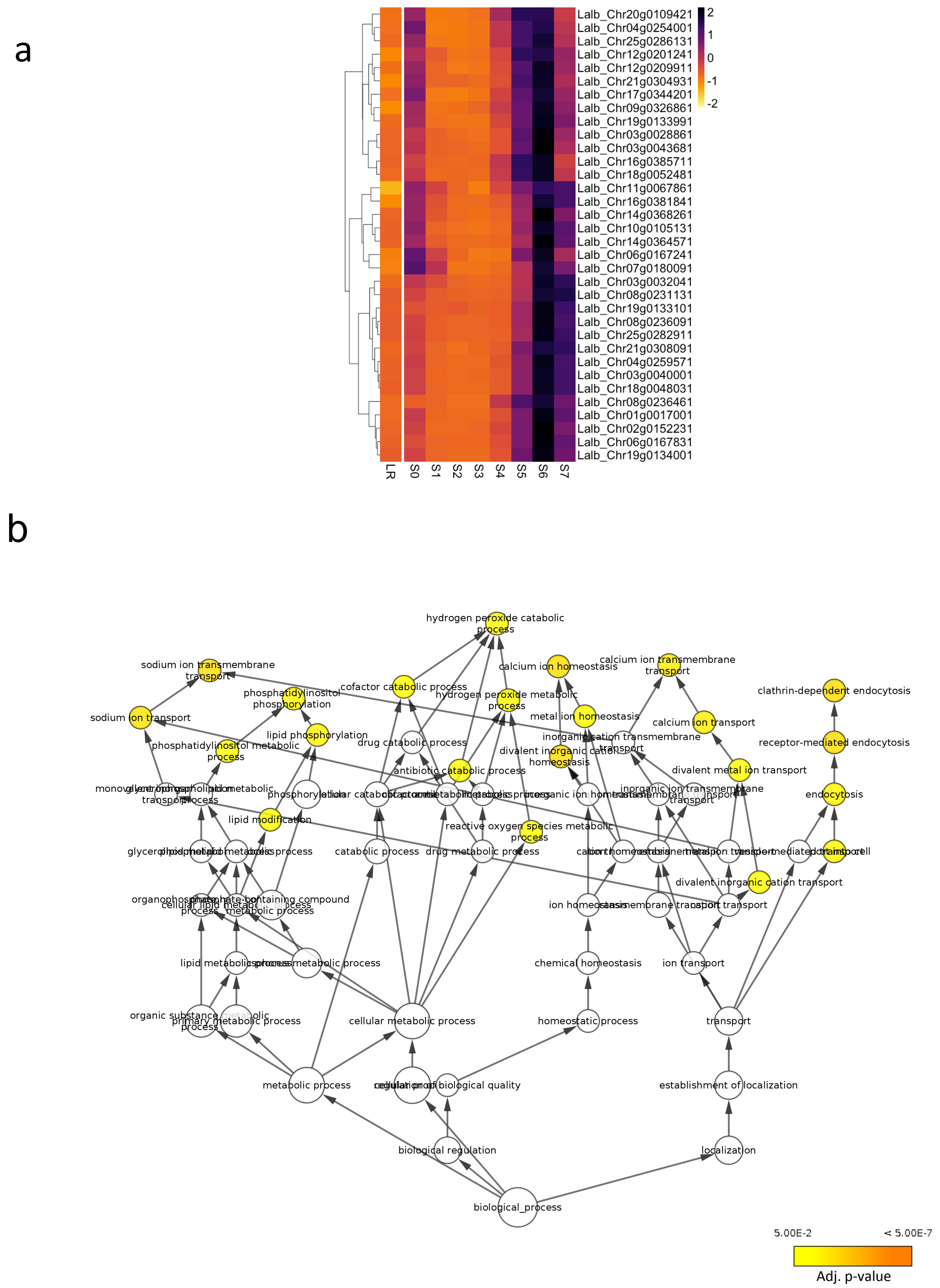

**Supplementary Figure 9. GO terms enrichment for the genes up regulated in region S6. (a)** Heatmap of the 34 genes up-regulated in the zone S6 of the cluster root. **(b)** Network of the GOs of these genes. The color represents the p-value.

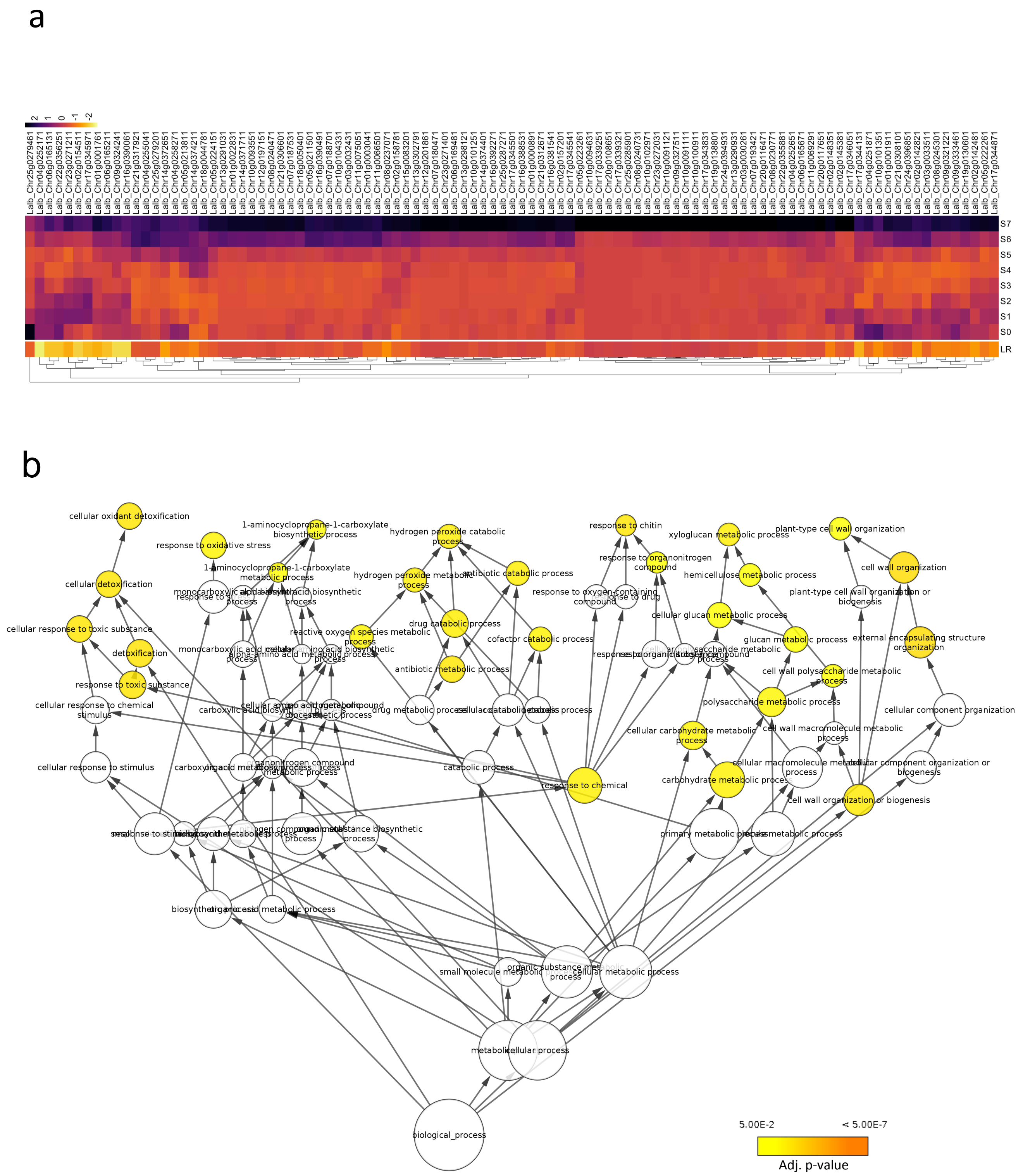

**Supplementary Figure 10. GO terms enrichment for the genes up regulated in region S7. (a)** Heatmap of the 101 genes up-regulated in the zone S7 of the cluster root. **(b)** Network of the GOs of these genes. The color represents the p-value.

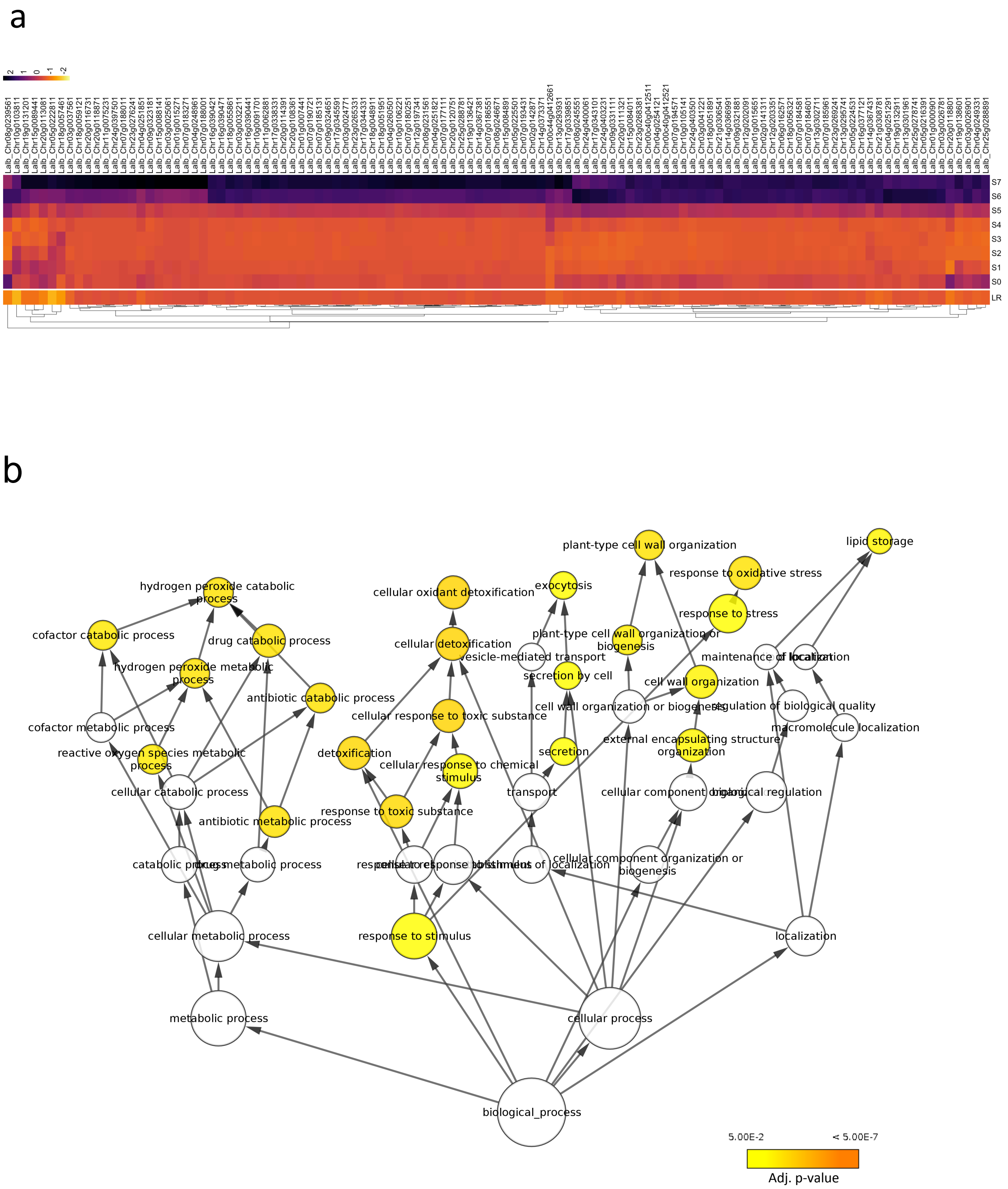

**Supplementary Figure 11. GO terms enrichment for the genes up regulated in region S6 and S7. (a)** Heatmap of the 111 genes up-regulated in the zones S6 and S7 of the cluster root. **(b)** Network of the GOs of these genes. The color represents the p-value.

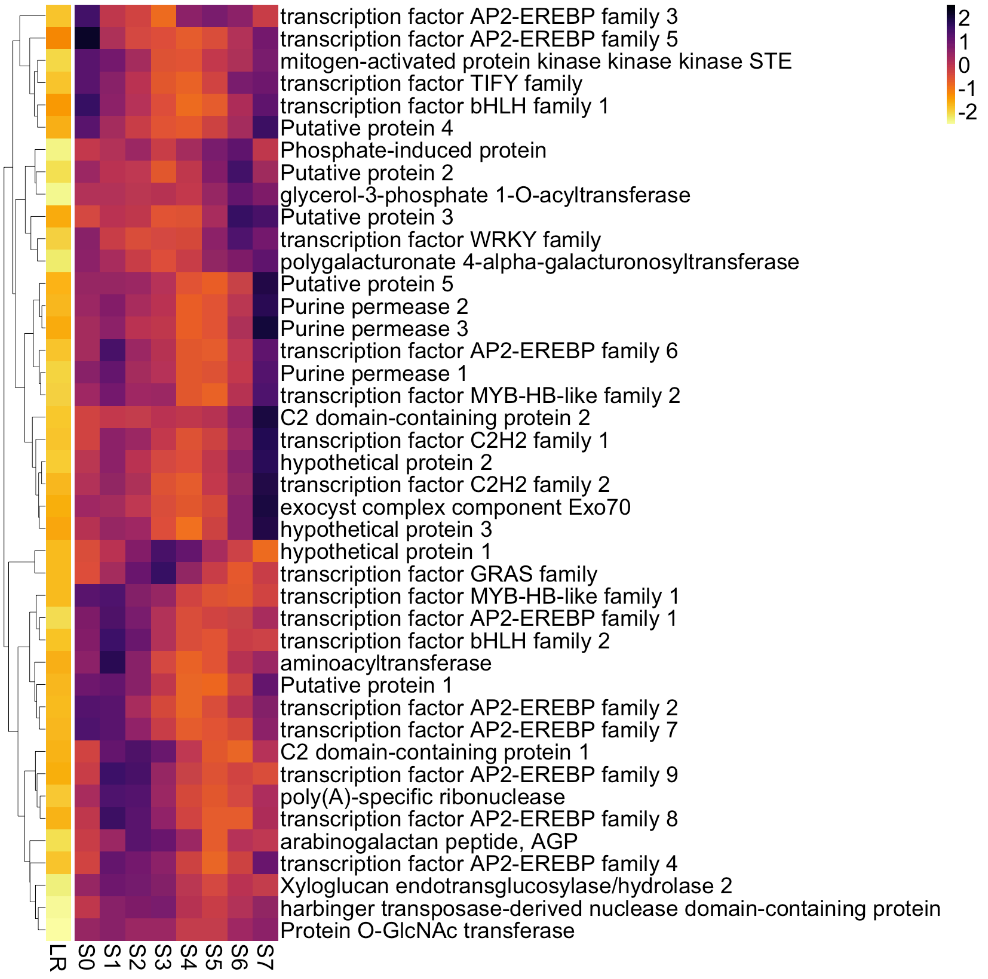

**Supplementary Figure 12. Group of 42 genes overexpressed in all cluster root zones.** Automatic annotation is shown. There is an enrichment of transcription factors (43%).

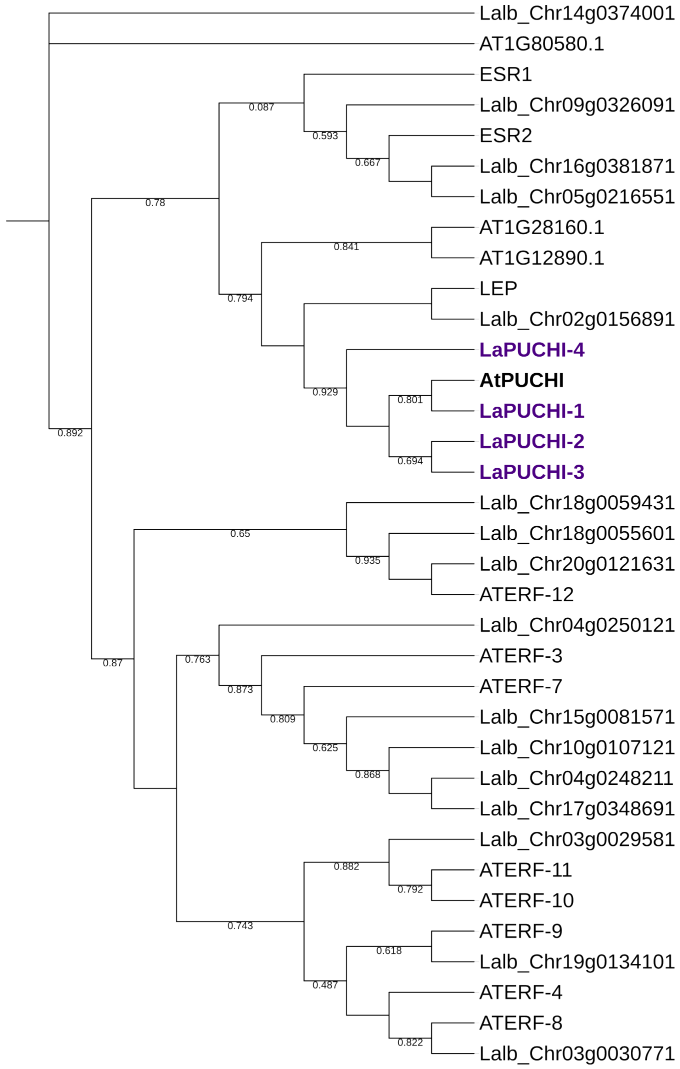

**Supplementary Figure 13. Maximum likelihood phylogenetic tree of Arabidopsis ERF (ethylene response factor) subfamily B-1 of ERF/AP2 transcription factor family and white lupin orthologs.** These proteins contains an AP2 domain. There are 15 members in this subfamily in Arabidopsis and there are 20 orthologs in white lupin genome. The Arabidopsis thaliana PUCHI (*AtPUCHI*) is highlighted, as well the four white lupin homologs named as *LaPUCHI-1* (Lalb_Chr07g0177601 ), *LaPUCHI-2* (Lalb_Chr13g0303751), *LaPUCHI-3* (Lalb_Chr187g0059441) and *LaPUCHI-4* (Lalb_Chr15g0086951).

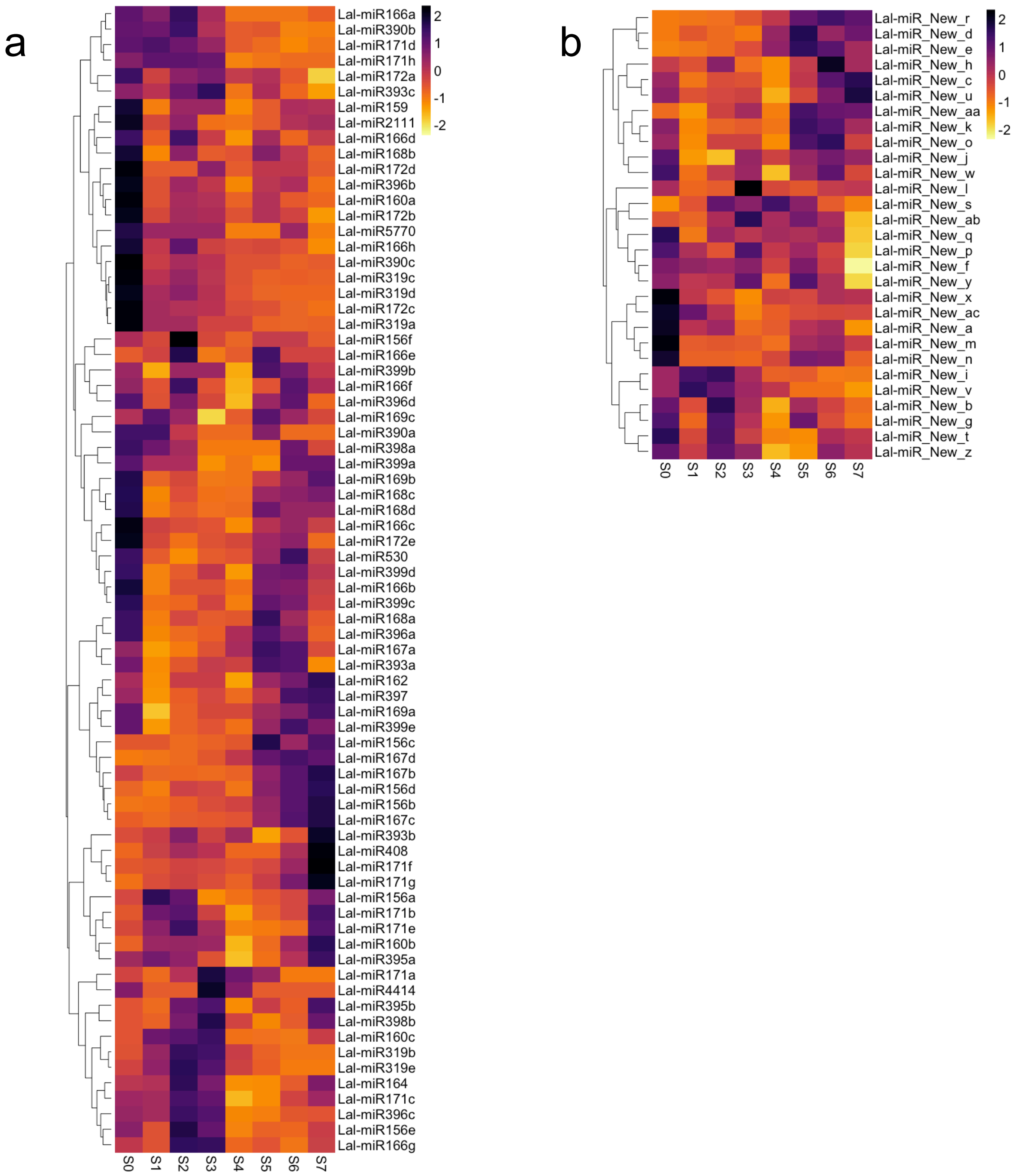

**Supplementary Figure 14. Heatmap of expression pattern of known miRNA families in the cluster root zones.** 103 miRNA cluster families were identified in the CR, out of which 74 are known **(a)** and 28 are new-described miRNA cluster families **(b)**. More information about the white lupin miRNA cluster families can be found in Supplementary Table 16.

a

b

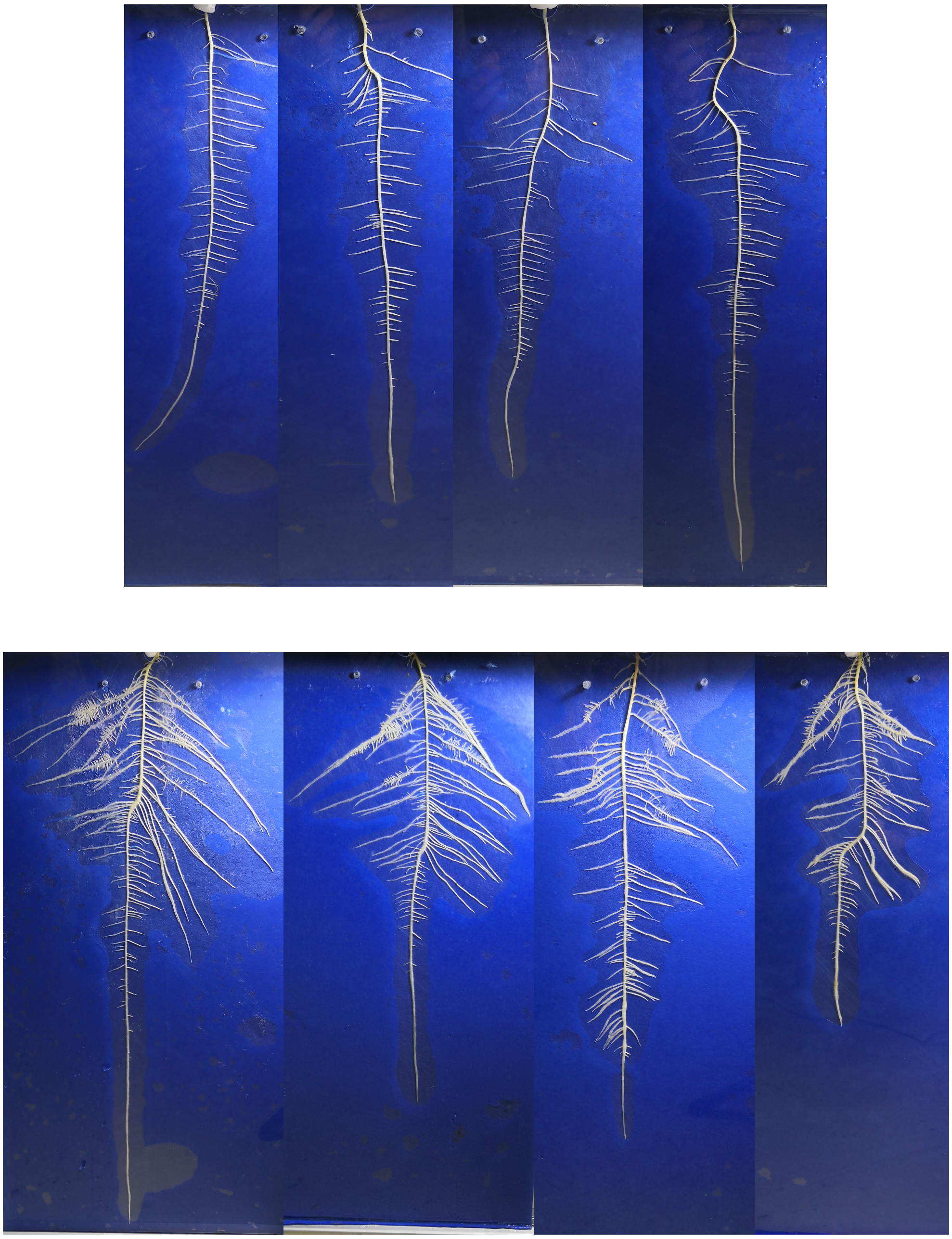

**Supplementary Figure 15. Root system architecture of wild variety GRAECUS (a) and *cv.* AMIGA (b).** Four representative images of 11-day-old white lupin grown vertically on a blotting paper in low phosphate conditions.

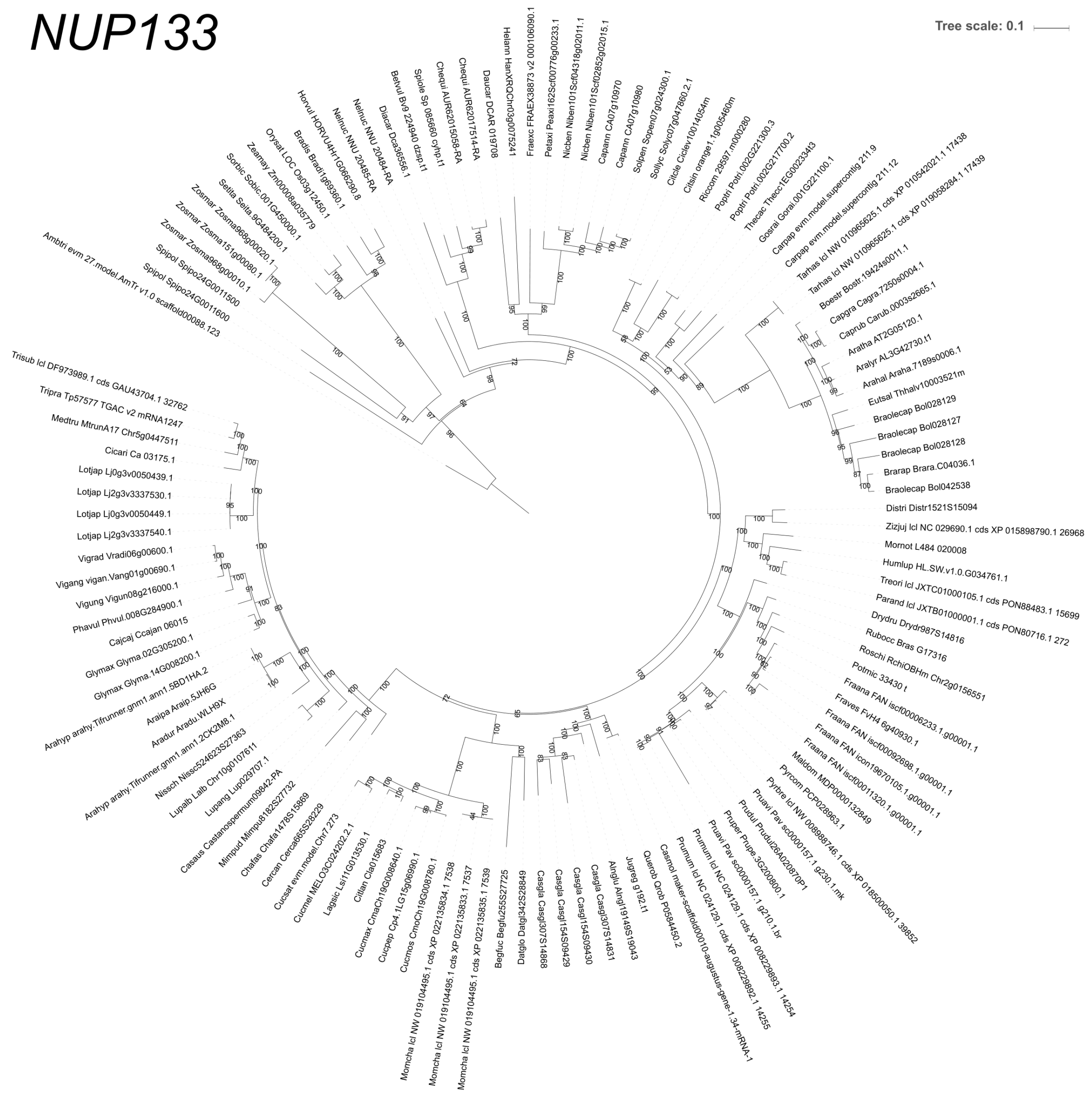

**Supplementary Figure 16. Phylogenetic tree of *NUP33* homologs of 96 plant species, covering different Angiosperm orders and symbiotic abilities.**

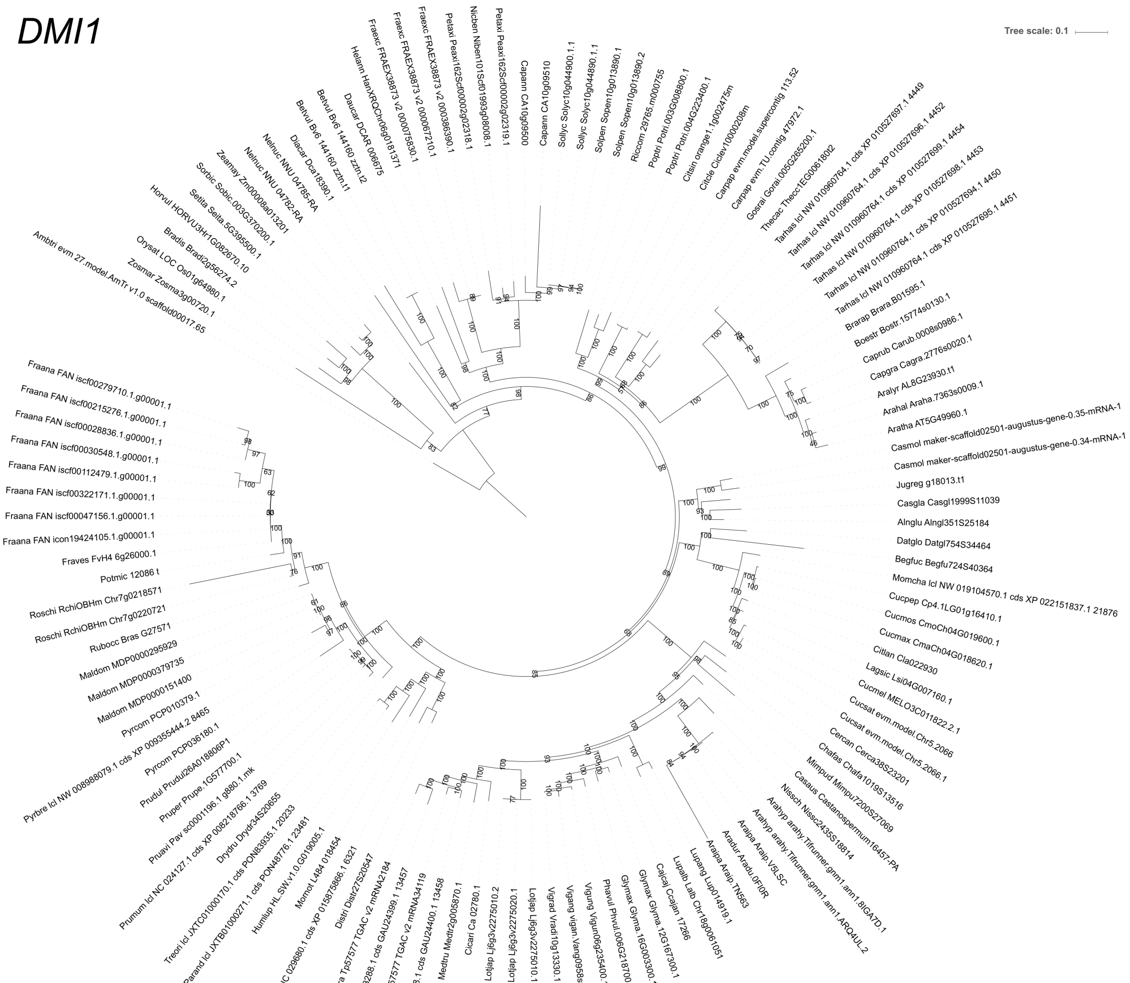

**Supplementary Figure 17. Phylogenetic tree of *DMI1* homologs of 96 plant species, covering different Angiosperm orders and symbiotic abilities.**

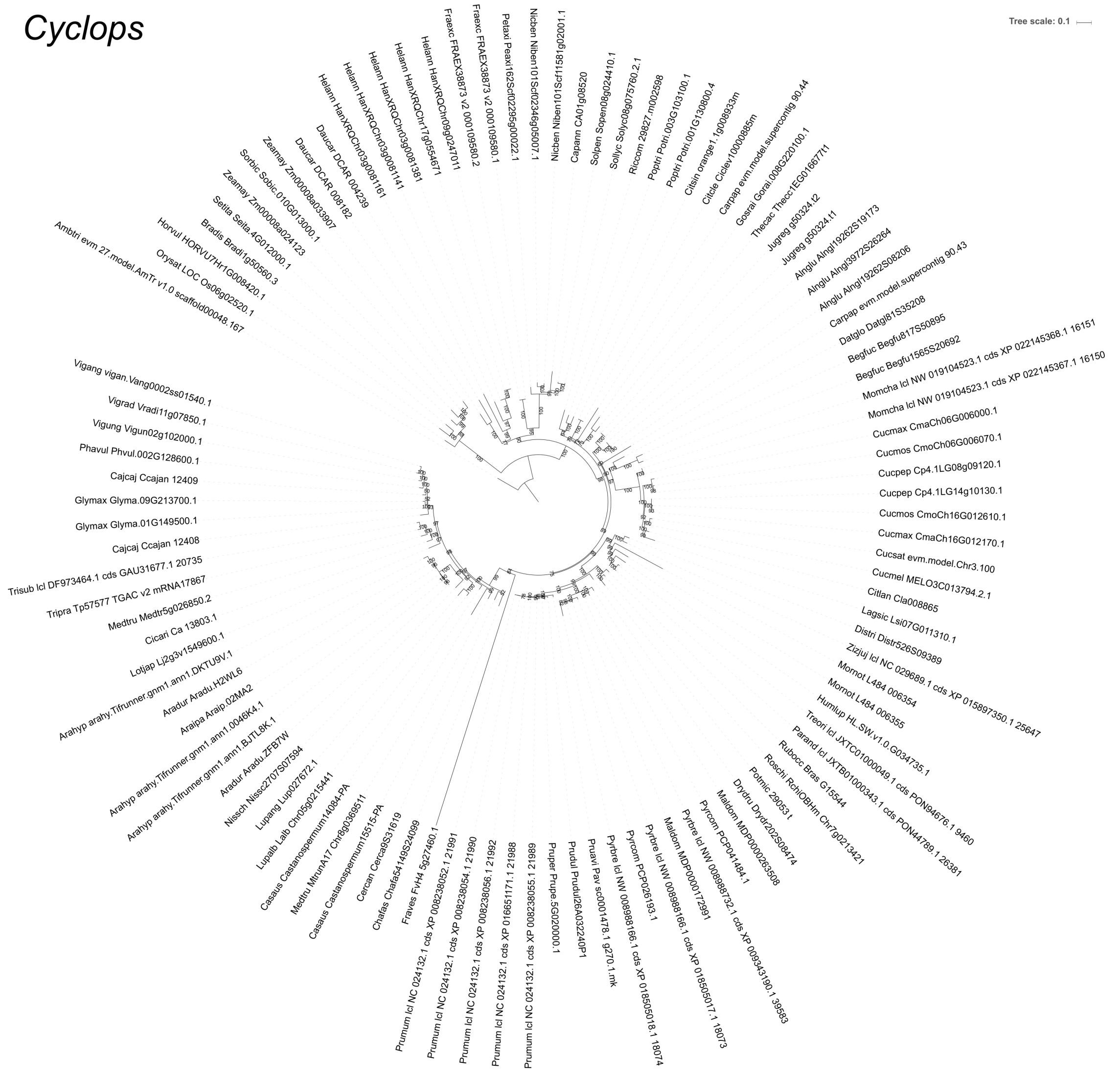

**Supplementary Figure 18. Phylogenetic tree of *CYCLOPS* homologs of 96 plant species, covering different Angiosperm orders and symbiotic abilities.**

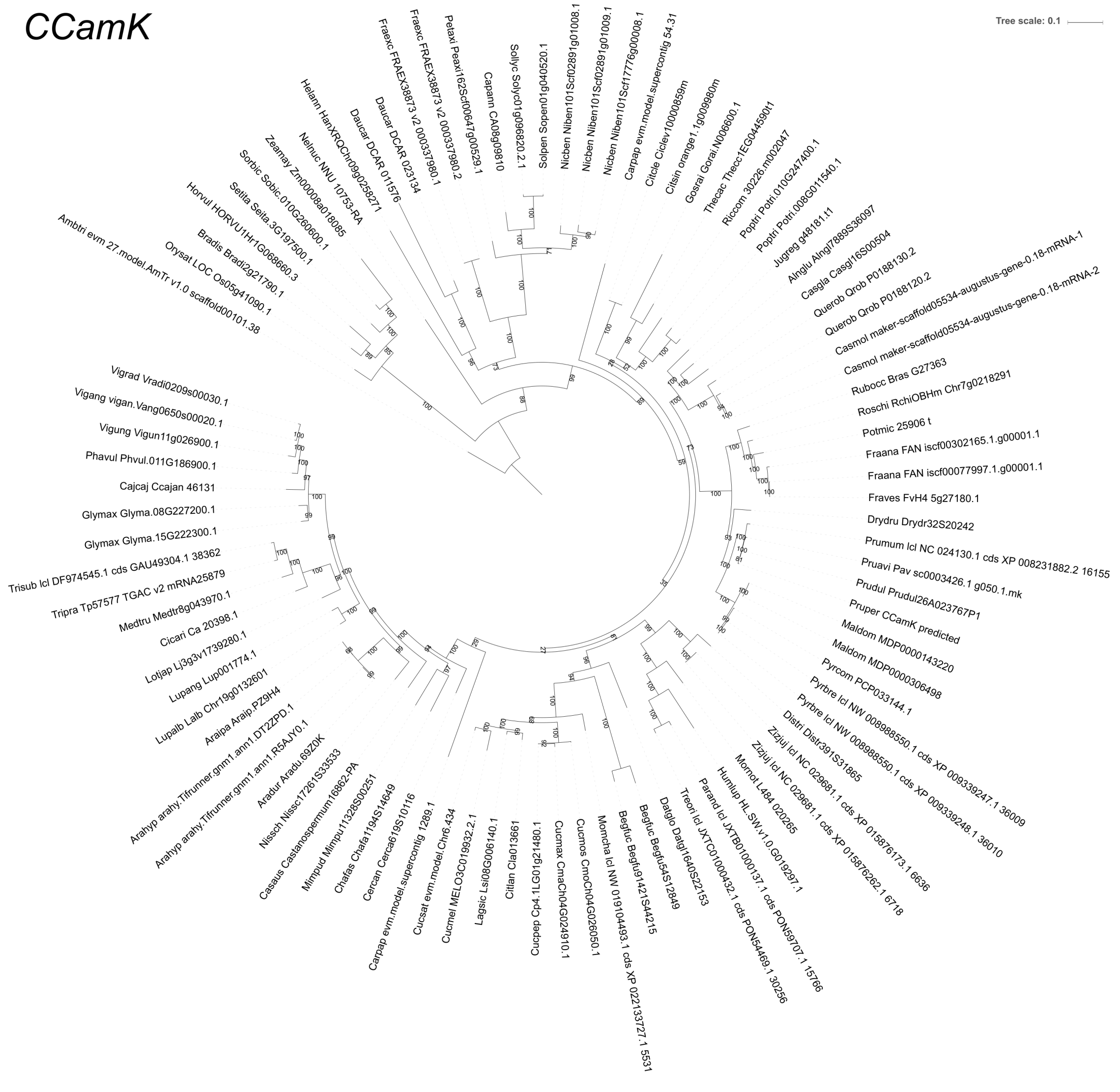

**Supplementary Figure 19. Phylogenetic tree of *CCamK* homologs of 96 plant species, covering different Angiosperm orders and symbiotic abilities.**

### *NIN*

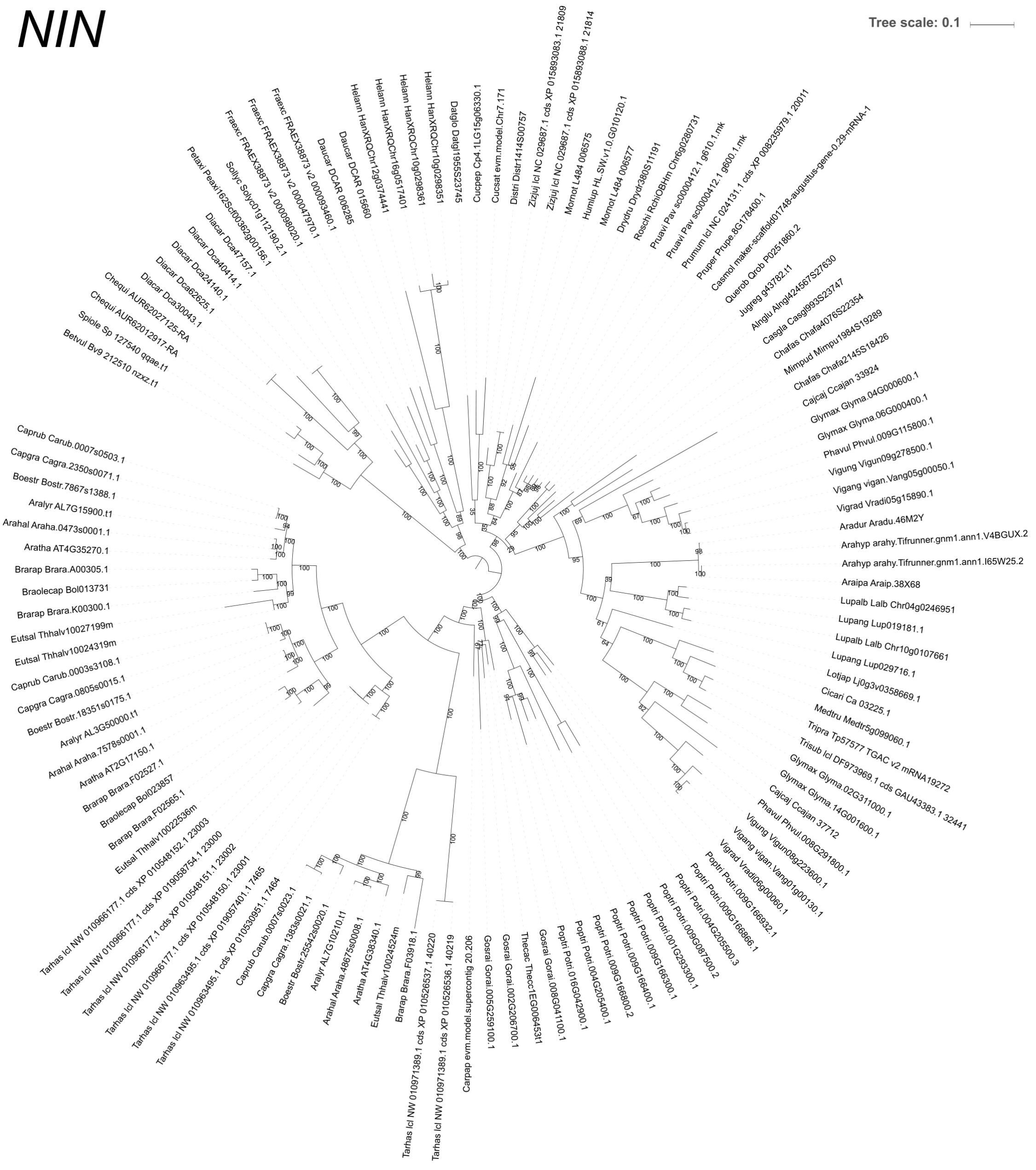

**Supplementary Figure 20. Phylogenetic tree of *NIN* homologs of 96 plant species, covering different Angiosperm orders and symbiotic abilities.**

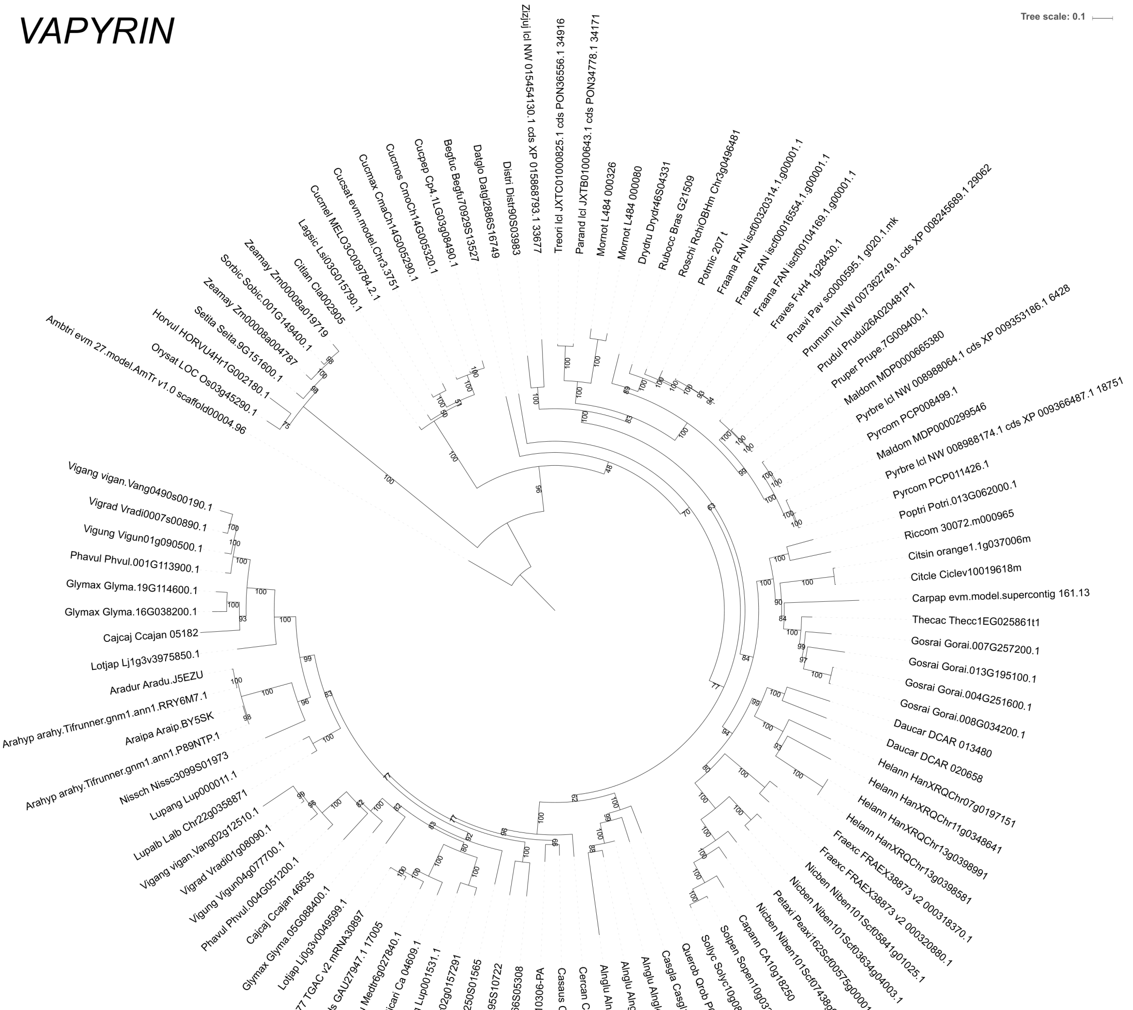

**Supplementary Figure 21. Phylogenetic tree of *VAPYRIN* homologs of 96 plant species, covering different Angiosperm orders and symbiotic abilities.**

*LYK3*

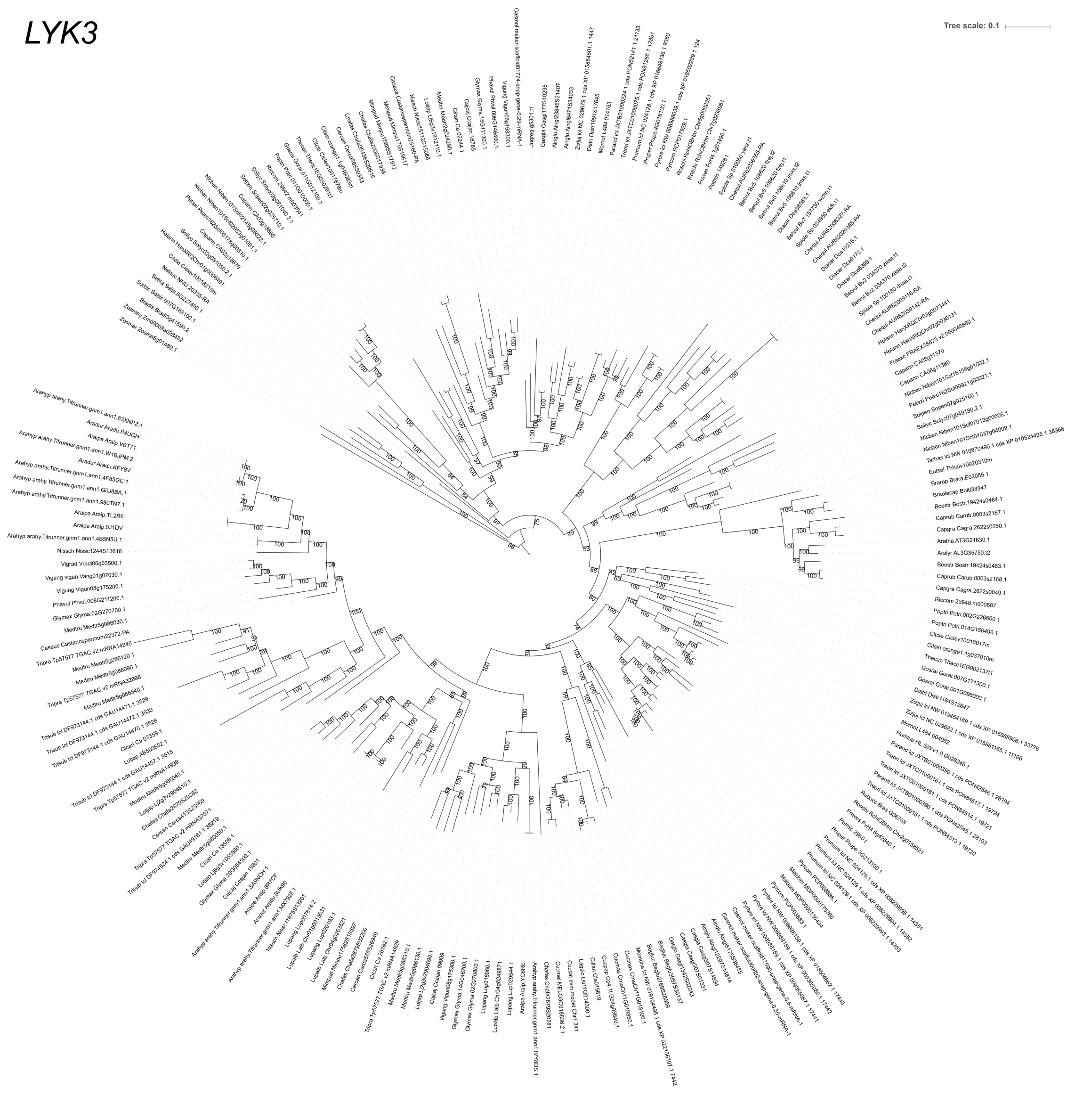

**Supplementary Figure 22. Phylogenetic tree of *LYK3* homologs of 96 plant species, covering different Angiosperm orders and symbiotic abilities.**

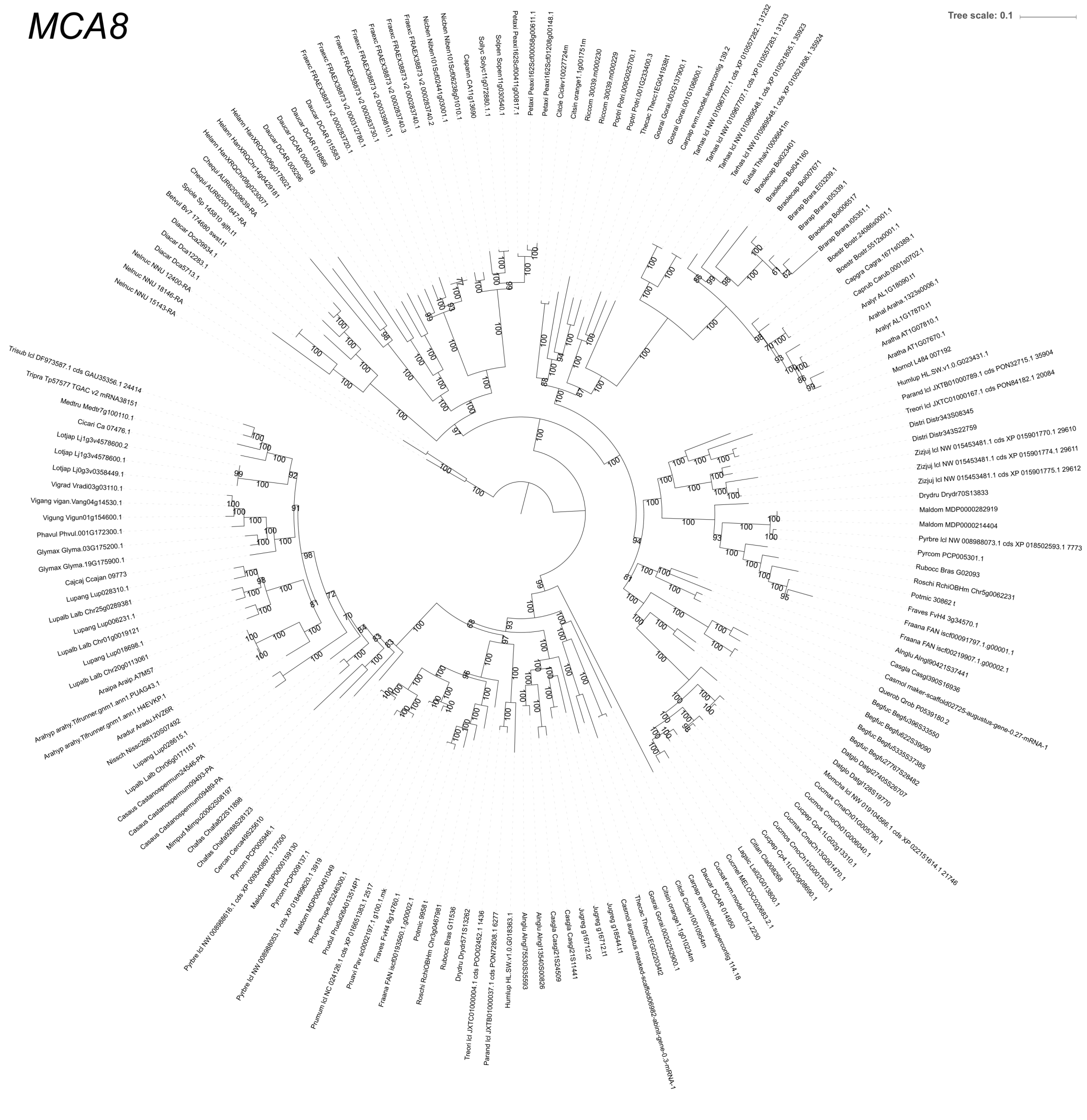

**Supplementary Figure 23. Phylogenetic tree of *MCA8* homologs of 96 plant species, covering different Angiosperm orders and symbiotic abilities.**

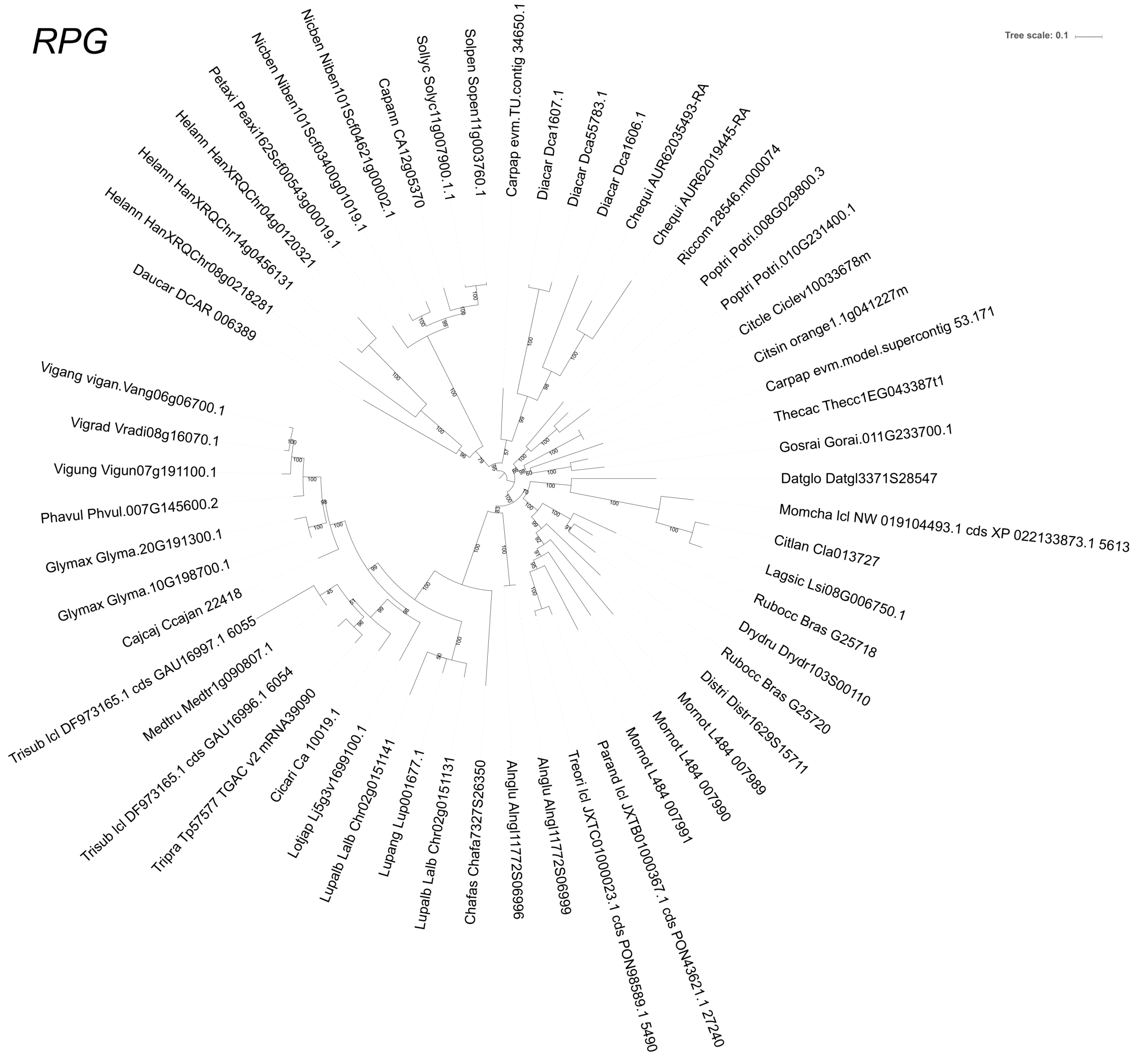

**Supplementary Figure 24. Phylogenetic tree of *RPG* homologs of 96 plant species, covering different Angiosperm orders and symbiotic abilities.**

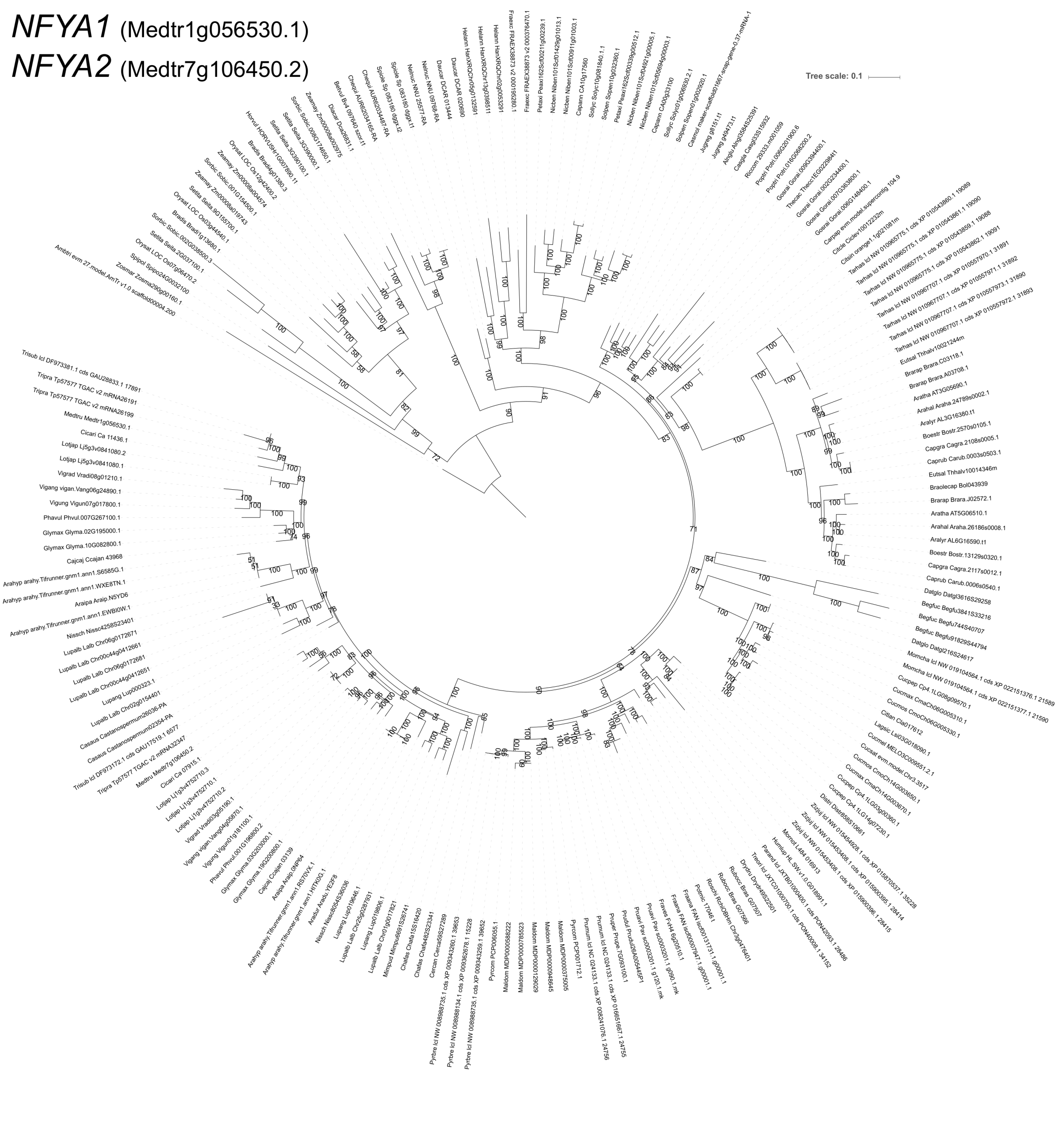

**Supplementary Figure 25. Phylogenetic tree of *MFYA1 and MFYA2* homologs of 96 plant species, covering different Angiosperm orders and symbiotic abilities.**

### *CRE1*

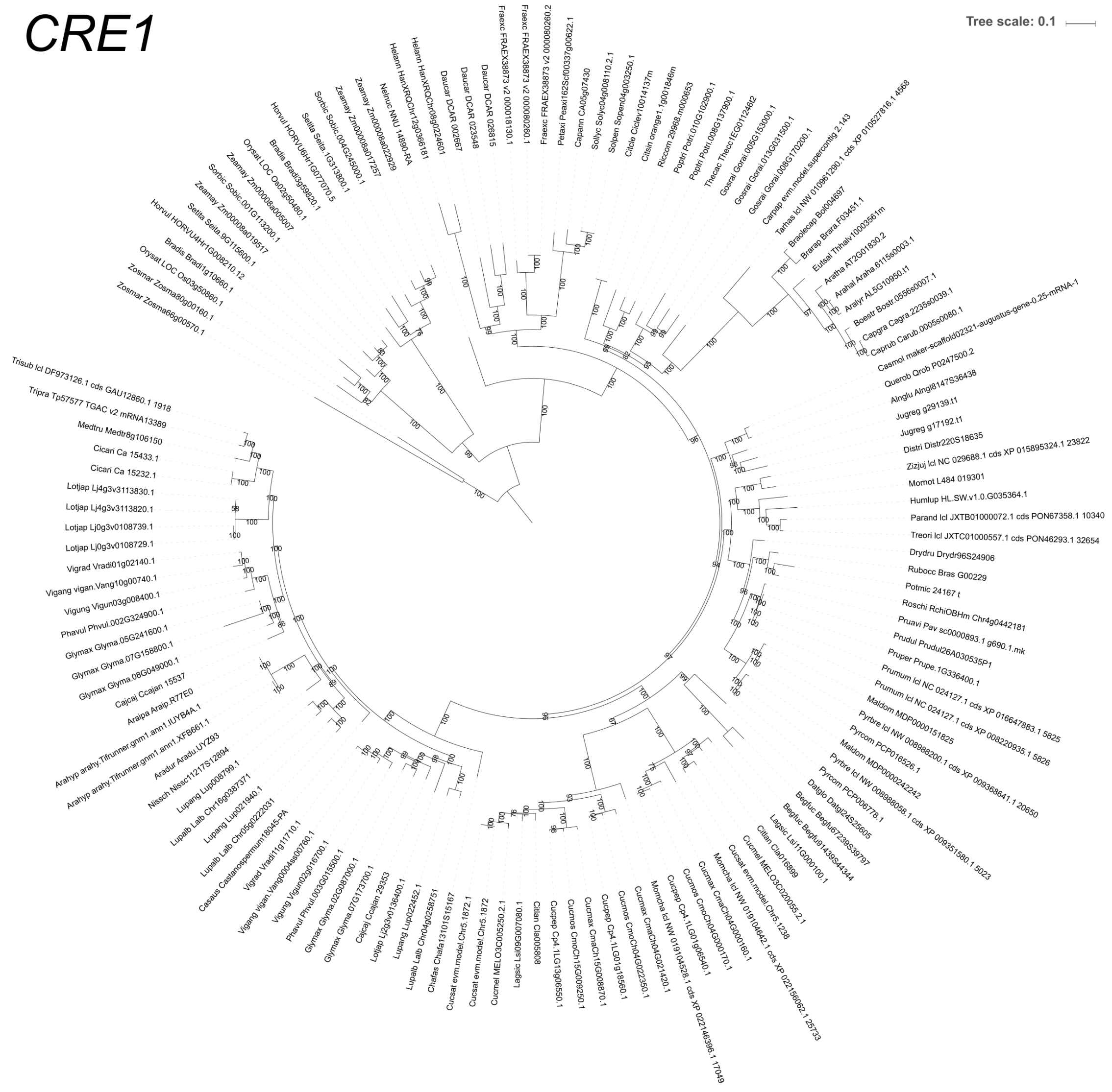

**Supplementary Figure 26. Phylogenetic tree of *CRE1* homologs of 96 plant species,  covering different Angiosperm orders and symbiotic abilities.**

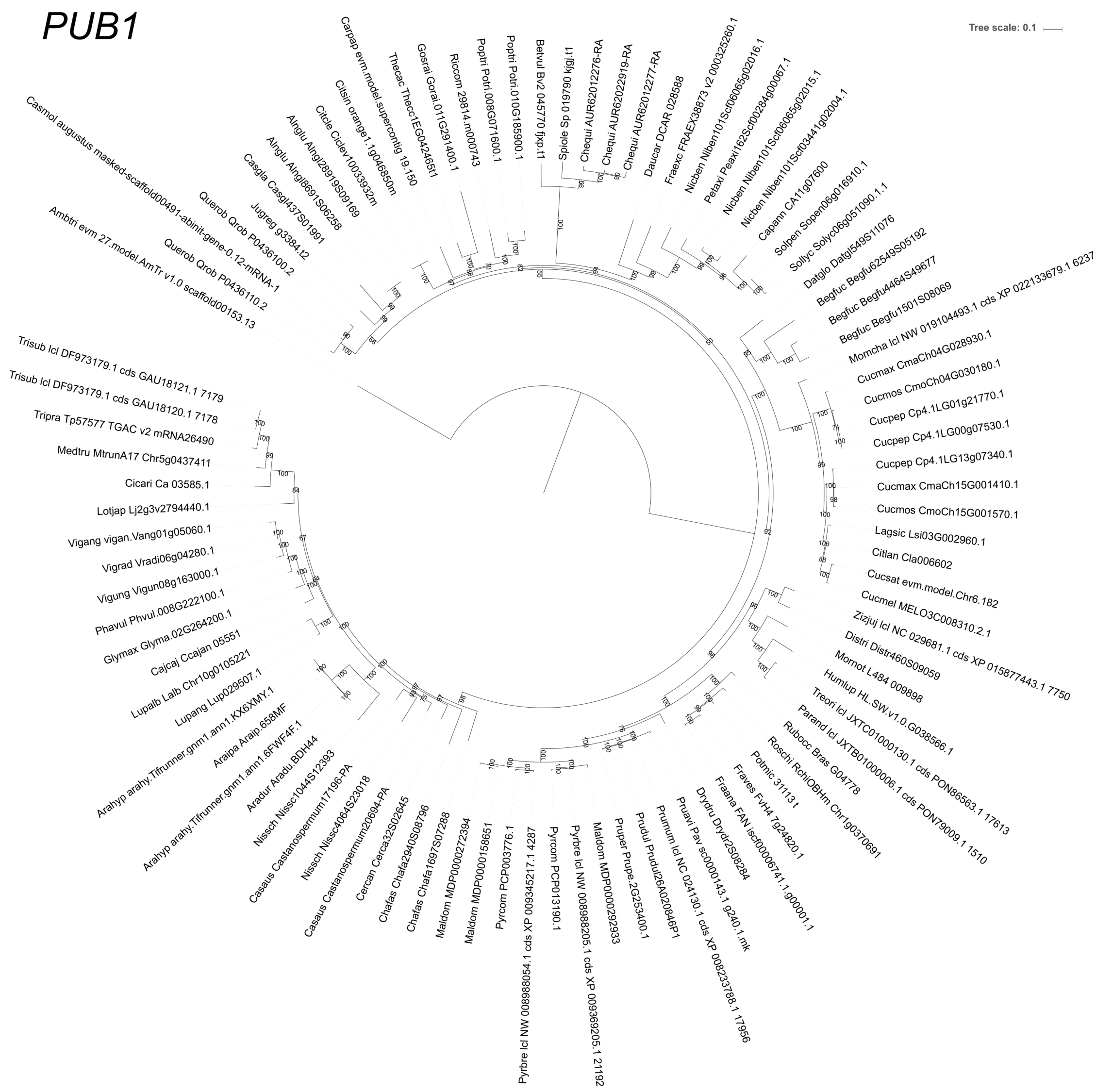

**Supplementary Figure 27. Phylogenetic tree of *PUB1* homologs of 96 plant species, covering different Angiosperm orders and symbiotic abilities.**

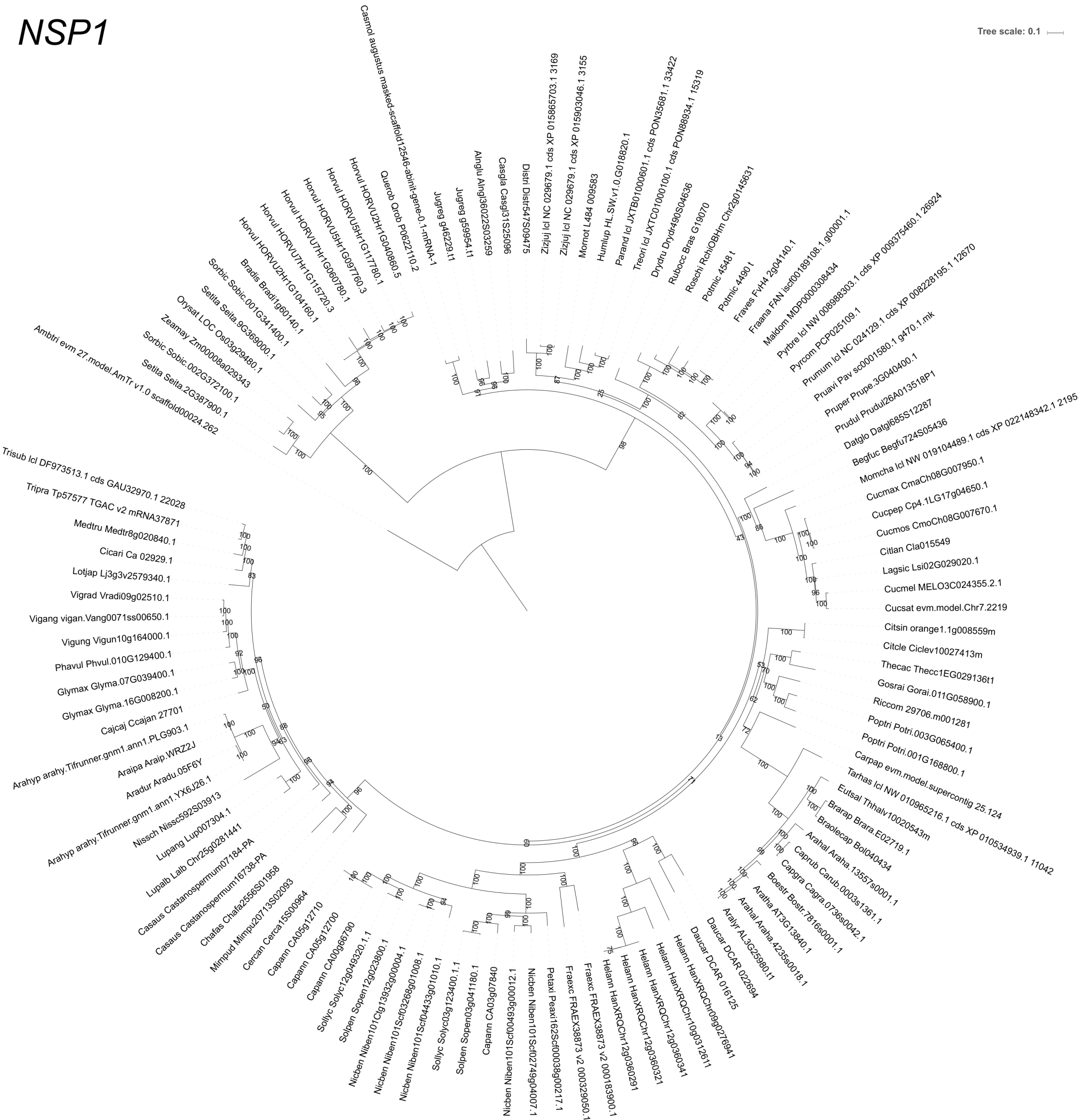

**Supplementary Figure 28. Phylogenetic tree of *NSP1* homologs of 96 plant species, covering different Angiosperm orders and symbiotic abilities.**

*ERN1* (Medtr7g085810.1)
*ERN2* (Medtr6g029180.1)

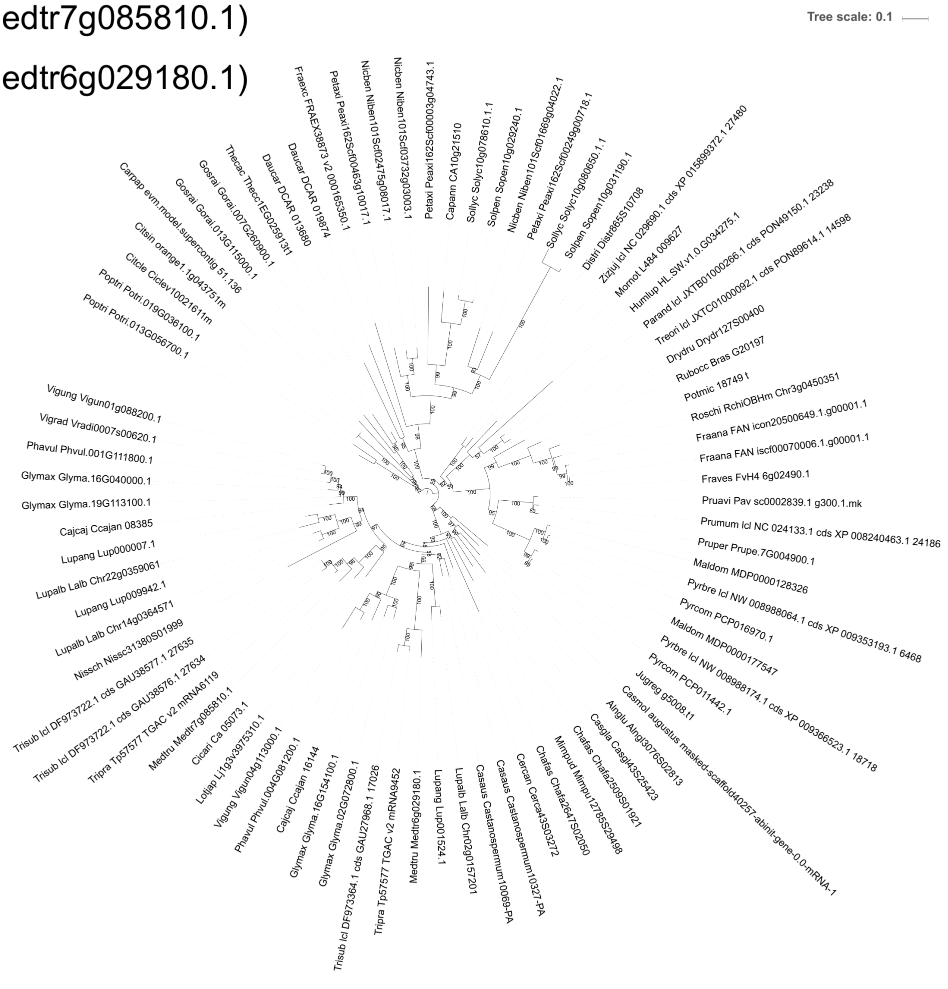

**Supplementary Figure 29. Phylogenetic tree of homologs of *ERN1* and *ERN2* 96 plant species, covering different Angiosperm orders and symbiotic abilities.**

*SUT1*

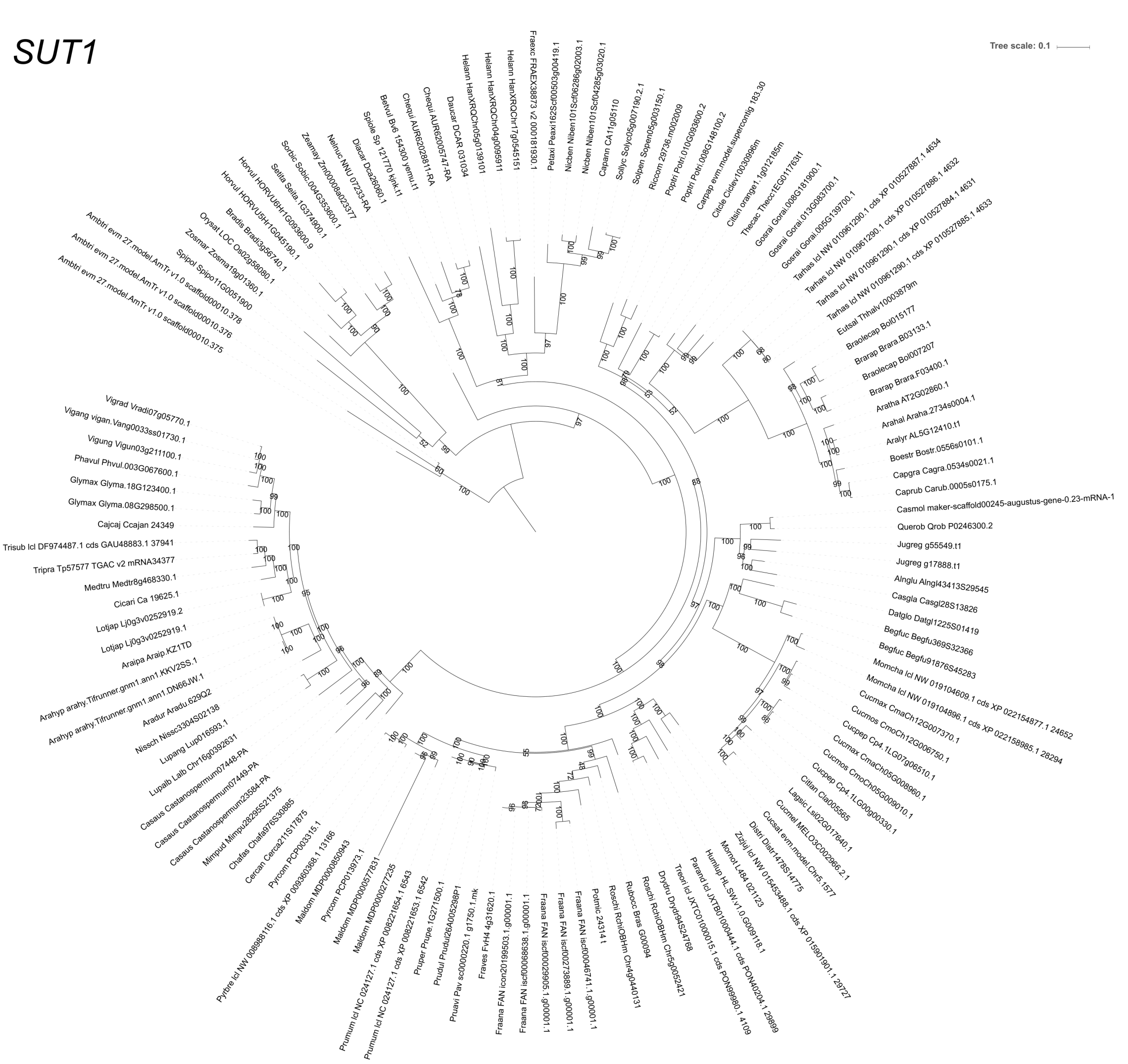

**Supplementary Figure 30. Phylogenetic tree of *SUT1* homologs of 96 plant species, covering different Angiosperm orders and symbiotic abilities.**

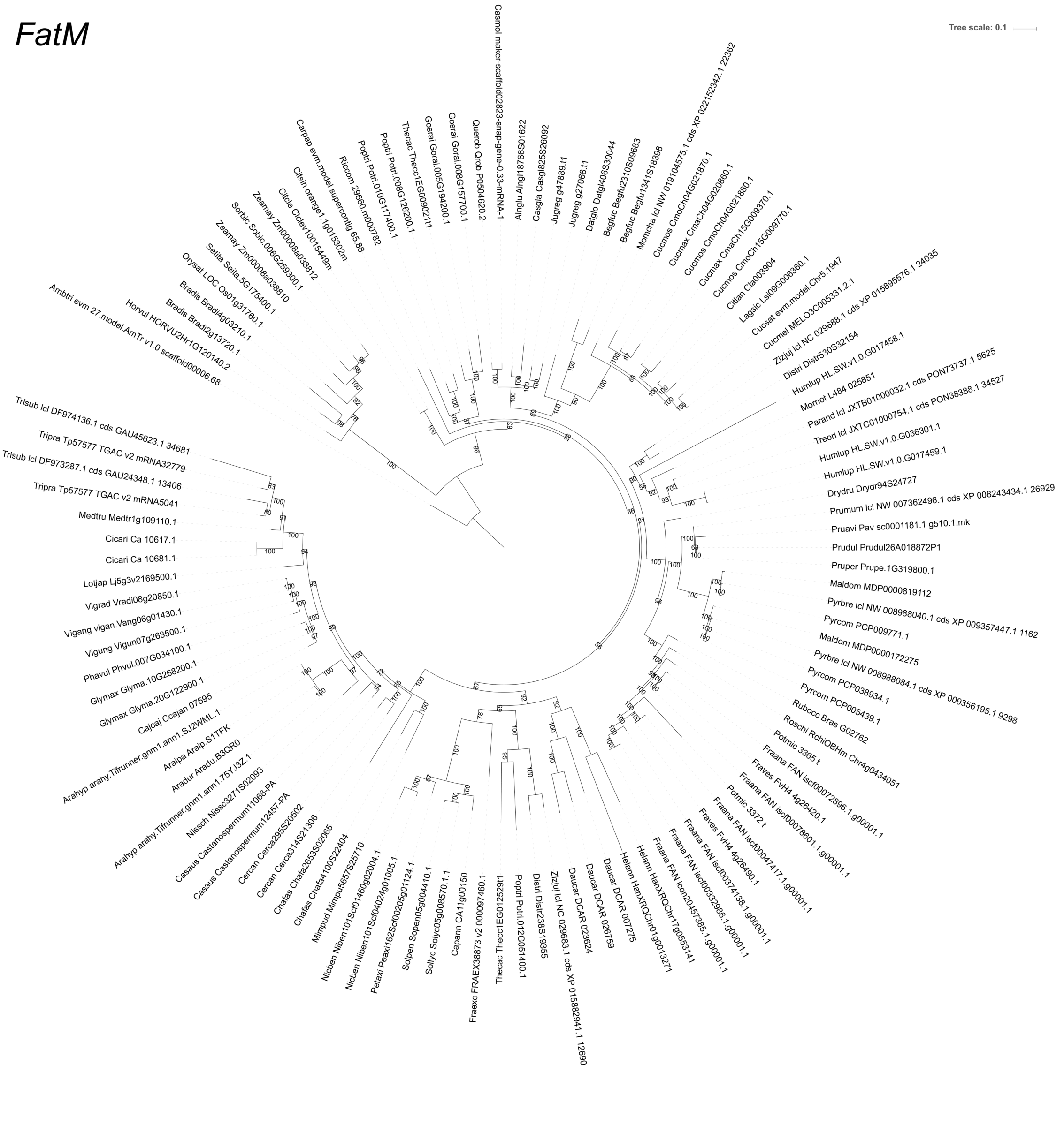

**Supplementary Figure 31. Phylogenetic tree of *FatM* homologs of 96 plant species, covering different Angiosperm orders and symbiotic abilities.**

*PT4*

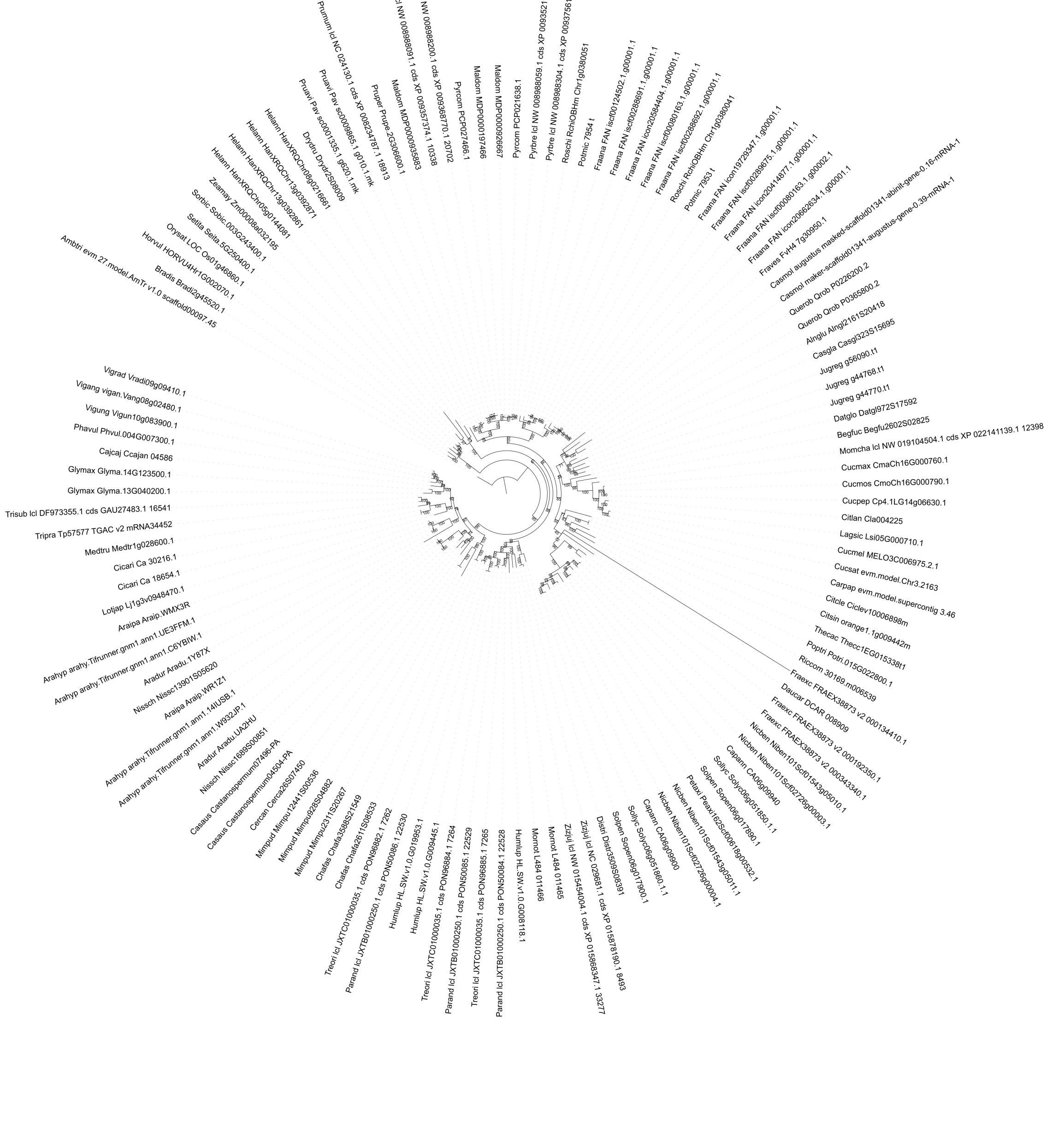

**Supplementary Figure 32. Phylogenetic tree of *PT4* homologs of 96 plant species, covering different Angiosperm orders and symbiotic abilities.**

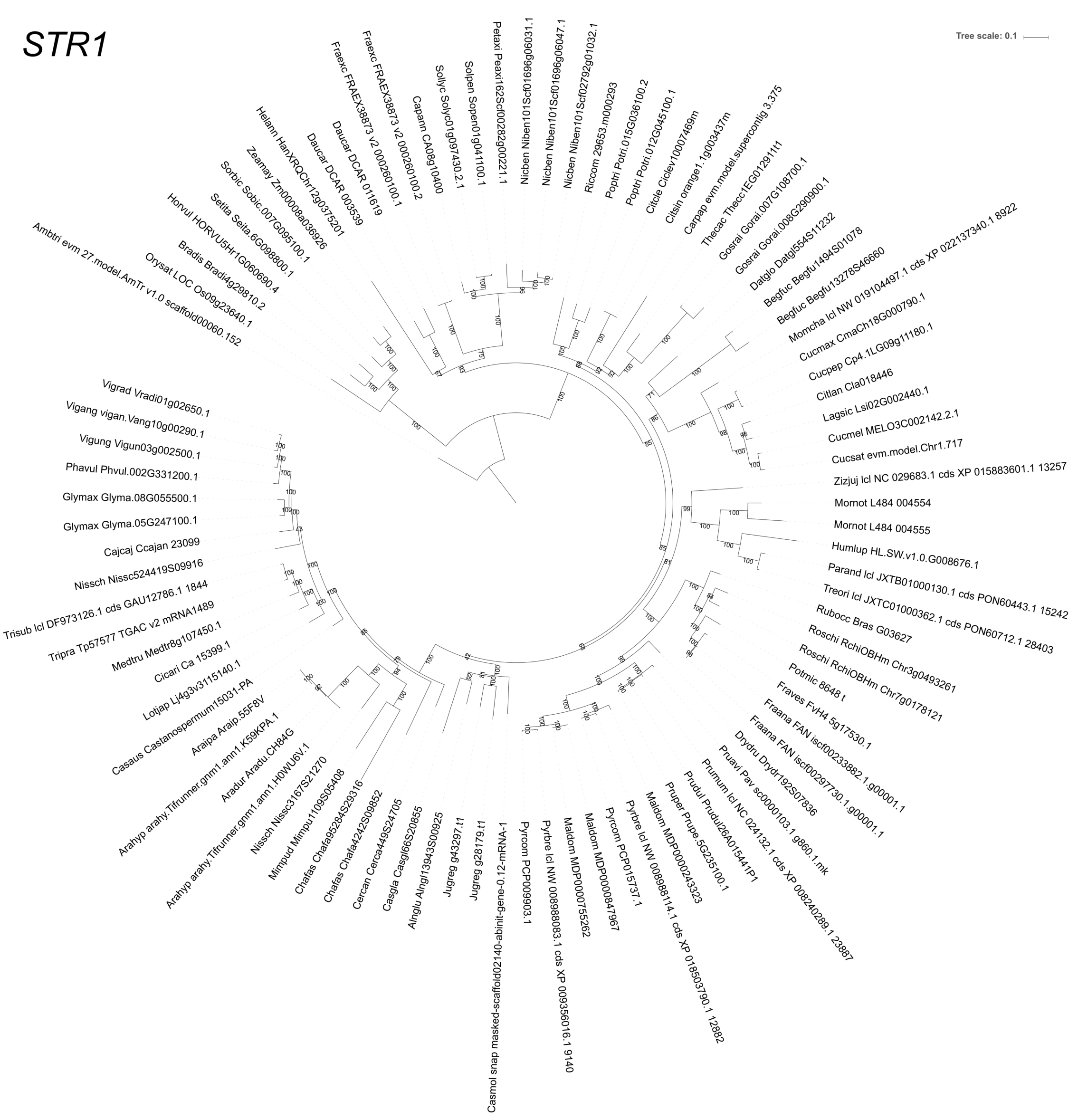

**Supplementary Figure 33. Phylogenetic tree of *STR1* homologs of 96 plant species, covering different Angiosperm orders and symbiotic abilities.**

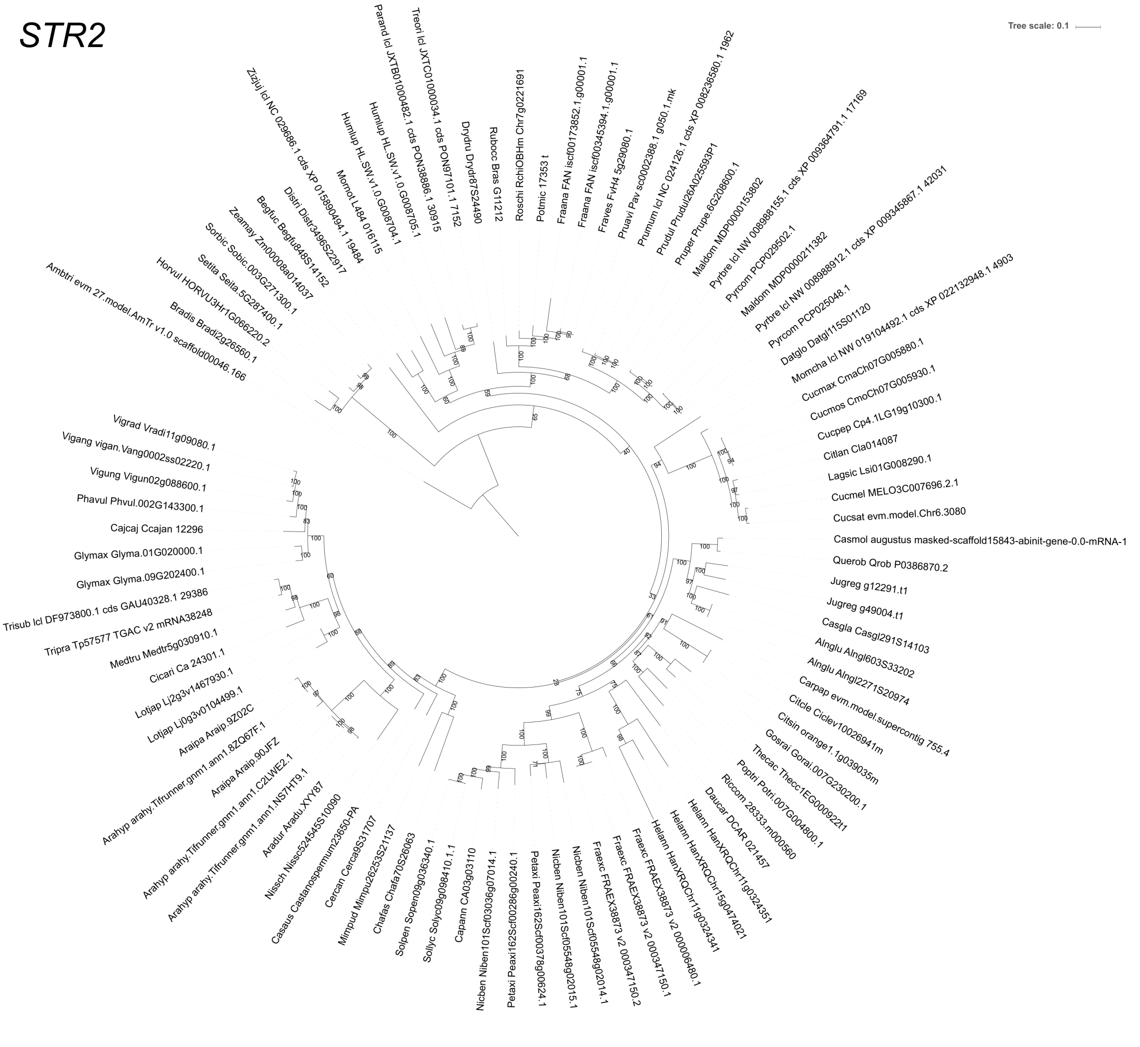

**Supplementary Figure 34. Phylogenetic tree of *STR2* homologs of 96 plant species, covering different Angiosperm orders and symbiotic abilities.**

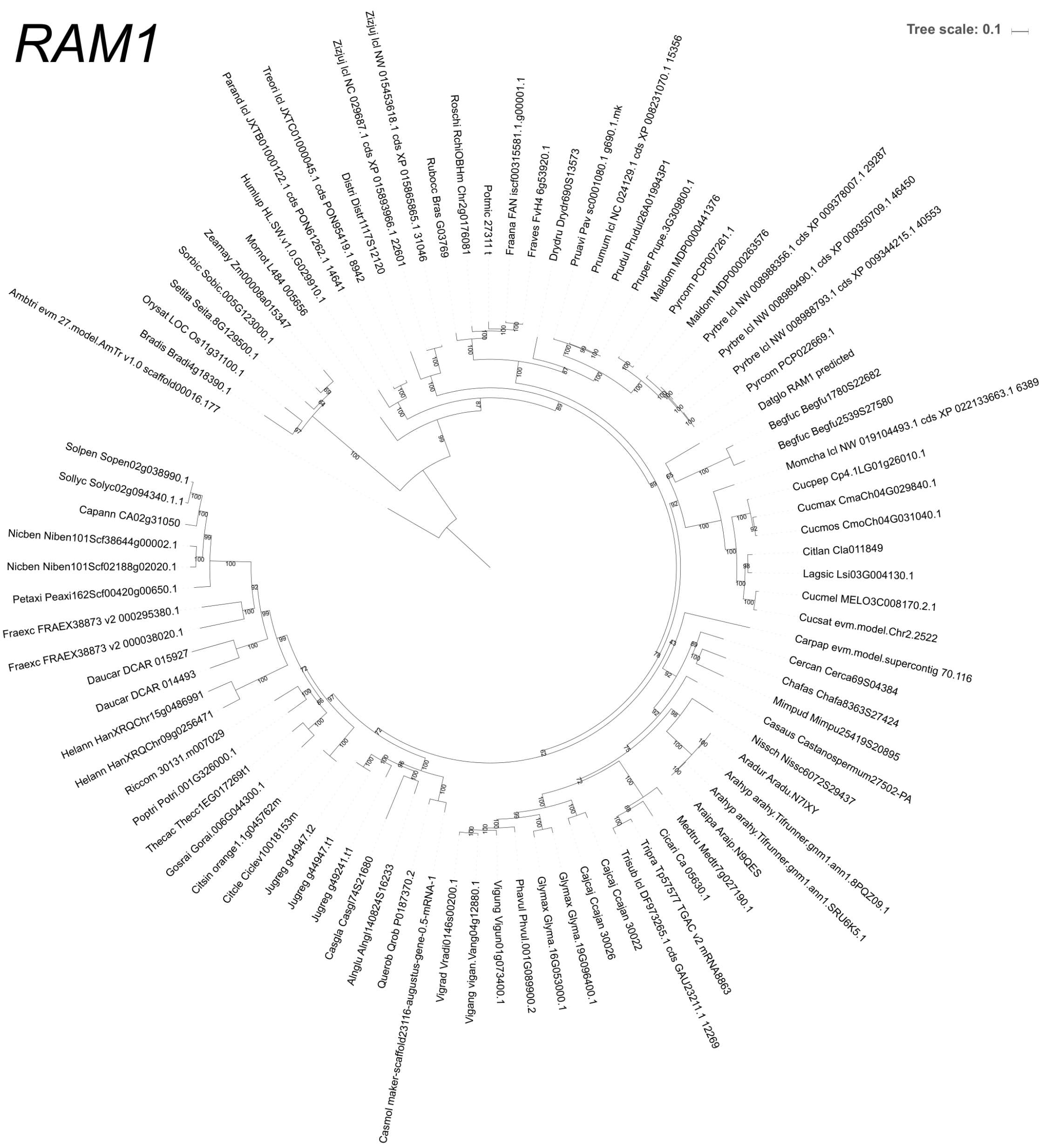

**Supplementary Figure 35. Phylogenetic tree of *RAM1* homologs of 96 plant species, covering different Angiosperm orders and symbiotic abilities.**

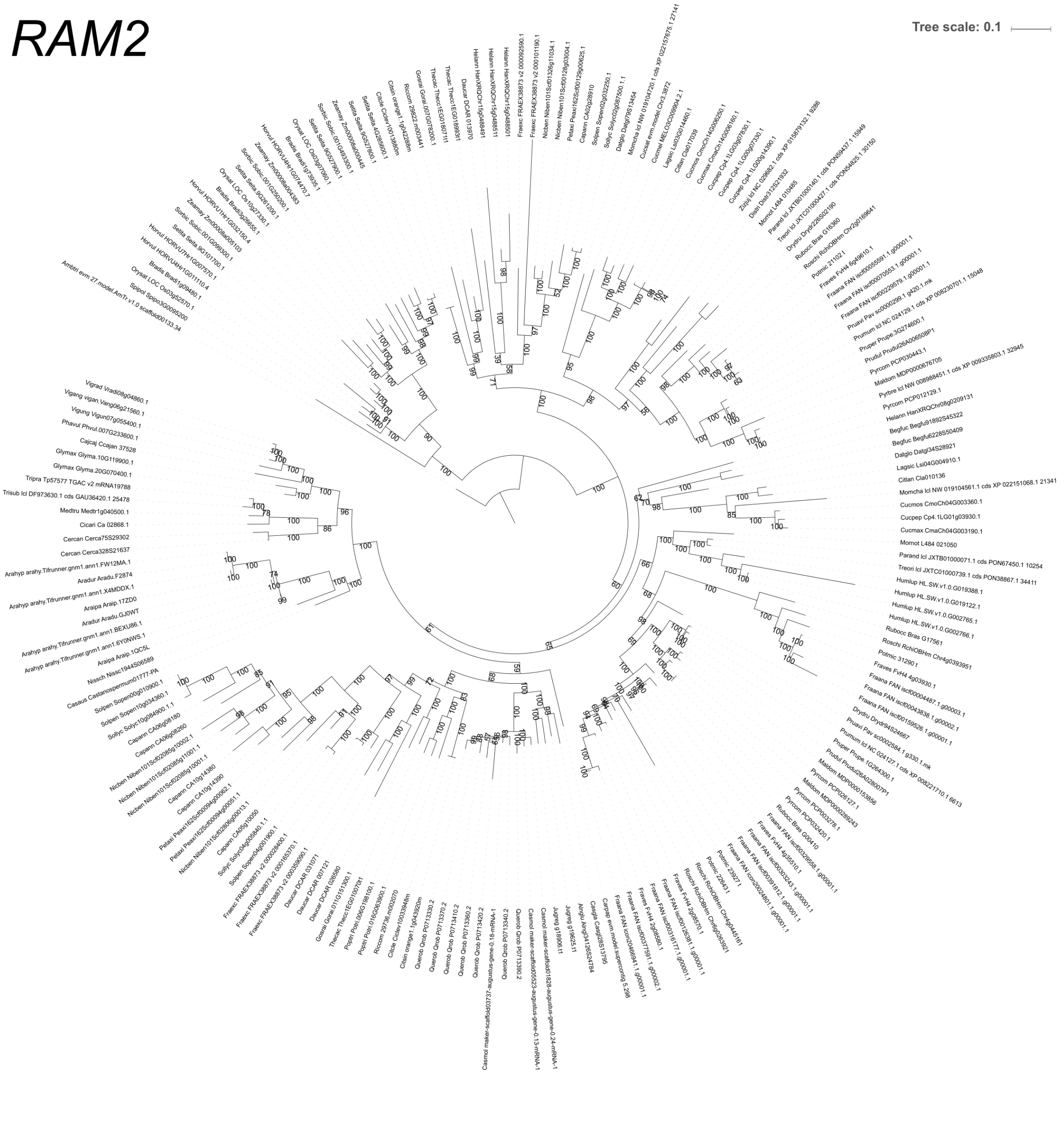

**Supplementary Figure 36. Phylogenetic tree of *RAM2* homologs of 96 plant species, covering different Angiosperm orders and symbiotic abilities.**

*HA1*

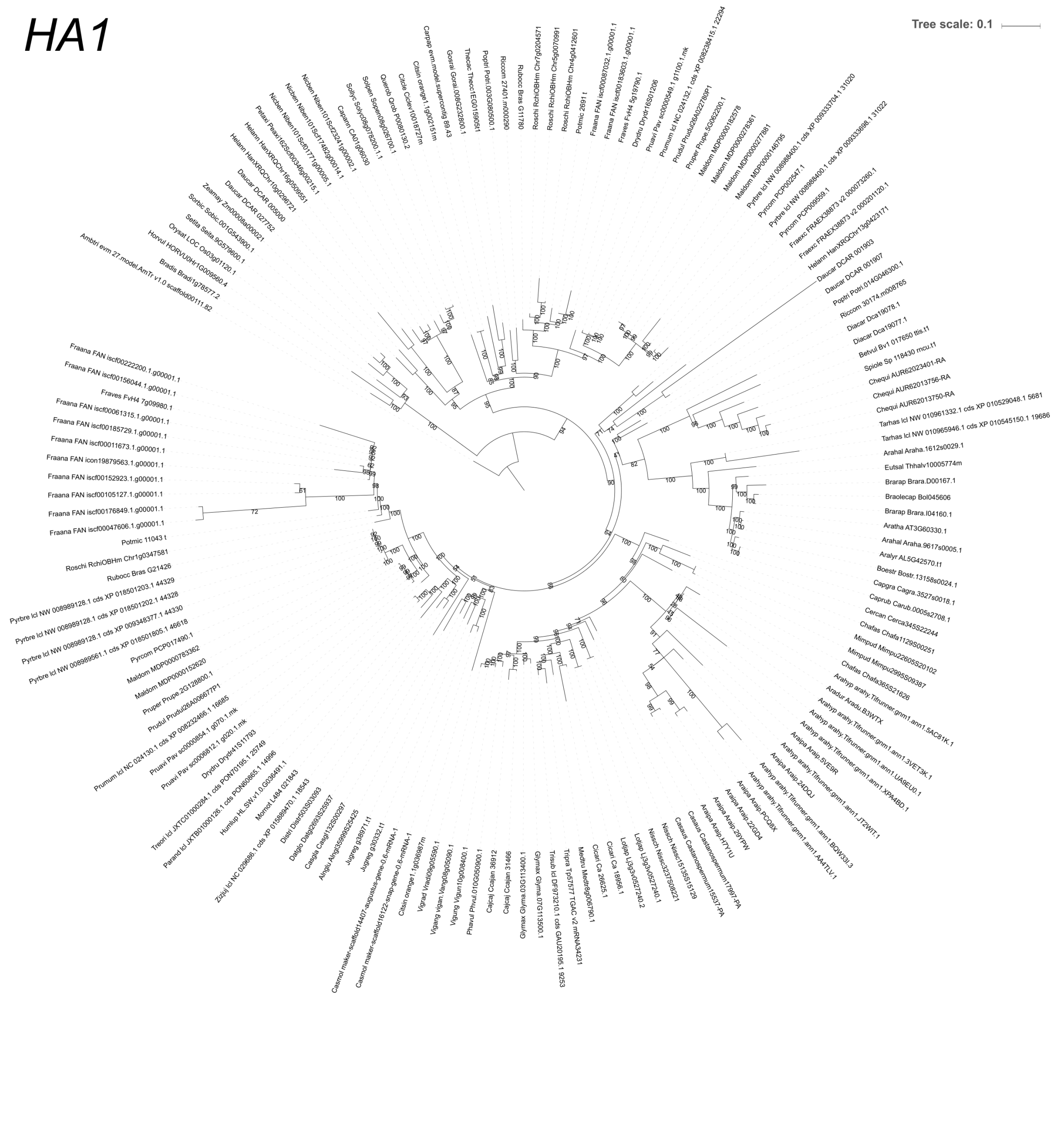

**Supplementary Figure 37. Phylogenetic tree of *HA1* homologs of 96 plant species, covering different Angiosperm orders and symbiotic abilities.**

### *Exo70*

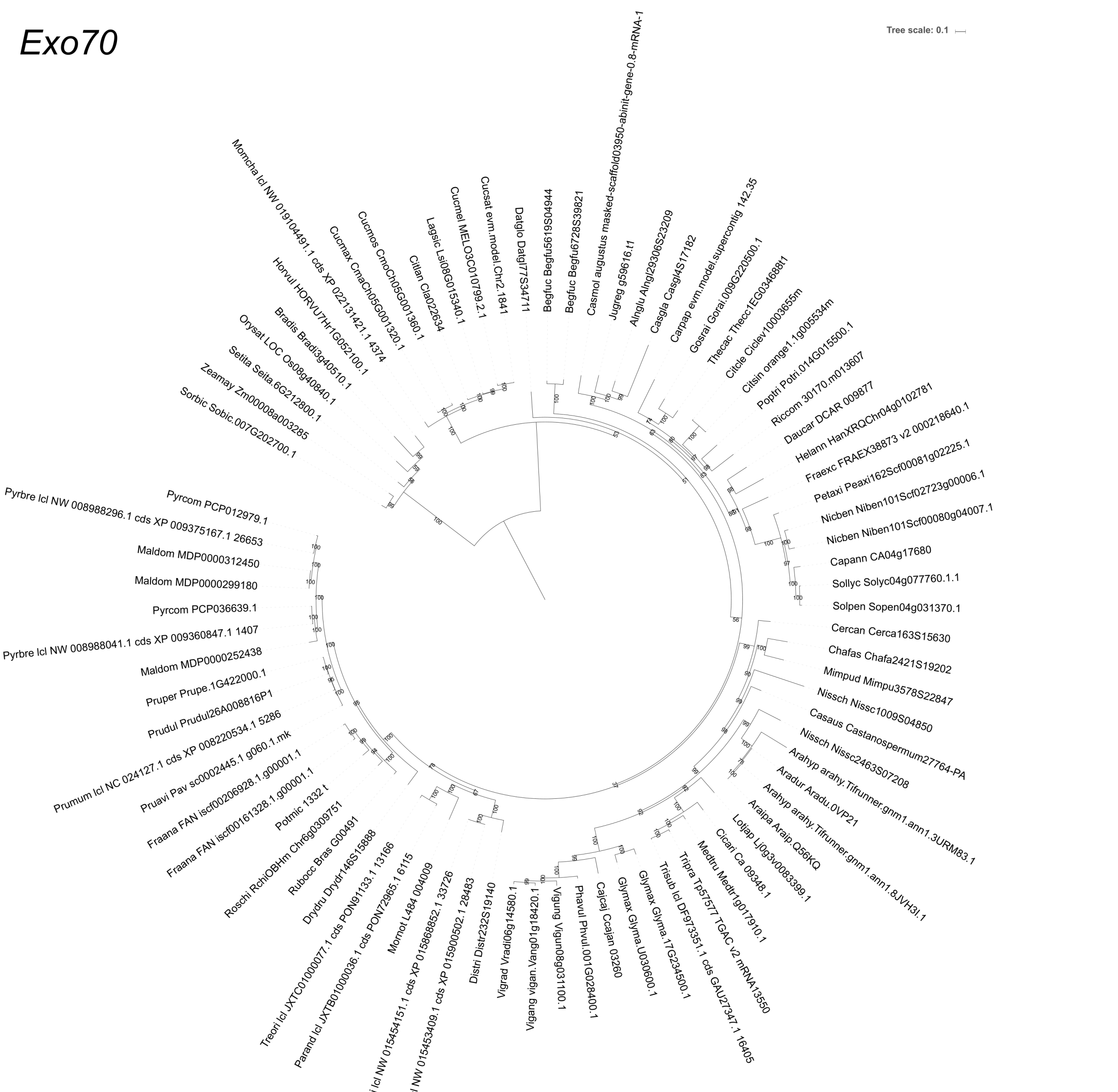

**Supplementary Figure 38. Phylogenetic tree of *Exo70* homologs of 96 plant species, covering different Angiosperm orders and symbiotic abilities.**

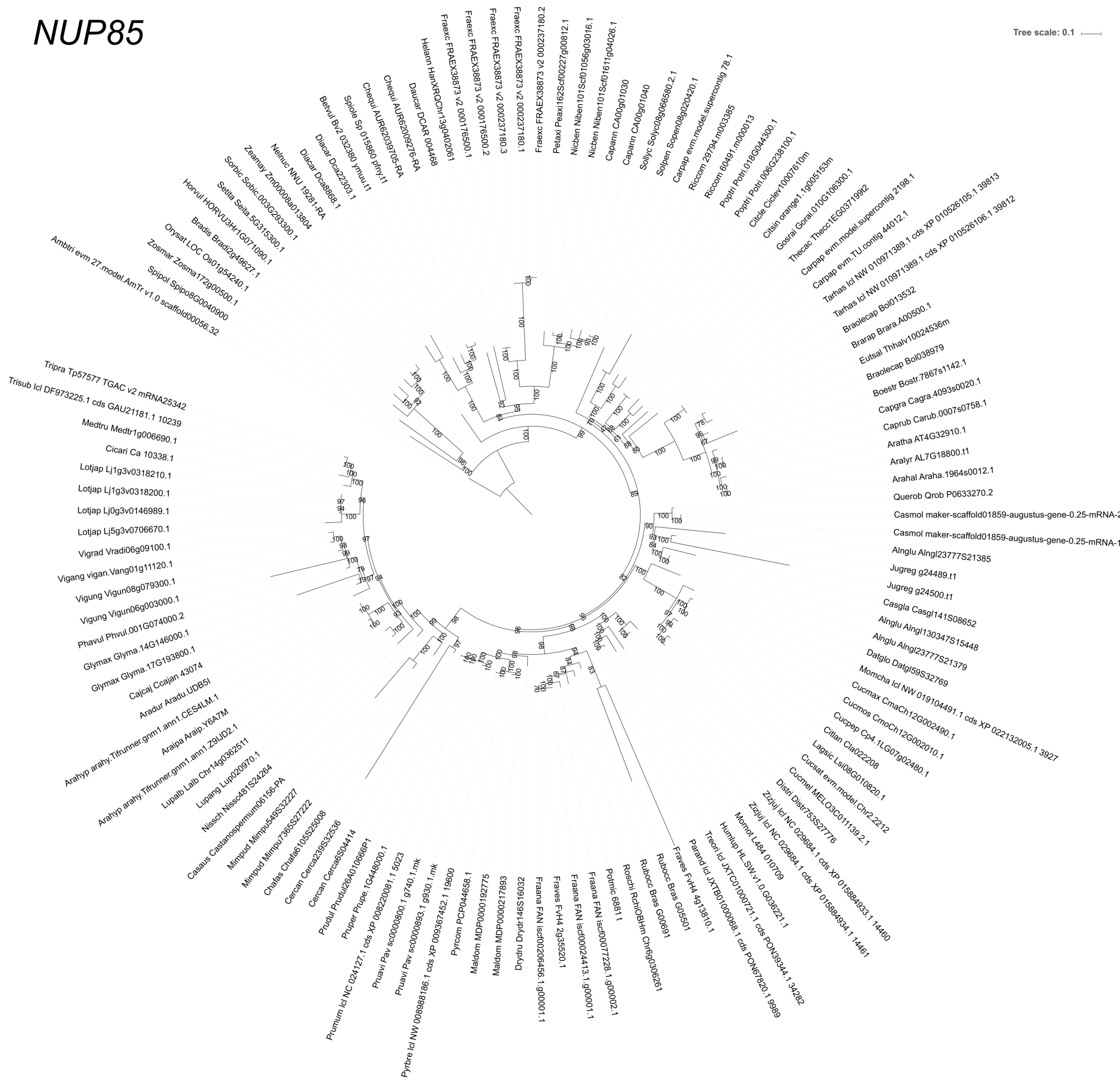

**Supplementary Figure 39. Phylogenetic tree of *NUP85* homologs of 96 plant species, covering different Angiosperm orders and symbiotic abilities.**

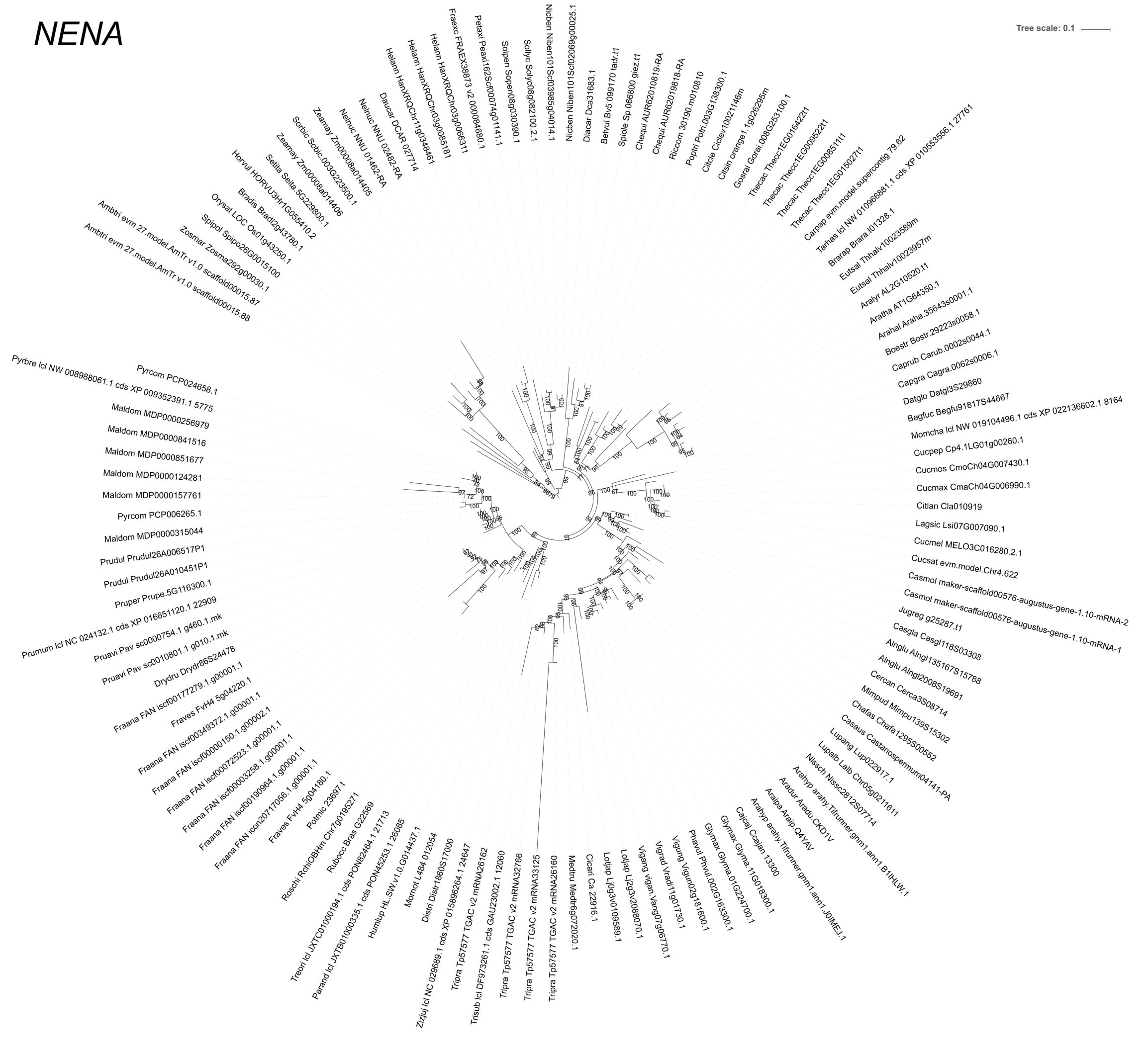

**Supplementary Figure 40. Phylogenetic tree of *NENA* homologs of 96 plant species, covering different Angiosperm orders and symbiotic abilities.**

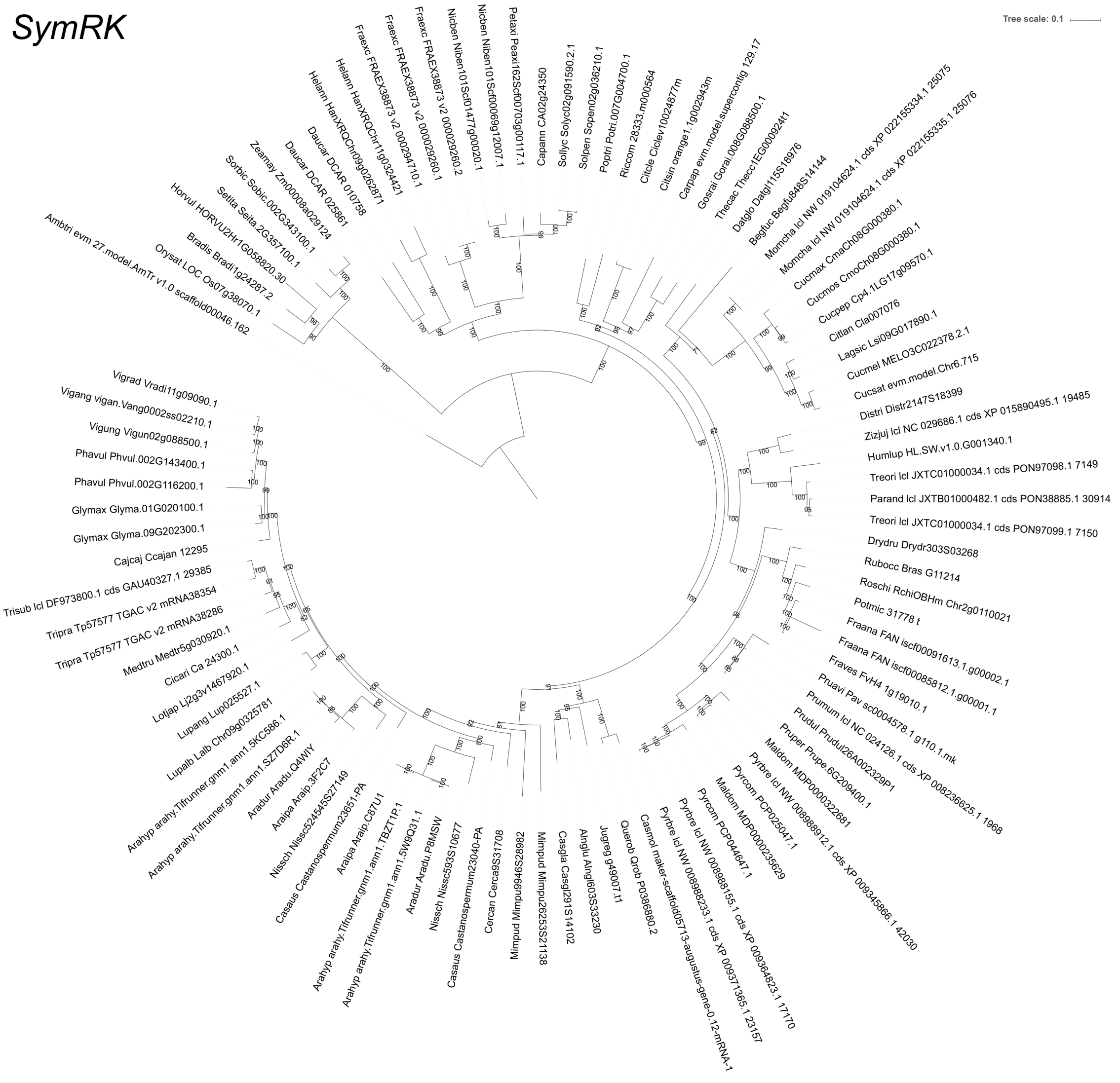

**Supplementary Figure 41. Phylogenetic tree of *SymRK* homologs of 96 plant species, covering different Angiosperm orders and symbiotic abilities.**

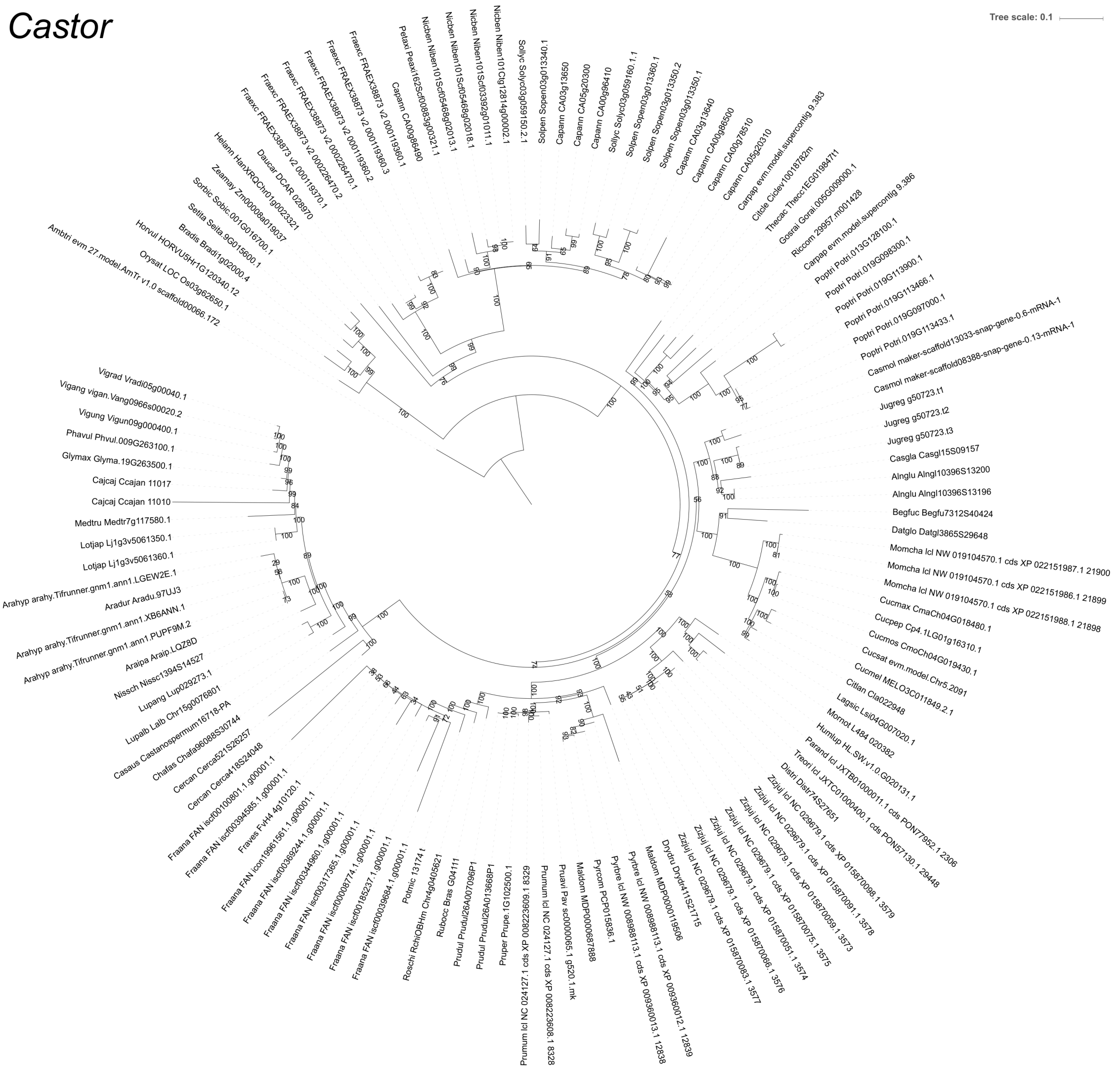

**Supplementary Figure 42. Phylogenetic tree of *Castor* homologs of 96 plant species, covering different Angiosperm orders and symbiotic abilities.**
